# Age-Associated Insolubility of Parkin in Human Midbrain is Linked to Redox Balance and Sequestration of Reactive Dopamine Metabolites

**DOI:** 10.1101/2020.11.20.392175

**Authors:** Jacqueline M. Tokarew, Daniel N. El-Kodsi, Nathalie A. Lengacher, Travis K. Fehr, Angela P. Nguyen, Bojan Shutinoski, Brian O’Nuallain, Ming Jin, Jasmine M. Khan, Andy C. H. Ng, Juan Li, Qiubo Jiang, Mei Zhang, Liqun Wang, Rajib Sengupta, Kathryn R. Barber, An Tran, Stephanie Zandee, Xiajun Dong, Clemens R. Scherzer, Alexandre Prat, Eve Tsai, Masashi Takanashi, Nobutaka Hattori, Jennifer A. Chan, Luigi Zecca, Andrew B. West, Arne Holmgren, Lawrence Puente, Gary S. Shaw, Gergely Toth, John M. Woulfe, Peggy Taylor, Julianna J. Tomlinson, Michael G. Schlossmacher

**Affiliations:** Program in Neuroscience, Ottawa Hospital Research Institute, Ottawa, ON, Canada; Graduate Program in Cellular and Molecular Medicine (Neuroscience), Faculty of Medicine, University of Ottawa, Ottawa, ON, Canada; BioLegend Inc., Dedham, MA., USA; Department of Pathology and Laboratory Medicine, The Ottawa Hospital, Ottawa, ON, Canada; Department of Biochemistry, Karolinska Institute, Stockholm, Sweden; Present Address: Amity Institute of Biotechnology, Amity University, Kolkata, West Bengal 700135, India; Department of Biochemistry, University of Western Ontario, London, ON, Canada; Department of Neuroscience, Faculty of Medicine, University of Montreal, Montreal, QC, Canada; Ann Romney Center for Neurologic Diseases, Brigham & Women’s Hospital, Boston, MA, USA; Division of Neurosurgery, Department of Surgery, The Ottawa Hospital, Ottawa, ON, Canada; Department of Neurology, Juntendo University School of Medicine, Tokyo, Japan; Department of Pathology & Laboratory Medicine, University of Calgary, Calgary, AB, Canada; Institute of Biomedical Technologies, Italian National Research Council, Segrate (Milano), Italy; Departments of Neurobiology and Pharmacology & Cancer Biology, Duke University, Durham, NC, USA; Proteomics Core Facility, Ottawa Hospital Research Institute, Ottawa, ON, Canada; Institute of Organic Chemistry, Research Center for Natural Sciences, Budapest, Hungary; University of Ottawa Brain and Mind Research Institute, Ottawa, ON, Canada; Division of Neurology, Department of Medicine, The Ottawa Hospital, Ottawa, ON, Canada

**Author notes:** these authors contributed equally to this work (Jacqueline M. Tokarew, Daniel N. El-Kodsi). deceased (Arne Holmgren). Lead contacts (Julianna J. Tomlinson, Michael G. Schlossmacher).

**Keywords:** Early-Onset Parkinson disease, Parkin, *PRKN* Gene, Redox Chemistry, Dopamine Metabolism, Neuromelanin

## Abstract

The mechanisms by which parkin protects the adult human brain from Parkinson disease remain incompletely understood. We hypothesized that parkin cysteines participate in redox reactions, which are reflected in its posttranslational modifications. We found that in human control brain, including the *S. nigra*, parkin is largely insoluble after age 40 years, which is linked to its oxidation, *e.g.,* at Cys95 and Cys253. In mice, oxidative stress increases posttranslational modifications at parkin cysteines and reduces its solubility. Oxidation of recombinant parkin also promotes insolubility and aggregate formation, but in parallel, lowers hydrogen peroxide (H_2_O_2_). This thiol-based redox activity is diminished by parkin point mutants, *e.g.,* p.C431F and p.G328E. Intriguingly, in parkin-deficient human brain H_2_O_2_ concentrations are elevated. In *prkn-*null mice, H_2_O_2_ levels are dysregulated under oxidative stress conditions, such as acutely by MPTP-toxin exposure or chronically due to a second genetic hit. In dopamine toxicity studies, wild-type parkin, but not disease-linked mutants, protects human dopaminergic M17 cells, in part through lowering H_2_O_2_. Parkin also neutralizes reactive, electrophilic dopamine metabolites via adduct formation, which occurs foremost at primate-specific Cys95. Further, wild-type but not p.C95A-mutant parkin augments melanin formation. In sections of normal, adult human midbrain, parkin specifically co-localizes with neuromelanin pigment, frequently within LAMP-3/CD63^+^ lysosomes. We conclude that oxidative modifications of parkin cysteines are associated with protective outcomes, which include the reduction of H_2_O_2_, conjugation of reactive dopamine metabolites, sequestration of radicals within insoluble aggregates, and increased melanin formation. The loss of these redox effects may augment oxidative stress in dopamine producing neurons of mutant *PRKN* allele carriers, thereby contributing to neurodegeneration.

## Introduction

Bi-allelic mutations in *PRKN*, which encodes parkin, lead to a young-onset, recessive form of Parkinson disease (PD)[1, 2]. Pathology studies of parkin-deficient brains have demonstrated that neuronal loss is largely restricted to the *S. nigra* and *L. coeruleus*, two brainstem nuclei that synthesize dopamine (reviewed in Doherty *et al.*[3]).

Parkin is a principally cytosolic protein. It has been associated with diverse cellular functions, foremost related to its ubiquitin ligase (E3) activity, the control of inflammation signaling, and maintenance of mitochondrial integrity, as mediated through participation in mitophagy and mitochondrial antigen presentation (MITAP)[4–11] (reviewed in Barodia *et al.* [12]). Although mitophagy has recently been shown to be co-regulated by parkin in the developing heart of mice [13], the diverse roles ascribed to parkin function have not yet explained its selective neuroprotection. For example, vertebrate models of genomic *prkn* deletion do not reproduce dopamine cell loss; one exception is the parkin-deficient *Polg* mouse, where mitochondrial DNA mutagenic stress had been added as a second, genetic hit [14]. The general lack of dopamine cell loss in genomic parkin deficiency-based models could be due to compensatory mechanisms [15], a shorter life span of non-human mammals, and possibly, unique aspects of dopamine’s breakdown in humans. The latter is exemplified by the generation of cytoplasmic neuromelanin in dopamine synthesizing neurons beginning after childhood [16]. Nevertheless, genomic *prkn*-null models have revealed biochemical and structural changes in high energy-producing cells of flies and murine tissues [12, 17, 18], which suggested the presence of elevated oxidative stress [19–21]. These observations pointed at a contribution of parkin to redox homeostasis *in vivo.*

Redox equilibrium invariably involves cysteine-based chemistry. There, thiols are subject to oxidative modifications by reactive oxygen-, reactive nitrogen- and reactive electrophilic species (ROS, RNS, RES) [22, 23], some of which are reversible. Proteins irreversibly conjugated by RES, including by electrophilic dopamine radicals, are either degraded or sequestered within inclusions. It is thought that the latter process occurs via lysosomal functions and underlies neuromelanin formation throughout adulthood [24].

Human parkin contains 35 cysteines (Cys; single letter code, C) [1], its murine homologue 34. Of these, 28 cysteines are involved in the chelation of eight zinc ions within four RING domains [25]. Although Cys431 has been identified as critical in catalyzing parkin’s E3 ligase function, 6 other cysteines are structurally unaccounted for, including Cys95 located within parkin’s ‘linker’ domain. Several reports have demonstrated unique sensitivity of parkin to ROS and RES in cells [26–28]. Further, RNS and sulfhydration also alter its cysteines residues, and NO-/NO_2_-modified parkin variants have been described in cells and brain tissue [29–33]. Oxidation of parkin has been linked to both activating (‘gain-of-function’) and detrimental (‘loss-of-function’) outcomes when tested in the context of parkin’s E3 ligase activity *in vitro* [27, 29, 31, 34].

We found that wild-type parkin is highly oxidized and insoluble in adult human midbrain, leading us to explore non-E3 ligase-mediated protective functions, as informed by its metabolism. Owing to its number of cysteine-based thiols, we hypothesized that parkin could confer neuroprotection by acting as an anti-oxidant molecule and that it contributes to redox balance *in vivo* by reducing ROS/RNS levels and conjugating dopamine radicals (RES). We posit that selective neurodegeneration in *PRKN*-linked, autosomal-recessive PD (ARPD) could be partially explained by the absence of parkin-mediated sequestration of toxic metabolites during decades of human ageing.

## Materials and methods

### Tissue collection

All tissues were collected in accordance with Institutional Review Board-approved guidelines. Fresh frozen samples of cortical human brain from subjects under 50 years of age were acquired through the University of Alabama and the Autism Tissue Program. *Post mortem*, frozen brain samples from frontal cortices were also obtained from the NICHD Brain and Tissue Bank at the University of Maryland. Brain tissues, including midbrain specimens, with short PMI were also obtained from patients diagnosed with clinical and neuropathological multiple sclerosis (MS) according to the revised 2010 McDonald’s criteria (n=4) [35]. Tissue samples were collected from MS patients with full ethical approval and informed consent as approved by the Montreal-based CRCHUM research ethics committee. Autopsy samples were preserved and lesions classified using Luxol Fast Blue / Haematoxylin & Eosin staining and Oil Red-O staining as previously published [36, 37]. No inflamed tissue areas were used in this current study.

Additional, fresh-frozen and paraffin-embedded human samples were obtained from the Neuropathology Service at Brigham and Women’s Hospital in Boston, MA. and from archived autopsy specimens in the Department of Pathology and Laboratory Medicine of The Ottawa Hospital, Ottawa, ON. Human spinal cord and muscle tissues were collected *post mortem* from organ donors at The Ottawa Hospital with approval from the Ottawa Health Science Network Research Ethics Board.

### Mouse tissues

Brains and hearts were collected from wild-type C57Bl/6J from Jackson laboratories, *prkn*-null from Dr. Brice’s laboratory [21], *Sod2* +/− mice from Jackson laboratories; the bi-genic mouse (*prkn*^−/−^//*Sod2*^+/−^) was created by crossing *prkn*-null mice with *Sod2* haploinsufficient mice, and interbreeding heterozygous offspring. These bi-genic mice have been characterized elsewhere (El Kodsi et al., in preparation [38]). Mouse brains collected were homogenized on ice in a Dounce glass homogenizer by 20 passes in Tris salt buffer with or without the addition of 1% H_2_O_2_, transferred to ultracentrifuge tubes and spun at 55,000 and 4°C for 30 mins to extract the soluble fraction. The resulting pellets were further homogenized in the tris salt buffer with the addition of 2-10% SDS, transferred to ultracentrifuge tubes and spun at 55,000 rpm and 10°C for 30 minutes to extract the insoluble fraction.

Wild-type C57Bl/6J mice were used for analysis of the effects of *post mortem* interval on murine parkin in the brain. Mice ranging from 4 to 8 months in age were perfused with PBS and their brains were collected for *post mortem* interval experiments.

### Sequential extraction of parkin from tissue

Roughly 1 cm^3^ samples of human brain frontal cortex and midbrain (age range 5-85 years of age) were weighed and placed in 3X volume/weight of Tris Salt buffer (TSS) (5mM Tris, 140 mM NaCl pH 7.5) containing complete EDTA-free protease inhibitor cocktail, and 10 mM iodoacetamide (IAA). The samples were homogenized on ice in a Dounce glass homogenizer by 50 passes, transferred to ultracentrifuge tubes and spun at 55,000 rpm and 4°C for 30 mins. The TS supernatant was transferred to a fresh tube and the pellet was extracted further with addition of 3x volume/weight of Triton X-100 buffer (TX, TS + 2 % Triton X-100). The samples were mixed by vortex, incubated on ice for 10 min and centrifuged again using the same prior setting. The TX supernatant was transferred to a fresh tube and the pellet was extracted further with addition of 3x volume/weight of SDS buffer (SDS, TS + 2 % SDS). The samples were mixed by vortex, incubated at room temperature for 10 min and centrifuged again at 55,000 rpm and 12°C for 30 mins. The SDS supernatant was transferred to a fresh tube and the pellet was either stored at −80°C or extracted further with addition of 3X volume/weight of 6X non-reducing Laemmli buffer (LB, 30 % SDS, 60 % glycerol, 0.3 % bromophenol blue, 0.375 M Tris. pH 6.8, 100mM DTT), mixed by vortex and incubated at room temperature for 10 min. Samples were centrifuged again at 55,000 rpm and 12°C for 30 mins and the LB supernatant was transferred to a fresh tube. Extracted proteins from TS, TXS and SDS buffers including pellet (20-30 μg) and 10-20 μL of LB extracts were run on SDS-PAGE using reducing (100 mM dithiothreitol, DTT) and/or non-reducing (0 mM DTT) loading buffer. Following transfer to membranes, Ponceaus S staining was used to confirm loading, and samples were blotted for parkin (Biolegend 808503, 1: 5,000), DJ-1 (ab18257, 1: 2,000), α-synuclein (syn1 1:1,000 or MJFR1 1:2000), LC3B (3868 1:2000), VDAC (MSA03 1:5000), MnSOD and GLO1 (1:1000), calnexin (MAB3126), cathepsin D (sc-6486), GRP75 (sc-1058). ImageJ software (1.52 k, National Institutes of Health, USA) was used for signal quantification purposes.

### mRNA Analysis

*PRKN* mRNA isolated from individual *S. nigra* dopamine neurons (SNDA), cortical pyramidal neurons (PY) and non-neuronal, blood mononuclear cells (NN) were processed, as described [39] and as annotated in the Human BRAINcode database (www.humanbraincode.org).

### ROS (H_2_O_2_) measurements in recombinant protein preparations, tissues and cell lysates

Amplex® Red hydrogen peroxide/peroxidase assay kit (Invitrogen A22188) was used to monitor endogenous levels of H_2_O_2_ in tissues and cells, and residual levels of H_2_O_2_ after incubation with recombinant parkin (WT, or pre-incubated with increasing concentrations of H_2_O_2_, NEM, or EDTA), DJ-1, SNCA, BSA, RNF43 (BioLegend), HOIP (Boston Biochem), GSH, catalase, NEM and EDTA for 30 minutes. Pre-weighed cortex pieces from human brains (or pelleted cells) were homogenized on ice in the 1x reaction buffer provided, using a Dounce homogenizer (3 times volume to weight ratio). Homogenates were diluted in the same 1x reaction buffer (10x and 5x). A serial dilution of the H_2_O_2_ standard provided was prepared (20, 10, 2 and 0 μM). 50 μL of standards and samples were plated in a 96 well black plate with clear flat bottom. The reaction was started by the addition of 50μL working solution which consisted of 1x reaction buffer, Amplex® red and horseradish peroxidase. The plate was incubated at room temperature for 30 minutes protected from light. A microplate reader was used to measure either fluorescence with excitation at 560 nm and emission at 590 nm, or absorbance at 560 nm. The obtained H_2_O_2_ levels (μM) were normalized to the tissue weight (g) or protein concentration (μg/μL). The same assay was also used to measure parkin and glutathione’s peroxidase activity compared to horseradish peroxidase (HRP).

### Recombinant protein expression using a pET-SUMO vector

Wild-type and truncated (amino acid 321-465) human parkin proteins were expressed as 6His-Smt3 fusion proteins in *Escherichia coli* BL21 (DE3) Codon-Plus RIL competent cells (C2527, New England Biolabs) as previous described [40–42]. *DJ-1* and *SNCA* coding sequences were cloned from a pcDNA3.1 vector into the pET-SUMO vector using PCR and restriction enzymes. ARPD-associated parkin mutants in the pET-SUMO vector were generated using site-directed mutagenesis. All proteins were overexpressed in *Escherichia coli* BL21 Codon-Plus competent cells (C2527, New England Biolabs) and grown at 37 °C in 2 % Luria Broth containing 30 mg/L kanamycin until OD600 reached 0.6, at which point the temperature was reduced to 16°C. All parkin-expressing cultures were also supplemented with 0.5 mM ZnCl_2_. Once OD600 reached 0.8, protein expression was induced with isopropyl β-D-1-thiogalactopyranoside, except ulp1 protease, which was induced once OD_600_ had reached 1.2. The concentration of isopropyl β-d-1-thiogalactopyranoside (IPTG) used for each construct is as follows: 25 μM for wild-type and point mutants of parkin, and 0.75 mM for truncated parkin, DJ-1, α-synuclein, and ulp1 protease. Cultures were left to express protein for 16-20 h. Cells were then harvested, centrifuged, lysed and collected on Ni-NTA agarose beads in elution columns.

Plasmid encoding for human Parkin with a p.C95A substitution was generated with the use of a restriction-free cloning strategy (PMID: 20600952) using the following primers: *PRKN* forward: CAGAAACGCGGCGGGAGGCgcTGAGCGGGAGCCCCAGAGCT and *PRKN* reverse: CATCCCAGCAAGATGGACCC.

### Protein redox chemistry and oxidation of cysteine-containing proteins *in vitro*

The recombinant protein samples were first prepared by removing excess TCEP, present in the elution buffer using repeat centrifugations (8 times 4000 × g at 4°C for 10 min) in Amicon Ultra 10kDa MWCO filters. The protein concentrations were measured and adjusted to 20μM. Stock solutions of hydrogen peroxide (H_2_O_2_, 9.8 mM) were prepared.

Aminochrome was freshly synthesized from dopamine (see below). An aliquot of 10μL of each protein sample (at 20 μM) was reacted with oxidants at the following concentrations: 0, 2, 20, 50, 200 aminochrome; 0, 20, 200, 500, 750, 1000, 2000 μM H_2_O_2_ or 0, 10, 50, 100, 200, 500, 1,000 μM DTT. The samples were treated for 30 min at 37°C and centrifuged at 14,000 rpm for 15 min. The supernatant was transferred to a fresh tube and the remaining pellet was extracted with 10μL of T200-TCEP containing either 10 % SDS or 100 mM DTT. The pellets were incubated again for 30 min at 37°C and centrifuged at 14,000 rpm for 15 min. Laemmli buffer (10 μL, containing 100 mM mercaptoethanol) was added to both the pellet and supernatant fractions and samples were separated on two SDS-PAGE. One gel was used for in-gel protein staining and the other was used for NBT staining. Specific bands of aminochrome treated wild-type, full length r-parkin were excised from silver-stained gels and analyzed by LC-MS/MS as described below.

### Aminochrome synthesis

A solution of 0.1 M sodium phosphate buffer pH 6.0 was prepared from a mixture of 12 mL of 1M NaH_2_PO_4_ and 88.0 mL of 1M Na_2_HPO_4_. The reaction buffer (0.067 M sodium phosphate, pH 6.0) was prepared by adding 33 mL of 0.1 M sodium phosphate buffer to 17 mL water. A solution of 10mM dopamine in reaction buffer was prepared by adding 19 mg of dopamine hydrochloride to 1 mL of reaction buffer. Oxidation was activated by adding 5 μL of tyrosinase (25,000 U/mL) and the mixture was incubated at room temperature for 5 min. The tyrosinase was separated from the oxidized dopamine using a 50 kDa cut-off Amicon Ultra centrifugation filter by centrifuging at 14,000 rpm for 10 min. The absorbance of the filtrate was measured at a wavelength of 475 nm using Ultrospec 21000 pro spectrophotometer and the concentration of aminochrome was determined using the Beer-Lambert equation and extinction coefficient of 3058 L × mol^−1^ × cm^−1^.

### Protein staining methods

All proteins were separated on pre-cast 4-12 % Bis-Tris SDS-PAGE gels (NPO321BOX, NPO322BOX, NPO336BOX) from Invitrogen using MES running buffer (50mM MES, 50mM Tris, 1mM EDTA and 0.1 % SDS, pH 7.3) and Laemmli loading buffer (10% SDS, 20% glycerol, 0.1% bromophenol blue, 0.125M Tris HCl, 200mM DTT or β-mercaptoethanol). Proteins were stained in gel using SilverQuest™ Silver Staining Kit (LC6070) from Invitrogen or Coomassie brilliant blue R-250 dye (20278) from ThermoFisher Scientific using the following protocol: The gel was transferred to a plastic container and rocked for 30 min in Fix Solution (10% acetic acid, 50% methanol), followed by staining for 2-24 h (0.25% Coomassie R250) until the gel turned a uniform blue. The stain was replaced with Destain Solution (7.5% acetic acid and 5% methanol) and the gel was rocked until crisp blue bands appeared. Following a wash with water the gel was stored in 7 % acetic acid. Proteins transferred to PVDF (1620177, Bio-Rad) membranes were stained with Ponceau S solution for 20 min, washed three times with water, imaged and then destained with 0.1M NaOH prior to Western blotting.

### Dynamic light scattering assay

For each recombinant protein preparation tested, the buffer (50mM Tris, 200mM NaCl and 250μM TCEP, pH 7.5) was exchanged for a 20mM phosphate buffer with 10mM NaCl (pH 7.4). 20 μM full-length wild-type r-parkin was centrifuged at 14,000 rpm for 60 min at 4 °C and light scattering intensity of the supernatant was collected 30 times at an angle of 90° using a 10 sec acquisition time. Measurements were taken at 37 °C using a Malvern Zetasizer Nano ZS instrument equipped with a thermostat cell. The correlation data was exported and analyzed using the nanoDTS software (Malvern Instruments). The samples were measured at 0-, 1-, 3- and 5 hours. Following 24 hr incubation, 2 mM DTT was added to the sample and the light scattering intensity of the supernatant was measured again.

### Far UV circular dichroism spectroscopy

15 μM of reduced and partially oxidized full-length wild-type r-parkin was measured at t = 0 and t = 5 days of incubation under native conditions in 20 mM phosphate, 10 mM NaCl buffer. The aggregates rich phase and the monomer rich phase in the samples were separated with ultracentrifugation (100,000 g for 2 hours). Far UV circular dichroism (CD) spectra were recorded for the monomer and aggregated rich phase of protein samples using a JASCO J-720 spectrometer. The final spectrum was taken as a background-corrected average of 5 scans carried out under the following conditions: wavelength range 250–190 nm at 25 °C; bandwidth was 1 nm; acquisition time was 1 sec and intervals was 0.2 nm. Measurements were performed in a 0.01 cm cell. CD spectra were plotted in mean residue molar ellipticity units (deg cm^2^ dmol^−^1) calculated by the following equation: [Θ] = Θ_obs_/(10*ncl)*, where [Θ] is the mean residue molar ellipticity as a function of wavelength, Θ_obs_ is the measured ellipticity as a function of wavelength (nm), *n* is the number of residues in the protein, *c* is the concentration of the protein (M), and *l* is the optical path length (cm). Secondary structure analysis of proteins using CD spectroscopic data was carried out using the BeStSel (Beta Structure Selection) software [43, 44].

### Chemiluminescence-based, direct reactive oxygen species (ROS) assay

The assay was modified from Muller et al. 2013 [45] to measure the ROS-quenching ability of parkin proteins, DJ-1, SNCA, BSA, GSH, and catalase. Protein concentrations were quantified using Bradford assay and adjusted to 5, 10, 15 and 30 μM in buffer not containing TCEP. BSA (10 and 20 μM), GSH (15, 20, 200, 400, 800 and 2000 μM), and catalase (0.015, 0.15, 0.25 and 15 μM) were prepared. Stock solutions of H_2_O_2_ for standard curve were prepared at 5, 10, 20, 40 and 50 mM in 0.1 M Tris HCl pH 8.0 using 30 % H_2_O_2_. Stock solutions of 300 mM luminol and 40 mM 4-iodophenol were prepared in DMSO and protected from light. Signal reagent, containing 1.94 mM luminol and 0.026 mM 4-iodophenol, was prepared in 0.1 M Tris HCl pH 8.0 and protected from light. A 0.4 % horseradish peroxidase solution was prepared using HRP-linked anti-rabbit secondary antibody diluted in Stabilizyme solution (SurModics SZ02). Each read was set up in triplicate on a white polystyrene 96-well plate (ThermoFisher 236105) and to each well was added 80 μL Stabilizyme, 15 μL of 0.4 % horseradish peroxidase (HRP) and 25 μL of sample or controls. One of the injectors in a Synergy H1Multi-Mode Plate Reader (Bio Tek) was primed and set to inject 15 μL of signal reagent and 15 μL of each H_2_O_2_ stock solution was manually added to corresponding controls and samples just prior to reading. Final concentrations of reagents were 0.04 % HRP, 500, 1000, 2000, 4000 and 5000 μM H_2_O_2_, 194 μM luminol, 2.6 μM 4-iodophenol and 0.8, 1.7, 2.5 or 5 μM of protein. The plate reader was set to measure luminescence every 1 min for a total of 10 min. The resulting kinetic data was converted to area under the curve (AUC) using Prism version 6. For samples pre-incubated with 20 mM iodoacetamide, a stock solution of 1 M iodoacetamide was prepared. To each well containing 25 μL of sample, 0.52 μL of 1 M iodoacetamide and 0.48 μL of buffer not containing TCEP was added and the samples were incubated for 2 h at 37°C. Following incubation, the reagents for chemiluminescence were added as above except 79 μL of Stabilizyme was used instead of 80 μL and the samples were analyzed as above.

### Thiol quantification in recombinant proteins

Recombinant protein samples were first prepared by exchanging the T200 protein buffer (50 mM Tris, 200 mM NaCl and 250 μM TCEP, pH 7.5) for T200-TCEP using repeat centrifugations (8 times 4000 × g at 4°C for 10 min) in Amicon Ultra 10 kDa MWCO filters. The protein concentrations were measured and recorded. The glutathione stock solution of 32,539 μM was prepared by dissolving 1 mg glutathione (GSH) in 1 mL of T200-TCEP and the standards 0, 50, 101, 203, 406, 813 and 1000 μM were prepared by serial dilution in T200-TCEP. The reaction buffer (0.1 M sodium phosphate, pH 8.0) was prepared by adding 93.2 mL 1M Na_2_HPO_4_ and 6.8 mL of NaH_2_PO_4_ in 1 L of water. Thiol detecting reagent (Ellman’s reagent) was prepared by dissolving 2 mg of 5,5’-dithio-bis-[2-nitrobenzoic acid] (DNTB) in 1 mL of reaction buffer. The assay was performed in 96-well clear round bottom plates by adding 50 μL of thiol detecting reagent to 50 μL of sample or standard and incubating for 15 min at room temperature. The resulting 5-thio-2-nitrobenzoic-acid (TNB) produced was measured by absorbance at 412 nm using a Synergy H1Multi-Mode Plate Reader (Bio Tek). The amount of free thiols detected in each sample was calculated using the regression curve obtained from the glutathione standards and dividing by the concentration of the sample.

### Cysteine labeling for mass spectrometry

The recombinant protein samples were first prepared by exchanging the T200 buffer for PBS. The protein concentrations were measured and adjusted to 10 μM using PBS. Stock solutions of 500 mM DTT, 100 mM iodoacetamide (IAA), 100 mM hydrogen peroxide and 250 mM ethylenediaminetetraacetic acid (EDTA) were prepared in PBS. A stock of 500 mM N-ethyl-maleimide (NEM) was prepared in ethanol immediately before use. For the first optimization and comparison of IAA and NEM labelling (*i.e.*, Supplementary Table 2), r-parkin was treated with 2 mM DTT for 30 min at 37°C followed by incubation with 5 mM IAA or 85 mM NEM for 2 h at 37°C. The stepwise Cys labeling procedure was as follows: A 10 μL aliquot of protein (at 10 μM) was reacted with hydrogen peroxide at various concentrations, as indicated (Table 1) for 30 min (and up to 60 min) at 37°C as indicated. Any unreacted cysteines were alkylated with incubation with 5 mM IAA (either with or, in some runs, without 10 mM EDTA) for 2 hrs at 37°C. Previously oxidized cysteines were then reduced by treatment with 40 mM DTT for 30 min at 37°C. Newly reduced cysteines were alkylated by incubation with 85 mM N-ethyl maleimide (NEM) for 2 hrs at 37°C. The samples were separated on SDS-PAGE using Laemmli buffer containing 100 mM DTT and proteins visualized using Coomassie staining. Appropriate bands were excised and analyzed by liquid chromatography mass spectrometry (LC-MS/MS).

### Protein identification by LC-MS/MS

Proteomics analysis was performed at the Ottawa Hospital Research Institute Proteomics Core Facility (Ottawa, Canada). Proteins were digested in-gel using trypsin (Promega) according to the method of Shevchenko [46]. Peptide extracts were concentrated by Vacufuge (Eppendorf). LC-MS/MS was performed using a Dionex Ultimate 3000 RLSC nano HPLC (Thermo Scientific) and Orbitrap Fusion Lumos mass spectrometer (Thermo Scientific). MASCOT software version 2.6.2 (Matrix Science, UK) was used to infer peptide and protein identities from the mass spectra. For detection of dopamine metabolites on Parkin, the following variable modifications were included: 5,6-indolequinone (+C_8_O_2_NH_3_, m/z shift +145), aminochrome (+C_8_O_2_NH_5_, +147), aminochrome +2H (+C_8_O_2_NH_7_, +149), and dopamine quinone (+C_8_O_2_NH_9_, +151). These samples were prepared for analysis without any use of dithiothreitol or iodoacetamide. The observed spectra were matched against human sequences from SwissProt (version 2018-05) and also against an in-house database of common contaminants. The results were exported to Scaffold (Proteome Software, USA) for further validation and viewing. Analysis of the holoprotein and of three runs of H_2_O_2_-exposed r-parkin (Supplemental Table 2) were performed at the University of Western Ontario. There, samples were run on a QToF Ultima mass spectrometer (Waters) equipped with a Z-spray source and run in positive ion mode with an Agilent 1100 HPLC used for LC gradient delivery (University of Western Proteomics Facility).

### MaxQuant analysis of mass spectrometry data

For applicable experiments, the raw MS data files were further processed with MaxQuant software version 1.6.5 and searched with the Andromeda search engine[47]. The reference fastas were set to uniprot-human (version 2019-02-12) and uniprot-ecoli. The E. coli proteome was included to account for bacterial proteins present in the recombinant protein samples. The ‘second peptides’ and ‘match between runs’ settings were enabled. All other settings were left as default. Selected variable modifications included oxidation (Met), acetylation (protein N-terminus), and carbamidomethyl (Cys), as well as custom modifications for pyro-carbamidomethyl (N-terminal Cys), N-ethylmaleimide (Cys), and NEM+water (Cys). For data analysis, site-level intensity values were obtained from the MaxQuant-generated “CarbamidomethylSites” table which combines the intensity of MS1 signals from all peptides covering a particular cysteine residue.

### Immunoprecipitation (IP) of brain parkin

Conjugation of anti-parkin antibody (Prk8, 808503, lot B209868) and clone A15165-B (this report: Suppl. Fig. 8c) to magnetic beads at a final concentration of 10 mg of antibody / mL of beads was done following the Magnetic Dynabeads Antibody Coupling Kit from Invitrogen (14311D). Human tissue lysates were also prepared using the “Sequential Extraction of Proteins from Tissue” protocol as described above with addition of 10 mM iodoacetamide prior to homogenization. TS tissue extracts (n=4) and SDS tissue extracts (n=8) were diluted in TS buffer, resulting in a final SDS concentration of 0.0175 % and 0.05 % respectively. For the IP, Prk8 conjugated agarose beads were first prepared by multiple washes with 1 mL of TS buffer using centrifugation (1000 × g at 4°C for 3 min) and adhesion to a strong magnet. Amounts of Prk8 conjugated agarose beads used for each experiment were approximated based on the amount of parkin (μg) / sample calculated by densitometry when the sample was compared to recombinant parkin protein standards using Western blotting with Prk8 primary antibody. The mixture was incubated for 16 h at 4°C with slow rotation. Unbound proteins, which did not bind to the Prk8 conjugated agarose beads, were separated from the beads by centrifugation (1000 × g at 4°C for 3 min) followed by adhesion to a strong magnet and saved as the IP “unbound” fraction.

Beads from cellular or human IP were washed three times with 900 or 1000 μL respectively of ice-cold RIPA buffer (1 % nonionic polyoxyethylene-40, 0.1 % SDS, 50 mM Tris, 150 mM NaCl, 0.5 % sodium deoxycholate, 1 mM EDTA) using centrifugation (1000 × g at 4°C for 3 min) and adhesion to a strong magnet. Approximately 5-10 μL of each wash was combined and saved as the IP “wash” fraction. To elute Prk8 bound proteins, 15-35 uL of 6X reducing Laemmli buffer (30 % SDS, 60 % glycerol, 0.3 % bromophenol blue, 0.375 M Tris, 100 mM DTT, pH 6.8) was added to the beads and the samples were boiled for 5 min. Following centrifugation (1000 × g at 4°C for 3 min), the supernatant was transferred to a fresh tube labeled “IP elute” and the beads were discarded. To assess IP efficiency, eluted fractions (IP elute), along with controls (input, unbound, wash and recombinant parkin protein standards) were run on SDS/PAGE and blotted with anti-parkin antibody (MAB5512 or 2132S). Human IP elutes used in subsequent for mass spectrometry (MS) analysis were incubated with 500 mM N-ethyl maleimide (as indicated for select runs) for 16 h at 4°C prior to SDS-PAGE and further processing for MS (as described above). Gel slices corresponding to band sizes 50-75 kDa were excised and analyzed by LC-MS/MS.

### *In vitro* melanin formation assay

The recombinant protein samples were first prepared by exchanging the T200 protein buffer (50 mM Tris, 200 mM NaCl and 250 μM TCEP, pH 7.5) for T200-TCEP (50 mM Tris and 200 mM NaCl, pH 7.5) using repeat centrifugations (8 times 4000 × g at 4°C for 10 min) in Amicon Ultra 10 kDa MWCO filters. The protein concentrations were measured and adjusted to 20 μM using T200-TCEP. A 0.067 M sodium phosphate buffer, pH 6.0, was prepared by adding 33 mL of 0.1 M sodium phosphate buffer to 17 mL water and adjusting the pH using HCl. A stock solution of 100 mM dopamine HCl was prepared in 0.067 M sodium phosphate buffer and stock solutions of 100 mM reduced glutathione (GSH) and hydrogen peroxide were prepared in T200-TCEP.

Samples and controls were prepared in 100 μL total volume and contained: 10 μL of 20 μM protein or T200-TCEP, 10 μL of 100 mM dopamine or 0.067 M sodium phosphate buffer, 10 μL of 100 mM glutathione or T200-TCEP buffer, and 70 μL T200-TCEP. The final concentration of protein was 2 μM and the final concentration of reagents was all 10 mM. The samples and controls were plated in triplicate, and absorbance read at 405 and 475 nm every 90 sec for 1 h and up to 4 h.

### 1-methyl-4-phenyl-1,2,3,6-tetrahydropyridine (MPTP) treatment

Eight to 12 mths-old WT and *prkn*-null mice were injected intraperitoneally with 40mg/kg of saline or MPTP and sacrificed an hour later[48]. The brains were harvested for ROS measurement, protein analysis by Western blot and immunoprecipitation of parkin and mass spectrometry analysis. For mass spectrometry, the brains harvested were first incubated in IAA prior to homogenization and fractionation as described above. Brain homogenates were then incubated with anti-parkin conjugated to magnetic beads (Dynabeads Coupling Kit; Invitrogen). A magnet was used to capture parkin bound to the beads, and several washes were used to remove unbound proteins. Eluted fractions (IP elute) along with controls (input, unbound, wash and recombinant parkin protein standards) were run on SDS/PAGE and blotted with anti-parkin. A sister gel was stained with Coomassie as described above and gel slices corresponding to band sizes 50-75 kDa were excised and analyzed by LC-MS/MS, as described in detail by Tokarew et al., 2020.

### Cell cytotoxicity assay

Human neuroblastoma cell line (M17 cells) wild-type, Vector (Myc), P5 (low stable expression of Myc-parkin) and P17 (high stable expression of Myc-parkin), or WT ectopically overexpressing flag-parkin (WT), flag vector and flag-parkin carrying the following mutations (p.C431F, p.G3289E and p.C95A) were grown in 6 well culture plates, at 0.3×10^6^ cell density (80% confluence) in Opti-MEM media (Gibco 11052-021) containing heat inactivated FBS (Gibco 10082-147), Pen/strep/Neo (5mg/5mg/10mg) (Gibco 15640-055), MEM non-essential amino acids (10mM) (Gibco 11140-050) and sodium pyruvate (100mM). For rescue experiments, flag-vector, flag-parkin, flag-p.G328E, flag-p.C431F and flag-p.C95A-encoding pCI-neo plasmids were expressed in M17 wild-type cells. There, 4 μg of cDNA was transfected using a 1:1 ratio of cDNA: Lipofectamine 2000 (52887, Invitrogen) in OPTI-MEM transfection medium. The cDNA and Lipofectamine 2000 was first incubated for 20 min at room temperature before being applied directly to the cells for 1 h at 37°C with 5 % CO_2_ followed by direct addition of fresh growth medium. The cells were incubated another 24 hours at 37°C with 5 % CO_2_.

Dopamine hydrochloride (Sigma H8502) 200 mM stock was prepared. The cells were washed with fresh media once and then incubated with media alone or supplemented with dopamine at final concentrations of 20 μM and 200 μM for 18-20 hours. Post dopamine stress, media was collected from all wells for cytotoxicity assay, the cells were harvested and lysed with TS buffer and centrifuged. The supernatant was collected and saved for Western blot analysis and to assess total cell toxicity signal. The pellet was suspended in SDS buffer and centrifuged.

Vybrant ™ cytotoxicity assay kit (Molecular Probes V-23111) was used to monitor cell death through the release of the cytosolic enzyme glucose 6-phosphate dehydrogenase (G6yPD) from damaged cells into the surrounding medium. 50 μl of media alone (no cells), media from control and stressed cells and cell lysates were added to a 96-well microplate. Fifty μl of reaction mixture, containing reaction buffer, reaction mixture and resazurin, was added to all wells, and the mircroplate was incubated at 37°C for 30 mins. A microplate reader was used to measure either fluorescence with excitation at 560 nm and emission at 590 nm. A rise in fluoresence indicates a rise in G6PD levels i.e. a rise in cell death.

### Immunohistochemistry (IHC)

Immunohistochemistry was performed on paraffin-embedded sections and treated as previously described [49–51]. Briefly, prior to antibody incubation, sections were deparaffinized in xylene and successively rehydrated through a series of decreasing ethanol concentration solutions. Endogenous peroxidase activity was quenched with 3% hydrogen peroxide in methanol, followed by a standard citric acid-based antigen retrieval protocol to unmask epitopes. Sections were blocked in 10-20% serum in PBS-T to reduce non-specific signal. Sections were incubated overnight at 4°C in primary antibodies diluted in 1-5% serum in PBS-T according to the following concentrations: novel anti-parkin mAbs from Biolegend clones D (BioLegend, A15165D; 1:250), clone E (BioLegend, A15165E; 1:2000), and clone G (1:250), PRK8 (BioLegend, MAB5512; 1:500) as well as anti-LAMP-3/CD63 (Santa Cruz, SC5275; 1:100), anti-LC3B (Sigma, L7543-200uL; 1:100), anti-VDAC (MitoScience, MSA03; 1:100). Biotinylated secondary antibodies (biotinylated anti mouse IgG (H+L) made in goat; Vector Labs, BA-9200, biotinylated anti-rabbit IgG (H+L) made in goat; Vector Labs, BA-1000) were diluted to 1:225 and sections were incubated for 2 hours at room temperature. The signal was amplified with VECTASTAIN® Elite® ABC HRP Kit (Vector Labs, PK-6100), and visualized via standard DAB solution, 55mM DAB, or Vina green (Biocare Medical, BRR807AH), or most commonly metal enhanced DAB (Sigma, SIGMAFAST™ DAB with Metal Enhancer D0426). Samples were counterstained with Harris Modified Hematoxylin nuclei stain and dehydrated through a series of increasing ethanol concentration solutions and xylene. Permount (Fisher Scientific, SP15-100) was used for mounting and slides were visualized with high magnification images via a Quorum Slide Scanner (Ottawa Hospital Research Institute).

### Immunofluorescence (IF) and confocal microscopy

Paraffin-embedded human midbrain sections were stained by routine indirect immunofluorescence with the following details. Antigen retrieval was performed in Tris-EDTA buffer pH 9 for 10 mins. Primary antibodies were incubated overnight at 4C. Details for primary antibodies anti-Parkin Clone E (1:500), anti-LAMP3 (1:250) are described above. A 40 minute incubation with the following secondary antibodies was performed: goat anti-mouse alexa fluor 488 (1:200), goat anti-rabbit alexa fluor 594 (1:500). Slides were mounted with fluorescence mounting medium with DAPI. Stained sections were imaged using a Zeiss LSM 880 AxioObserver Z1 with an Airyscan Confocal Microscope then processed and analyzed using Zeiss Zen and Fiji software.

### Statistical analyses

All statistical analyses were performed using GraphPad Prism version 6 (GraphPad Software, San Diego, CA, USA, www.graphpad.com). Differences between two groups were assessed using an unpaired t-test. Differences among 3 or more groups were assessed using a one-way or two-way ANOVA followed by Tukey’s post hoc corrections to identify statistical significance. Subsequent post hoc tests are depicted graphically and show significance between treatments. For all statistical analysis a cut-off for significance was set at 0.05. Data is displayed with p values represented as *p < 0.05, **p < 0.01,***p < 0.001, and ****p < 0.0001. Linear regression (for continuous dependent variable, e.g., H_2_O_2_ level, mRNA level) or logistic regression (for binary dependent variable, *e.g.,* parkin present in TS fraction) modelling were performed. Furthermore, to address the effect of age on parkin solubility, receiver operating characteristic (ROC) curve and area under the ROC curve (AUC) were calculated, as reported [51].

## Results

### Parkin is mostly insoluble in the ageing human brain including the *S. nigra*

Parkin’s metabolism in the human brainstem *vs.* other regions has remained largely unexplored [52]. We serially fractionated 20 midbrain specimens (ages, 26-82 yrs) and >40 cortices (ages, 5-85 yrs) from human subjects (**Fig. 1**, **Supplementary Fig. 1**; **Supplementary Table 1**). In control brain, we found that before the age of 20 yrs, nearly 50% of cortical parkin was found in soluble fractions generated by salt [Tris-NaCl; TS]- and mild detergent [Triton X-100; TX]-containing buffers (**Fig. 1a,b; Supplementary Fig.1a**). In contrast, after age 50 yrs, parkin was found almost exclusively (>90%) in the 2% SDS-soluble (SDS) fraction and the 30% SDS extract of the final fractionation pellet (P). The same distribution was seen in adult midbrain (*e.g., S. nigra*; red nucleus), the pons (*e.g.*, *L. coeruleus*), and the striatum (**Fig. 1a,b; Supplementary Fig. 1a-c**).

**Figure 1:**
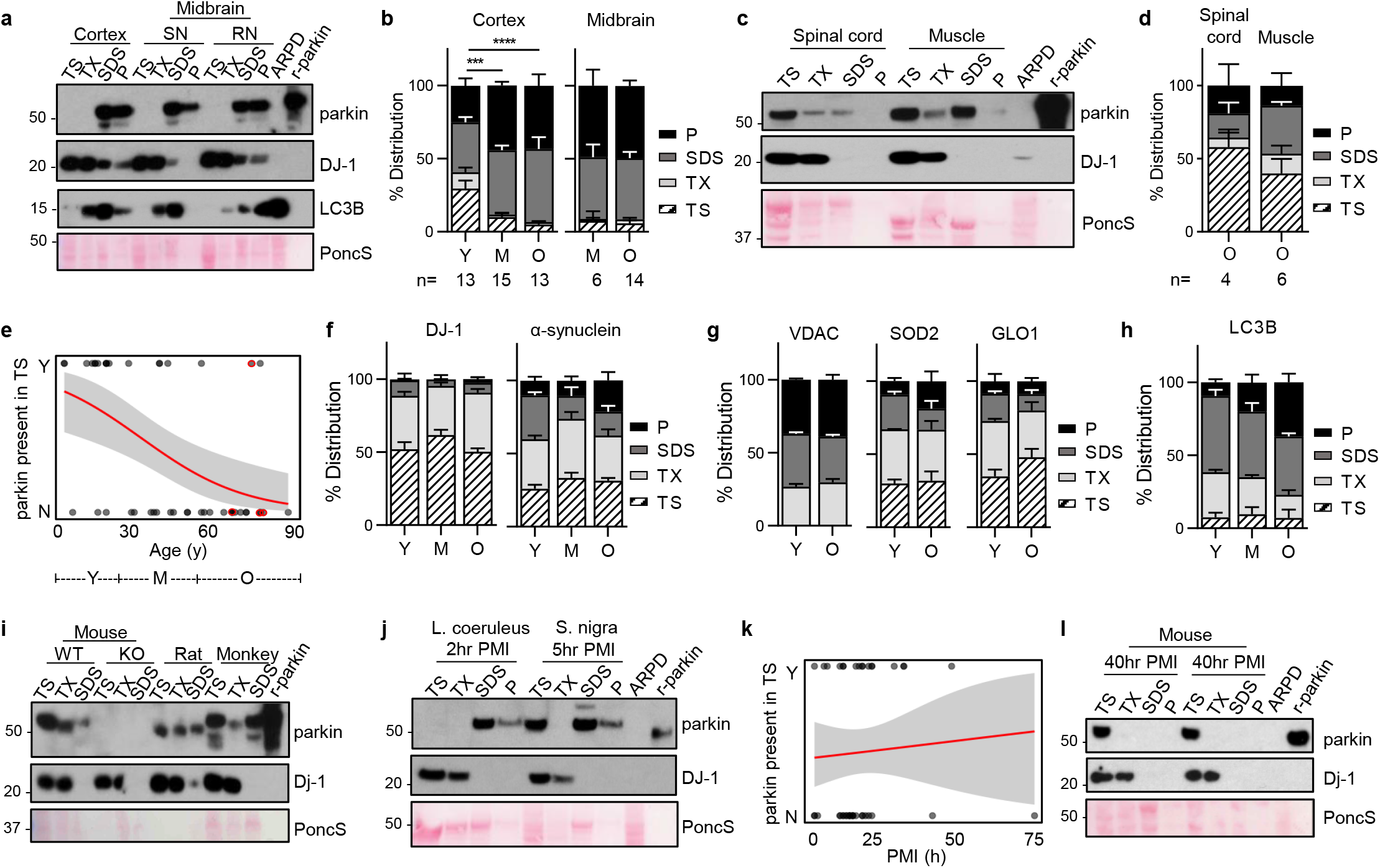
Parkin’s loss of solubility is specific to adult human brain and correlates with age. **(a)** Representative Western blots of parkin, DJ-1, and LC3B distribution in human cortex, *S. nigra* (SN) and red nucleus (RN) serially fractionated into Tris-NaCl buffer-soluble (TS), Triton X-100-soluble (TX), 2% SDS-soluble (SDS) extracts and the pellet (P) lysed in 30% SDS-containing buffer. SDS extracts from *PRKN*-linked Parkinson disease (ARPD) brain and recombinant, human parkin (r-parkin) are included. Ponceau S is shown as loading control. **(b)** Relative distribution of parkin signal within each fraction for cortex and midbrain grouped by age ranges: young (Y, ≤20 yrs; n=13); mid (M, >20 yrs but <50 yrs; n=15 for cortex, and n=6 for midbrain); older (O, ≥50 yrs; n=13 for cortex and n=14 for midbrain). Data shown as mean ± SEM. The significance in protein distribution between soluble (TS+TX) and insoluble (SDS+pellet) fractions was determined using 2-way ANOVA (***P<0.001; ****P<0.0001). Additional Western blots are shown in **Extended Data Fig. 1a-c**. Midbrains include both control and neurological disease cases, as listed in **Extended Data Table 1**. **(c)** Western blots of parkin and DJ-1 as well as Ponceau S staining of serial fractions from representative human spinal cord and skeletal muscle tissues from individuals >50 yrs. **(d)** Relative distribution of parkin as in (b) for human spinal cord (n=4) and skeletal muscle specimens (n=6) from donors aged 50-71 yrs. **(e)** Logistic regression analysis of parkin solubility in cortices as a function of age (n=45). Each brain is represented by an individual dot; red circles denote three cases of late-onset Parkinson’s not linked to *PRKN*; the logistic regression line (in red) and 95% confidence intervals (grey) are shown. Age ranges that correspond to Y-O-M in (b) are shown under the graph. **(f-h)** Relative distribution of **(f)** DJ-1, a-synuclein and **(g)** VDAC, MnSOD, glyoxalase (GLO1) and **(h)** LC3B in human cortices (n=3-5 per age group), as described in (b). Representative Western blots are shown in **Extended Data Fig. 1b,c**. **(i)** Western blots of parkin and Dj-1 and Ponceau S staining of serial fractions from whole brains of wild-type (WT; 8 mths of age) and *prkn* knock-out (KO) mice, WT rat (WT; 14 mths) and from frontal cortex of a cynomolgus monkey (60 mths). **(j)** Western blots of parkin and DJ-1 distribution in two human brainstem nuclei, *L. coeruleus* and *S. nigra*, which were collected within 2-5 hrs after death prior to freezing and processed as in (a, c). **(k)** Logistic regression analysis of parkin solubility in human brain as a function of length for *post mortem* interval (PMI; in hrs); the logistic regression analysis line (red) and 95% confidence intervals (grey) are shown (n=45 cortices). **(l)** Immunoblots for endogenous parkin and Dj-1 as well as ponceau S staining from serially extracted WT mouse brains (n=3) dissected after a 40 hr *post mortem* interval at 4°C.

Intriguingly, approximately half of detectable parkin remained soluble in human spinal cord and skeletal muscle specimens from older individuals (ages, ≥50 yrs) (**Fig. 1c,d**). We used logistic regression modeling to demonstrate a robust, negative correlation between parkin solubility in human control brain and age (**Fig. 1e**); the age coefficient was −0.0601 (95% CI: −0.106 to −0.024; P=0.004). The transition to insoluble parkin occurred between the ages of 28 yrs (at low sensitivity; high specificity values) and 42 yrs (high sensitivity; low specificity values; **Fig. 1e**).

The age-dependent partitioning of parkin was not seen for any other protein examined, including other PD-linked proteins, *e.g.,* DJ-1 and α-synuclein (**Fig. 1a,f**) and organelle-associated markers, *e.g.,* cytosolic glyoxalase-1, peroxiredoxin-1 and −3; and endoplasmic reticulum-associated calnexin. Notably, mitochondrial markers, *e.g.,* voltage-dependent anion channel (VDAC) and Mn^2+^-superoxide dismutase (MnSOD), also did not partition with parkin (**Fig. 1g; Supplementary Fig. 1b,c;** and data not shown). In contrast, parkin did co-distribute with LC3B, a marker of protein aggregation, foremost in the samples from older individuals (**Fig. 1a,h; Supplementary Fig. 1c**). The age-dependent loss of solubility for parkin was unique to human brain in that it remained soluble in the nervous system of other aged species, *e.g.,* mice, rats and cynomolgus monkey, which were processed in the same way (**Fig. 1i**).

In soluble fractions from older humans, we did not detect any truncated species of parkin using several, specific antibodies (data not shown). Despite the loss of parkin solubility with ageing, *PRKN* mRNA was detectable in individual neurons isolated from the *S. nigra* and cortex throughout all age groups; there, the transcript levels were independent of age (**Supplementary Fig. 1d,e**).

Most important, we also confirmed that parkin insolubility did not correlate with the length of *post mortem* interval (range, 2-74 hrs), as studied in both human and mouse brains (**Fig. 1j-l; Supplementary Fig. 2a,b**), was independent upon sex of the deceased person (not shown), and was not caused by either tissue freezing prior to protein extraction or the pH of the buffer used (**Supplementary Fig. 2c-f**). Further, using the commonly used ‘RIPA buffer’ instead of serial extraction buffers resulted in the release of parkin into the supernatant with some reactivity left in the pellet, as expected (**Supplementary Fig. 2g**).

### The insolubility of brain parkin correlates with rising hydrogen peroxide levels

We explored a possible association between parkin distribution, age and oxidative changes. Using sister aliquots from the brain specimens examined above, we found that hydrogen peroxide (H_2_O_2_) concentrations positively correlated with age (**Fig. 2a,b;** see also **Supplementary Table 1**), as expected from the literature [53]. In three brains of non*-PRKN*-linked cases of parkinsonism, the levels of H_2_O_2_ were similar to those of age-matched controls (**Fig. 2b**). When analyzing parkin distribution *vs.* H_2_O_2_ concentrations, we found that parkin solubility in human brain negatively correlated with H_2_O_2_, where the coefficient of the latter was - 0.939 (95% CI: −2.256 to −0.248; P=0.0415) (**Fig. 2c**).

**Figure 2:**
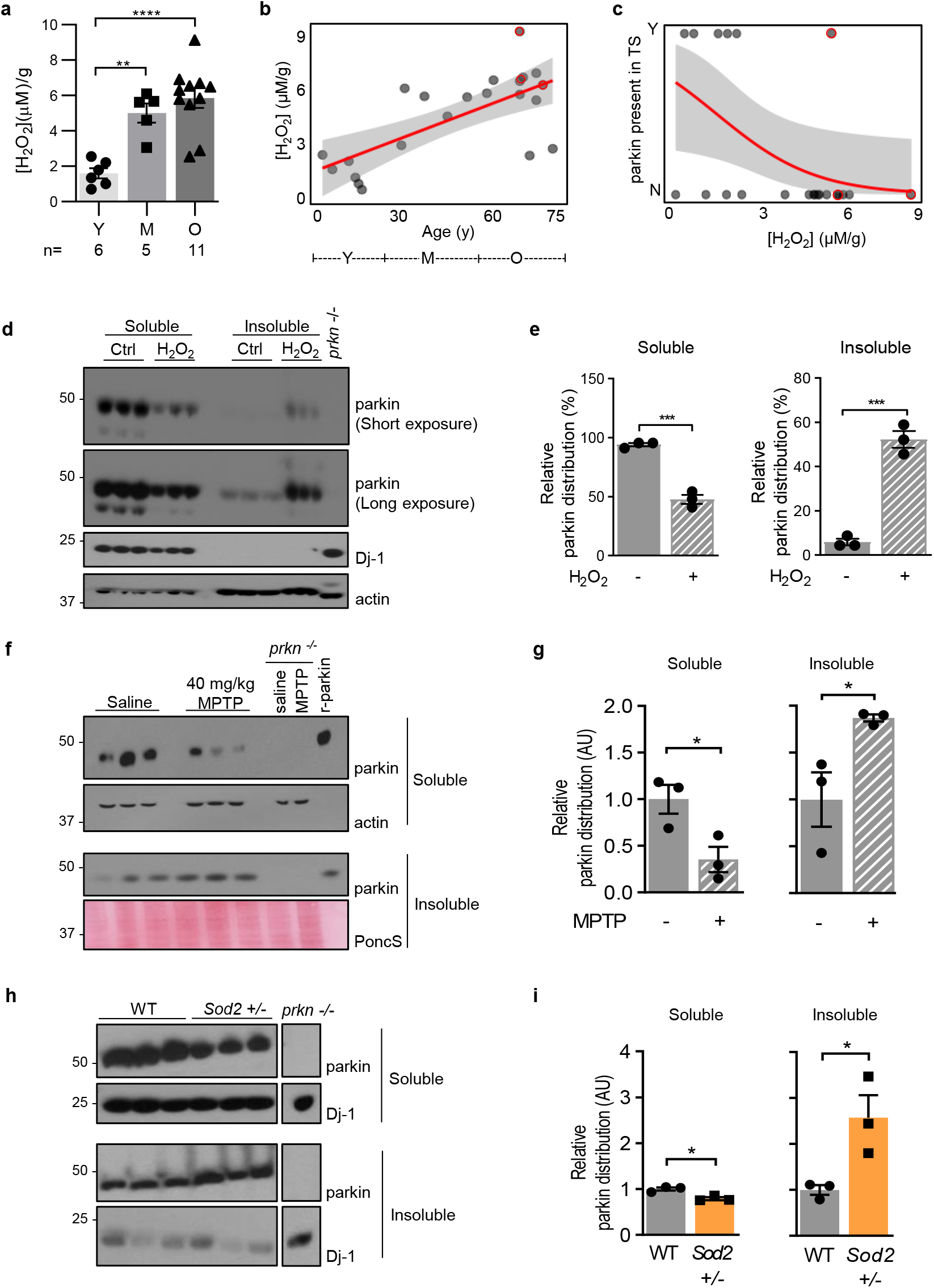
Parkin’s loss of solubility in the brain correlates with a rise in oxidative stress. **(a)** Mean concentrations of H_2_O_2_ in human brain cortices grouped by age range, as described in Figure 1. Individual data points represent separate brains, as reported in **Extended Data Table 1.** Results are plotted as mean ± SEM; significance was determined using 2-way ANOVA (**P<0.01; ***P<0.001). **(b-c)** Linear regression analysis of H_2_O_2_ concentrations in control cortices (mM/g tissue) as a function of age **(b**), and **(c)** logistic regression analysis of parkin solubility as a function of H_2_O_2_ levels in the same specimens (n=20). Red circles denote three disease cortices (AD; DLB; PD). **(d-e)** Western blots **(d)** of parkin distribution in brain lysates of 2-4 month-old wild-type C57Bl/6J mice containing either saline or 1% H_2_O_2_; **(e)** parkin signal distribution was quantified using image-J, as controlled for respective loading controls, in both soluble and insoluble fractions. A student t-test was used for statistical analysis (* = < 0.05). **(f-g)** Western blots **(f)** of parkin distribution in brains of wild-type C57Bl/6J mice 1 hour following intraperitoneal administration of either saline or MPTP neurotoxin (40mg/Kg); **(g)** parkin signals were quantified as in (e). **(h-i)** Western blots **(h)** of fractionated brain homogenates from C57Bl/6J wild-type and *Sod2*^+/−^ mice; **(i)** parkin signals were quantified and statistically analyzed as in (e) (* = < 0.05).

We next sought to validate the correlation between oxidative stress, ROS levels and parkin solubility in mice. We first used an *ex vivo* approach in which wild-type mouse brain homogenates were exposed to either saline or H_2_O_2_. There we saw a significant reduction in soluble parkin and an increase in insoluble parkin in H_2_O_2_-exposed lysates (**Fig. 2d,e**). We next examined two *in vivo* models. First, wild-type mice were injected intraperitoneally, one hour before sacrificing them, with 40 mg/kg of MPTP toxin to induce acute oxidative stress, but no cell death [48]. Brains were serially fractionated, and parkin distribution was quantified across soluble and insoluble compartments. There, we measured a decrease of murine parkin in the soluble fraction and a corresponding rise in the insoluble fractions of MPTP-*vs.* saline-injected animals (**Fig. 2f,g**). Second, we observed a similar shift in parkin distribution in adult mice that were haploinsufficient for the *Sod2* gene, which encodes mitochondrial MnSOD, in the absence of any exogenous toxin (**Fig. 2h,i**). Of note, in both models we confirmed the rise in H_2_O_2_ levels (see below and El Kodsi *et al.* [38]). In contrast to murine parkin, the solubility of endogenous Dj-1, encoded by a second, ARPD-linked gene, was not visibly affected on SDS/PAGE under these elevated oxidative stress conditions (**Fig. 2h**).

### Parkin is reversibly oxidized in adult human brain

The correlation of parkin solubility with H_2_O_2_ levels in human control brain suggested that its solubility could be associated with posttranslational, oxidative modifications. Indeed, in contrast to SDS-containing brain fractions carried out under reducing conditions (+dithiothreitol, DTT), when gel electrophoresis was performed under non-reducing (-DTT) conditions, we detected parkin proteins ranging in *M*_*r*_ from >52 to 270 kDa, invariably in the form of redox-sensitive, high molecular weight (HMW) smears (right *vs.* left panel; **Fig. 3a**). We saw the same pattern in fractions prepared from control midbrains; no such reactivity was seen in SDS-extracts of parkin*-*deficient ARPD brains, thus demonstrating detection specificity.

**Figure 3:**
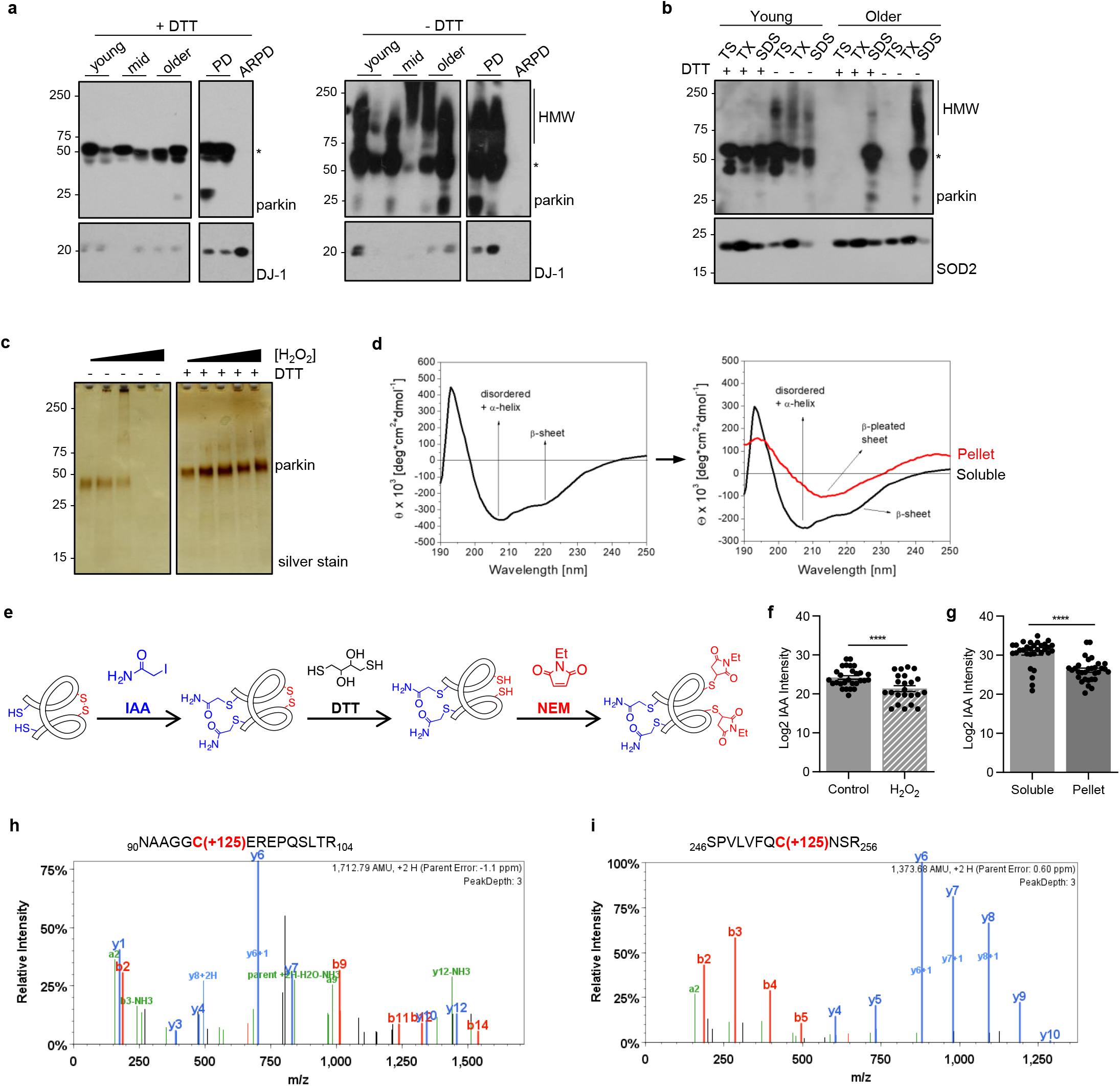
Parkin’s solubility and structure are altered by oxidative modifications. **(a)** Western blots of parkin and DJ-1 in SDS fractions from normal cortices (3 age groups are shown) and two age-matched patients, *i.e.,* idiopathic Parkinson’s (PD) and parkin-deficient ARPD. Sister aliquots of the same lysates were processed in parallel by SDS-PAGE either under reducing (+DTT) or non-reducing (-DTT) conditions. **(b)** Western blots of parkin and SOD2 distribution in serially fractionated human cortices from a young individual (age, 5 yrs) and an adult (62 yrs) subject, and separated by SDS-PAGE under reducing (+DTT) and non-reducing (-DTT) conditions. **(c)** Silver staining of the supernatant of sister aliquots of r-parkin following initial exposure to increasing concentrations of H_2_O_2_ (0-2mM) followed by the addition (or absence of) DTT (100mM) prior to centrifugation as indicated. **(d)** Circular dichroism spectra of soluble, untreated, wild-type r-parkin at the start of experiment (T=0; left panel), and spectra of soluble (black line) and aggregated (red line) states following incubation at 37°C for T=5 days (right panel). **(e)** Graphic depiction of strategy for LC-MS/MS-based analysis to identify cysteine oxidation state for untreated and H_2_O_2_-treated, parkin species, by using IAA-DTT-NEM fingerprinting to identify reduced cysteines with an iodacetamide (IAA) tag or reversibly-oxidized residues with a N-ethylmaleimide (NEM) tag. **(f-g)** Quantitative analyses of IAA-modified cysteines captured by LC-MS/MS for **(f)** untreated *vs.* H_2_O_2_-exposed, wild-type, human r-parkin, and **(g)** soluble compared to insoluble (pellet) fractions. Each dot represents the log2-transformed total IAA-signal intensities of individual cysteines (n=3 runs for each). The cysteine pool is shown with the mean ± SEM; significance **P<0.01, as determined using Student T-Test. **(h-i)** LC-MS/MS-generated spectra following trypsin digestion of labelled, oxidized r-parkin indicating NEM adducts (+125 mass gain) at Cys95 and Cys253; r-parkin was exposed to H_2_O_2_, and cysteines labelled as in (e). See **Extended Data Table 2** for a complete list of modified cysteines and oxidizing conditions.

We confirmed that reversible oxidation of brain parkin was also present in soluble (TS-, TX-) fractions, albeit at lesser intensities (**Fig. 3b**; data not shown). Of note, the formation of high *M_r_* parkin was not due to secondary oxidation *in vitro,* because specimens were processed and fractionated in the presence of iodoacetamide (IAA) prior to SDS/PAGE in order to protect unmodified thiols. These HMW parkin smears also did not arise from covalent ubiquitin-conjugation, such as due to auto-ubiquitylation of parkin, because such adducts cannot be reversed by reducing agents (*e.g.,* DTT).

Because we predicted that the loss in parkin solubility was due to thiol-based, posttranslational oxidation events [26], we sought to test this *in vitro* using purified, tag-less, full-length, recombinant (r-) parkin. There, we observed the H_2_O_2_ dose-dependent formation of HMW smears and loss of parkin solubility; however, protein solubility was recovered by adding DTT (**Fig. 3c**; **Supplementary Fig. 3a**) or β–mercaptoethanol (not shown). Demonstrating its sensitivity to bi-directional redox forces, the exposure of naïve r-parkin to excess DTT also rendered it increasingly insoluble (**Supplementary Fig. 3b**), likely due to loss of Zn^2+^ ion chelation at its four RING domains [25], which requires a zwitter-type redox state of their 28 cysteines [54]

Further, we also confirmed by mass spectrometry (MS; without any trypsin digestion of the holoprotein) that all 35 cysteine-based thiol groups of r-parkin are accessible to alkylation by IAA (right *vs.* left panel; **Supplementary Fig. 3c**). These results unequivocally demonstrated that each parkin cysteine theoretically possesses the capacity to function as a reducing thiol. Nevertheless, in these *in vitro* experiments we consistently observed a concentration-dependent change in r-parkin solubility, thereby suggesting that some thiols were more amenable than others to modification by reactive species (see below and **Supplementary Table 2**).

### Oxidative conditions alter parkin structure

The progressive insolubility of brain parkin and r-parkin due to redox stress suggested that the protein had undergone structural changes. Indeed, when we analyzed the effects of spontaneous oxidation using naïve r-parkin by far-UV-circular dichroism (**Fig. 3d**), soluble fractions initially contained both α-helically ordered as well as unstructured r-parkin proteins. Five days later, r-parkin preparations were separated by centrifugation and fractions re-analyzed. There, we found a marked shift to increased β-pleated sheet-positive r-parkin in insoluble fractions (**Fig. 3d**). Similarly, when we monitored r-parkin during spontaneous oxidization using dynamic-light scattering (**Supplementary Fig. 3d**), we observed a gradual shift in the hydrodynamic diameter from 5.1 nm, representing a folded monomer, to multiple peaks with larger diameters 5 hrs later. The latter indicated spontaneous multimer formation, which was partially reversed by the addition of DTT (right panel; **Supplementary Fig. 3d**). Thus, these structural and solubility changes of r-parkin were congruent with our immunoblot results for human brain parkin (**Fig. 3a**).

### Hydrogen peroxide modifies parkin at multiple cysteines

To determine whether oxidation of cysteines and/or methionine residues caused parkin insolubility, we analysed r-parkin that was treated with and without H_2_O_2_ and/or thiol-alkylating agents using liquid chromatography-based MS (LC-MS/MS). To differentiate reduced from oxidized cysteines we used a serial thiol-fingerprinting approach (**Fig. 3e**), which labelled reduced thiols with IAA; it and the tagging of reversibly oxidized thiols with N-ethylmaleimide (NEM) after their prior reduction with DTT (**Fig. 3e**). The first test was to determine how progressive oxidation affected thiol-accessibility. As expected, using the strong alkylating agent IAA on the nascent protein, we found that the majority of parkin cysteines were reactive (**Supplementary Fig. 3c; Supplementary Table 2**).

However, when treating naïve r-parkin with lower H_2_O_2_ concentrations, we identified an average of 19 cysteines (54.3%); in contrast, higher H_2_O_2_ concentrations increased this number to 32 cysteines (91.4%). These results suggested progressive protein unfolding with increasing oxidation (**Supplementary Table 2**).

Next, we sought to more precisely identify the number and pinpoint the location of oxidized cysteine residues. Using Scaffold PTM-software, we found a rise in the number of oxidized residues (NEM-Cys, range of 3-26), which was proportional to the increase in H_2_O_2_ concentrations and appeared to begin in the RING1 domain at three residues, *i.e.,* Cys238, Cys241 and Cys253 (**Supplementary Table 2; Fig. 3i**), but also involved Cys95 in the linker domain (**Fig. 3h**). Furthermore, when quantifying thiol modifications by MaxQuant software [47], we found a significant drop for the number of cysteines in the reduced state (IAA-cysteines) within the H_2_O_2_-treated samples (P=0.0016; **Fig. 3f**), as expected.

In accordance, when comparing cysteine oxidation events in soluble and insoluble fractions of untreated *vs.* oxidized r-parkin preparations, the number of IAA-Cys was significantly decreased in the pellets (P<0.0001; **Fig. 3g**). Of note, modifications at methionine residues did not correlate with r-parkin solubility. These collective results unequivocally demonstrated that H_2_O_2_-induced oxidation of cysteine-based thiols is linked to both progressive, structural change and the insolubility of human r-parkin.

### Parkin is also irreversibly oxidized in adult human and mouse brain

We next sought to confirm oxidation of parkin cysteine residues *in vivo* by LC-MS/MS. To this end, we examined both human cortex-derived parkin and parkin isolated from intraperitoneally, MPTP toxin-(*vs.* saline-) treated murine brains (**Fig. 4**). Specimens were processed with IAA during homogenization and fractionation to prevent any oxidation artefacts *post mortem*. Following immunoprecipitation and gel excision of endogenous parkin at the 50-53 kDa range (an example is shown in **Supplementary Fig. 4a,b**), we focused on cysteine mapping and the identification of thiol redox states (**Fig. 4a**). A graphic representation of theoretically possible, thiol-based redox modifications is provided in **Supplementary Fig. 4c**).

**Figure 4:**
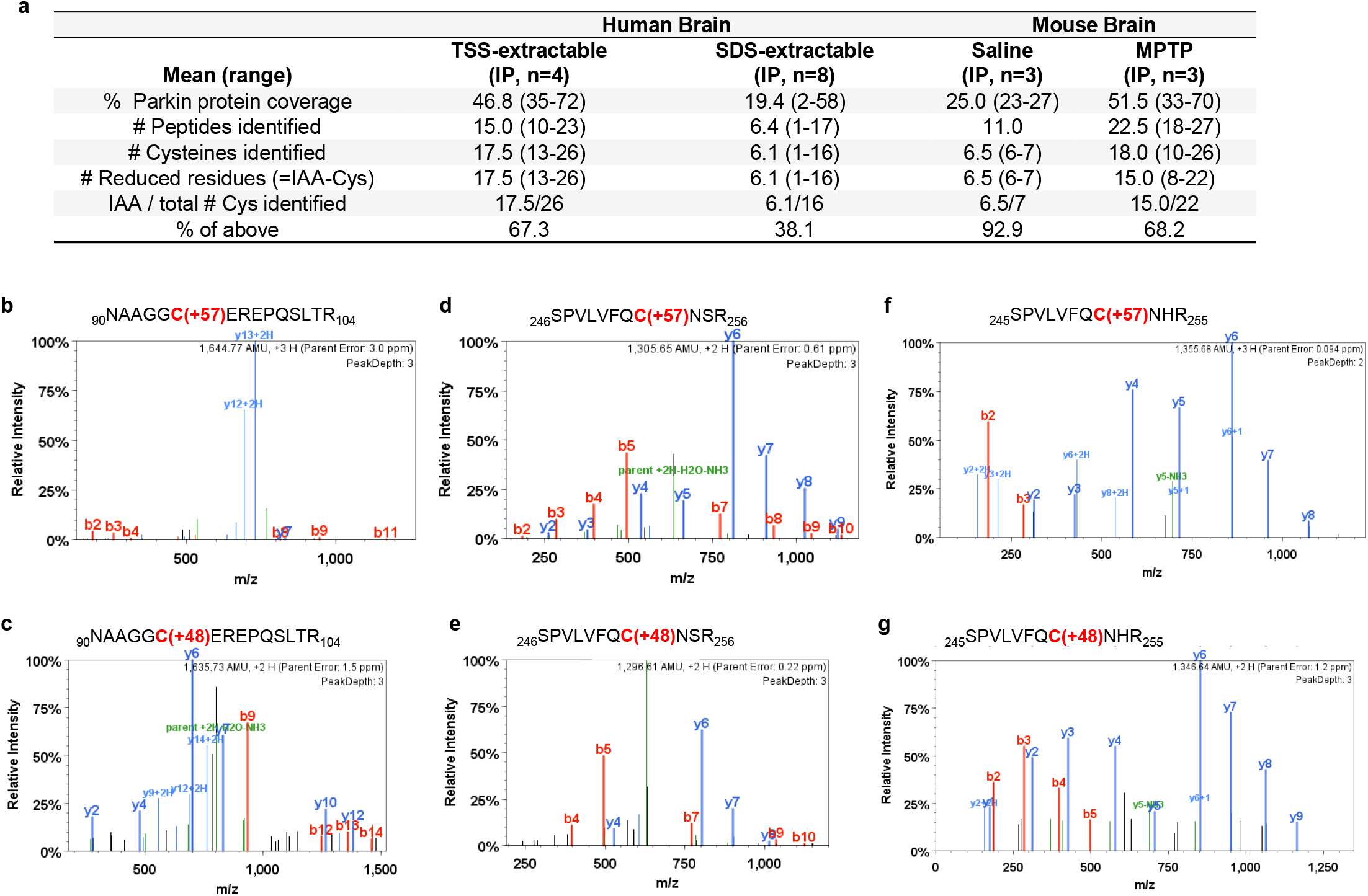
Parkin cysteine residues are oxidized in human and mouse brain. **(a)** Summary of results for 12 immunoprecipitation (IP) runs (TS extracts; n=4; SDS extracts, n=8) from human cortices and either saline- or acute (1hr) MPTP toxin-treated murine brain (as described in Fig. 2d,e) for endogenous parkin enrichment to identify the redox state of its cysteine residues (see also b-g). All specimens were fractionated in the presence of IAA. **(b-g)** Among the redox active residues identified, Cys95 and Cys253 in human brain parkin were found in either a reduced redox state **(b,d)** (*i.e.,* IAA-labelled; +57 mass gain), or **(c,e)** in irreversibly oxidized states, *e.g.,* to sulfonic acid (trioxidation; +48 mass). In mouse brain parkin **(f,g)**, Cys252 was found either reduced or oxidized as well.

In human control cortices (n=12 runs; summarized in **Fig 4a**), we mapped a mean of 46.8 and 19.4% of parkin wild-type sequence in the soluble and insoluble fractions, respectively. There, we found cysteines in either a redox reduced state (IAA-alkylated Cys+57; examples shown in **Fig. 4b,d**) or in oxidized states (*e.g.,* to sulfonic acid Cys+48). Irreversible oxidation events in human cortex occurred, for example, at Cys95 (**Fig. 4c**) and Cys253 **(Fig. 4e**). The relative frequencies of detection for parkin thiols that were reduced *in vivo* (and alkylated by IAA *in vitro*) in the soluble *vs.* insoluble fractions of human brain were 67.3 and 38.1%, respectively (**Fig. 4a**).

Likewise, in saline- and MPTP-treated mouse brains (n=6 runs), we mapped 25 and 51.5 per cent of parkin, respectively (summarized in **Fig. 4a**). Interestingly, like in the human studies, in these runs we identified the murine-corresponding residue Cys252 in either a reduced or irreversibly oxidized states (**Fig. 4f,g**). As mentioned, mice do not carry a cysteine at residue 95 (for sequence comparison, see below). The relative frequencies of detection for thiols that were reduced *in vivo* (and alkylated by IAA *in vitro*) in parkin from saline-*vs.* MPTP toxin-treated mouse brains were 92.9 and 68.2%, respectively (**Fig. 4a**). We concluded from these analyses that the decline in the relative number of reduced thiols in less soluble fractions of mammalian brain reflected a greater degree of oxidative, posttranslational modifications of wild-type parkin.

### Parkin thiols reduce hydrogen peroxide

A typical redox reaction involves the reduction of an oxidized molecule in exchange for oxidation of the reducing agent that occurs in parallel (**Supplementary Fig. 4c**). We asked whether parkin oxidation resulted in reciprocal reduction of its environment, *i.e*., anti-oxidant activity (**Fig. 5; Supplementary Fig. 5**). Using r-parkin, we confirmed that parkin could directly lower H_2_O_2_ levels in a concentration-dependent manner *in vitro* (**Fig. 5a; Supplementary Fig. 5h**). This reducing activity was not enzymatic, in that it did not mirror the dynamics of catalase, and r-parkin did not possess peroxidase activity (**Fig. 5a; Supplementary Fig. 5a**). Rather, the reaction was dependent on thiol integrity, because pre-treatment with NEM (or IAA) and pre-oxidation of the protein with H_2_O_2_ abrogated the ROS-reducing activity of r-parkin (**Fig. 5b; Supplementary Fig. 5b,g**).

**Figure 5:**
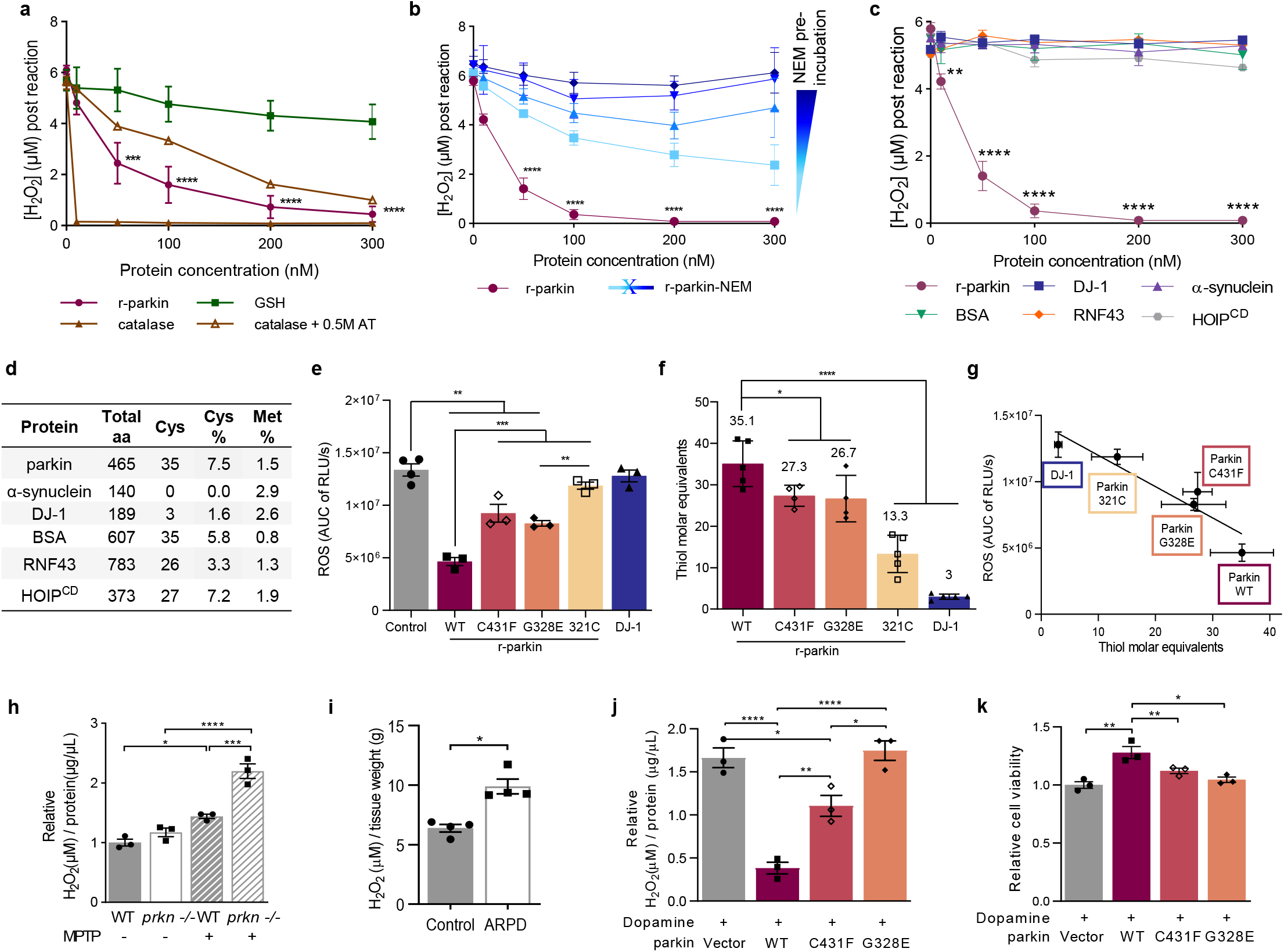
Wild-type parkin lowers hydrogen peroxide *in vitro*, in cells and the brain. (**a-c**) Quantification of H_2_O_2_ concentration using AmplexRed, demonstrating **(a)** full-length, human, recombinant (r-) parkin when incubated with H_2_O_2_ is able to reduce it to water in a r-parkin concentration-dependent manner. Effects of r-Parkin were compared to catalase and GSH at equimolar concentrations as well as following partial inhibition of catalase by amino-triazole (AT), as indicated. **(b)** Pre-incubation of r-parkin with a thiol-conjugating compound (NEM) inhibits parkin-dependent H_2_O_2_ reduction in a NEM-concentration-dependent manner. **(c)** Reducing capacity of wild-type r-parkin compared to two other, PD-linked proteins (DJ-1; α-synuclein), bovine serum albumin (BSA) and two RING-carrying ubiquitin ligases (RNF43; HOIP^cd^; cd = catalytic domain). Their respective cysteine and methionine contents are summarized in **(d)**. Two-way ANOVA was used for statistical analysis (*p < 0.05, **p < 0.01,***p < 0.001, and ****p < 0.0001) **(e)** Area under the curve (AUC) plots for results from *in vitro* colorimetric assays, where AUC integrates total H_2_O_2_ levels measured over the time course of the assay (see also **Extended Data Fig. 5e**). Comparison of WT r-parkin with DJ-1, two r-parkin point mutants, and r-parkin_321-465_ (321C). Results represent n=3 ± SD; *p < 0.05, **p < 0.01,***p < 0.001, and ****p < 0.0001 using one-way ANOVA with Tukey’s post hoc test. **(f)** Quantification of reactive thiol content (in molar equivalents) for r-parkin (WT; two point mutants; 321C) and full-length r-DJ-1 using the Ellman’s reagent assay. **(g)** Correlation curve between number of free thiols **(f*) vs***. the H_2_O_2_-reducing capacity **(e)** for indicated proteins. **(h-i)** Quantification of H_2_O_2_ levels in **(h)** saline *vs.* MPTP toxin-treated *prkn* wild-type (WT) and *prkn*^−/−^ mouse brain (n=3/genotype/condition), and **(i)** in human brain from parkin-deficient ARPD cortices compared to age- and *post-mortem* interval-matched controls (n=4/group) collected at the same institution. Results are represented as the mean concentration of H_2_O_2_ (μM) per total protein concentration (μg/μL) or tissue weight (g) analyzed ± SEM; *P<0.05, ***p < 0.001, and ****p < 0.0001 determined using a Student T-test or one-way ANOVA. **(j-k)** H_2_O_2_ quantification **(j)** and cell viability assay **(k)** for dopamine-treated, human M17 cells expressing either WT or two ARPD-linked parkin point mutants, as indicated relative to treatment with vehicle alone. Cells were exposed to 200 mM dopamine or vehicle for 20h, as indicated. Data points represent the mean of duplicates ± SEM (n=3 experiments); *P<0.05 and **p < 0.01, and ****p < 0.0001 by one-way or two-way ANOVA.

This effect by r-parkin was also dependent on its intact Zn^2+^ coordination (**Supplementary Fig. 5c**). Interestingly, RNF43 (another E3 ligase that contains a zinc-finger domain), HOIP (an E3 ligase containing a RING domain) and bovine serum albumin (BSA, which akin to parkin has 35 cysteines), did not show any H_2_O_2_-lowering capacity (**Fig. 5c,d; Supplementary Fig. 5d**). Further, PD-linked α-synuclein, which has no cysteines, also had no reducing effect (**Fig. 5c,d**). These results suggested that the cysteine-rich, primary sequence and the tertiary structure of r-parkin can confer anti-oxidant activity.

We next examined additional cysteine-containing, PD-linked proteins, *e.g.*, r-DJ-1, a C-terminal RING2-peptide of parkin (r-parkin_321C_), and two disease-linked variants of full-length r-parkin, p.G328E and p.C431F. We also used a second ROS quantification assay for further validation and to examine dose dependency (**Fig. 5e, Supplementary Fig. 5e-l**). There, r-DJ-1 and r-parkin_321C_ showed negligible H_2_O_2_-lowering capacity, and the two point-mutants conferred less activity than did wild-type, human r-parkin (**Fig. 5e**). As expected (**Supplementary Fig. 4c**), the lowering of ROS correlated with reciprocal r-parkin oxidation, as revealed by SDS/PAGE, which was performed under non-reducing conditions immediately after the reaction (**Supplementary Fig. 5m**).

These results suggested that anti-oxidant activity by parkin was dependent on its reactive thiol content, which we examined next using the Ellman’s reagent. There, wild-type r-parkin, r-parkin_321C_ (that contains two, non-RING-based cysteines) and r-DJ-1 showed the predicted number of reactive thiols, whereas the single point-mutant variants of r-parkin revealed fewer accessible thiols (**Fig. 5f**). From these results, we were able to calculate a linear correlation between thiol equivalencies and the degree of ROS reduction, demonstrating that a greater number of reactive and/or a greater number of accessible thiols in parkin proteins correspond well with more effective lowering of H_2_O_2_ (**Fig. 5g**).

### Hydrogen peroxide levels are increased in parkin-deficient brain

To explore whether parkin oxidation conferred ROS reduction *in vivo*, we first quantified H_2_O_2_ concentrations in the brains of wild-type and *prkn*^−/−^ mice. A trend, but no significant difference, was observed under normal redox equilibrium conditions. However, when analyzing brain homogenates from mice treated with MPTP toxin *vs.* saline, as described above (Fig. 2), we found significantly higher H_2_O_2_ levels in the brains of adult *prkn*^−/−^ mice compared to wild-type littermates (P<0.001; **Fig. 5h**). Similarly, in adult humans H_2_O_2_ levels were significantly increased in the cortex of *PRKN-*linked ARPD patients *vs.* age-, *post mortem* interval-, ethnicity- and brain region-matched controls [1] (P<0.05; **Fig. 5i**). Specimens of three non*-PRKN*-linked cases with parkinsonism showed H_2_O_2_ levels comparable to those from age-matched normal cortices (Fig. 2b, red circles). We concluded that the expression of wild-type *PRKN* contributes to the lowering of ROS concentrations in adult, mammalian brain.

### Parkin prevents dopamine toxicity in part by lowering hydrogen peroxide

To address the question of selective neuroprotection, we revisited the role of parkin in cellular dopamine toxicity studies [34, 55]. We first tested its effect on ROS concentrations in dopamine-synthesizing, human M17 neuroblastoma cells. There, dopamine exposure of up to 24 hrs caused a significant rise in endogenous H_2_O_2_ (P<0.05; **Fig. 5j**), as expected. Wild-type parkin expression effectively protected M17 cells against the dopamine stress-related rise in H_2_O_2_ levels (P<0.0001; **Fig. 5j)**. By comparing sister cultures that expressed similar amounts of exogenous parkin proteins, the E3 ligase-inactive p.C431F mutant had a partial rescue effect, whereas p.G328E, which we confirmed to retain its E3 ligase activity *in vitro*, showed no H_2_O_2_-lowering capacity in cells (**Fig. 5j**; and data not shown).

Under these conditions, only wild-type parkin, but none of the mutant variants we tested, increased M17 cell viability under rising dopamine stress conditions (P<0.01; **Fig. 5k**; and data not shown). This protective effect also correlated with parkin insolubility and HMW smear formation, as expected from previous studies [34]. These posttranslational changes in M17-expressed parkin were not reversible by DTT or SDS (**Supplementary Fig. 6a,b**), thereby suggesting irreversible dopamine-adduct formation. Notably, the protection from dopamine toxicity positively correlated with the level of *PRKN* cDNA transcribed, as confirmed in sister lines of M17 cells that stably express human parkin. There, we estimated that ~4 ng of parkin protein expressed in healthy, neural cultures neutralized each μM of dopamine added during up to 24 hrs (**Supplementary Fig. 6c,d**).

### Parkin binds dopamine radicals predominantly at primate-specific cysteine 95

We next explored which thiols of parkin were relevant for the neutralization of dopamine radicals. Covalent conjugation of RES metabolites at parkin residues had been previously suggested [34, 55], but not yet mapped by LC-MS/MS examining the whole protein. Aliquots of r-parkin were exposed to increasing levels of the relatively stable dopamine metabolite aminochrome. As expected, this led to the loss of protein solubility and HMW species formation at the highest dose tested (**Fig. 6a,b**). These reaction products were then used to map modified residues by LC-MS/MS. Specifically, proteins corresponding to r-parkin monomer (51-53 kDa) and two HMW bands, one at ~100 kDa, the other near the loading well, were gel-excised (**Fig. 6a**), trypsin digested and further analyzed.

**Figure 6:**
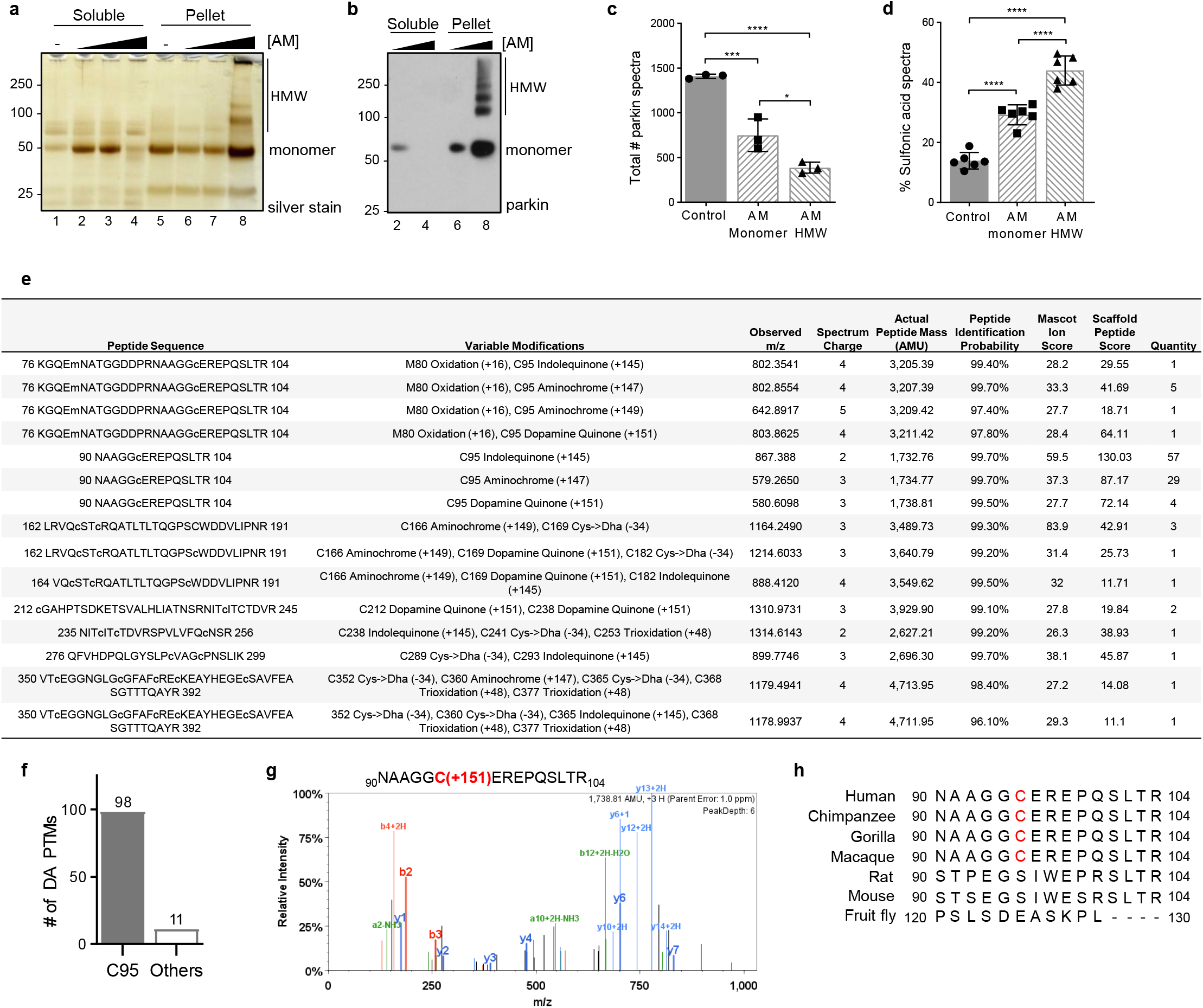
Human parkin conjugates dopamine radicals foremost at residue Cys95. **(a-b)** Silver staining **(a)** and Western blot **(b)** of r-parkin in soluble (supernatant) and insoluble (pellet) phases following exposure to increasing concentrations of aminochrome (AM; 0-200 μM) and analyzed under non-reducing conditions. See lane number for corresponding samples. **(c)** Mean total number of parkin spectra, as identified by LC-MS/MS following trypsin digestion, of control *vs.* monomeric *vs.* high molecular weight (HMW), AM-modified r-parkin. Data represent the mean of n=3 runs ± SEM. *P<0.05; ***P<0.001; ****P<0.0001 by 1-way ANOVA. **(d)** Percentage of peptides carrying a sulfonic acid modification in control *vs.* monomeric and HMW, AM-modified r-parkin. Each point represents one gel specimen submitted to MS. The percentage was calculated using only the subset of peptides that were ever detected as carrying a sulfonic acid modification. Statistics were done as in (c). **(e)** Table summarizing LC-MS/MS-based detection of adducts representing dopamine metabolites conjugated to cysteines identified in human r-parkin following exposure to aminochrome *in vitro*. Chemical structures for identified cysteine-conjugated adducts are shown in **Extended Data Fig. 7b**. Individual quantification of each peptide with adduct listed is shown on the right side of the table. **(f)** Frequency of occurrences for dopamine-metabolite adducts being detected on Cys95 *vs.* all other cysteine residues, as detected by LC-MS/MS and individually shown in (e). **(g)** LC-MS/MS-generated spectrum following trypsin digestion of AM-exposed r-parkin highlighting a dopamine (+151 mass gain) adduct covalently bound to Cys95. See also **Extended Data Fig. 7c-p** for additional spectra. **(h)** Species comparison for wild-type parkin proteins covering sequence alignment of aa90-104, with primate-specific residue Cys95 highlighted in red.

There, we made the following four related observations: i) Increasing aminochrome concentrations led to a significant decline in the total number of spectra readily identified as parkin-derived peptides, both in the monomeric and HMW bands (P<0.001 and P<0.0001), respectively (**Fig. 6c**). This indicated to us either a marked loss in solubility or a rise in heterogenous, complex modifications, which rendered the analyte undetectable by LC-MS/MS, or both; ii) Despite fewer spectra recorded, we identified a significant increase in oxidized cysteines (*e.g.*, irreversibly to sulfonic acid) following aminochrome exposure, in particular within the HMW bands of r-parkin (P<0.0001; **Fig. 6d**); iii) Under these conditions, four distinct forms of dopamine metabolites were found conjugated to parkin cysteines. Mass shifts of +145, +147, +149 and +151 were found, which represented conjugation to indole-5,6-quinone, two variants of aminochrome (O=; HO-), and dopamine quinone itself, respectively (**Fig. 6e; Supplementary Fig. 7a**); and iv) Unexpectedly, we identified Cys95 to be the most frequently dopamine-conjugated parkin residue (P<0.0001; n=98 spectra; **Fig. 6e-g; Supplementary Fig. 7b-g**). Other residues of r-parkin identified carrying dopamine metabolite adducts included Cys166, Cys169, Cys212, Cys238, Cys360, Cys365 and Met80 (all together, n=11 spectra; **Fig. 6e; Supplementary Fig. 7h-o**). No dopamine metabolite-related mass shifts were detected in control samples that had not been exposed to aminochrome, as expected.

### Parkin augments melanin formation *in vitro*, which requires primate-specific Cys95

Given the observed relations between r-parkin, dopamine radical conjugation, aggregate formation and protein insolubility, we next examined whether melanin formation was altered in the presence of parkin. Oxidation of dopamine, in the presence of proteins containing cysteine groups, generates covalent adduct-carrying proteins that share important structural characteristics of neuromelanin pigment in the human midbrain. Additionally, such synthetic versions behave similarly to human neuromelanin in cell cultures [56, 57]. Unexpectedly, we discovered that wild-type r-parkin augmented total melanin formation in a protein concentration- and time-dependent manner (**Fig. 7a**). Like the wild-type protein, two ARPD-linked, full-length r-parkin variants, p.C431F and p.G328E, also augmented melanin formation *in vitro*, when monitored over 60 mins, whereas r-DJ-1 and BSA had no such effect in this assay (**Fig. 7b**).

**Figure 7.**
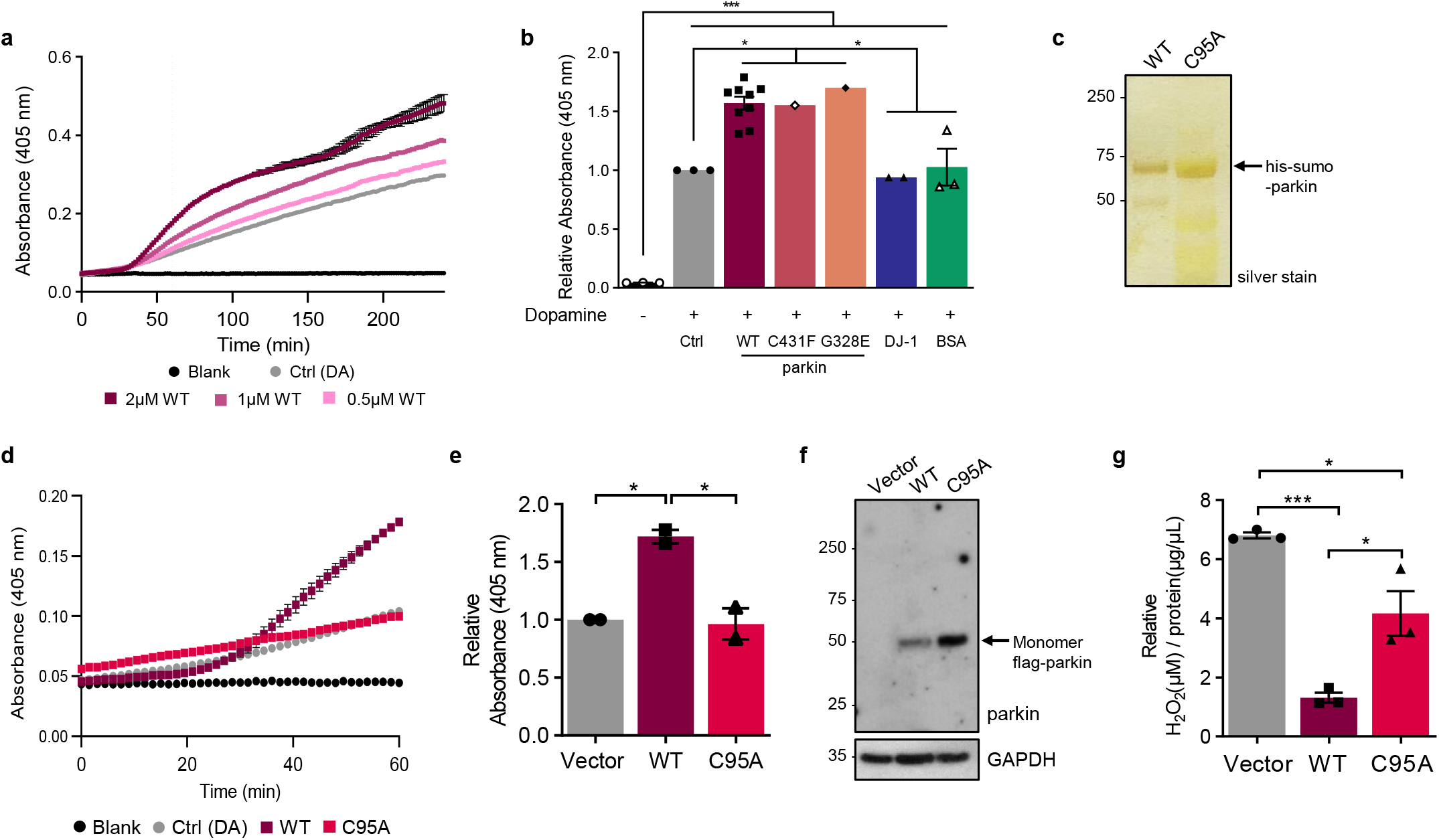
Parkin-dependent increase in melanin formation requires cysteine 95. **(a)** Kinetic curve of melanin production (read at absorbance 405nm) over time in the absence of exogenous protein (dopamine (DA Ctrl) alone) *vs.* increasing molar concentrations of wild-type (WT), full-length human r-parkin shown for three concentrations (0.5, 1, 2mm). Each condition was performed in triplicate. **(b)** Total melanin formation for indicated recombinant proteins at 60 mins, as expressed relative to its production under dopamine only control (Ctrl) condition. Data represent the mean of triplicates ± SEM. ***P<0.05 by 1-way ANOVA. **(c)** Silver gel for the analysis of His-SUMO-tagged, full-length, human r-parkin proteins of wild-type sequence and its variant carrying a p.C95A mutation. **(d-e)** Representative kinetic curve for melanin production **(d)** and relative total melanin formation at 60 mins **(e)**, where production in the presence of wild-type (WT) or p.C95A mutant r-parkin (each, 2 mM) is shown relative to dopamine (DA) (Ctrl) alone. Data represent mean of n=2, each performed in triplicate ± SEM. ***P<0.05 by 1-way ANOVA. **(f-g)** Protein expression, as shown by Western blotting **(f)**, and fold change in H_2_O_2_ levels **(g)** for dopamine-treated M17 cells-relative to vehicle treated sister wells-that transiently express either flag-vector, or WT *vs.* p.C95A-mutant human parkin-encoding cDNA plasmids. Results are shown as mean ± SEM (n=3) and all dopamine-treated samples (200mM dopamine) were normalized to their respective untreated samples. Anti-GAPDH immunoblotting served as loading control (in f). A one-way ANOVA was used for statistical analysis (*P<0.05 and ***P<0.001).

Interestingly, mutating residue Cys95 to alanine (p.C95A; **Fig. 7c**) completely abrogated the enhancing effect by r-parkin on the polymerization of melanin (**Fig. 7d,e**). Of note, in our study all recombinant proteins heretofore analyzed were used after their N-terminal His-SUMO-tag had been removed; however, the p.C95A-mutant was resistant to enzymatic digestion of the tag from the holoprotein. Therefore, both His-SUMO-r-parkin and His-SUMO p.C95A were utilized (**Fig. 7c-e**). Importantly, we saw no difference in the kinetics of melanin formation between wild-type r-parkin proteins that either carried a His-SUMO*-*tag or were tag-less (not shown).

Furthermore, when testing p.C95A-mutant parkin in the M17 cell-based dopamine toxicity assay, the variant showed only a partial effect in H_2_O_2_ lowering when compared to wild-type parkin, even when p.C95A was expressed at higher levels (**Fig. 7f,g**). These results were consistent with our collective LC-MS/MS results of oxidative modifications of parkin at Cys95 (shown in: **Figs. 3h; 4c; Supplementary Table 2**). We reasoned from these collective *ex vivo* results that wild-type parkin could be associated with the synthesis of neuromelanin *in vivo*. Therefore, we sought to explore this further in dopamine neurons of human midbrain.

### Anti-parkin reactivity localizes to neuromelanin in *S. nigra* of adult control brain

Subcellular localization studies of parkin in adult human control brain had previously been hindered by the lack of renewable antibodies (Abs) that reliably detect the protein *in situ* [49, 52, 58, 59]. We therefore developed and extensively validated several, epitope-mapped, monoclonal Abs of the IgG_2_b-subtype using preparations of untagged full-length, human r-parkin as immunogen. To this end, we generated four stable clones, *i.e.,* A15165<u>B</u>, A15165<u>D</u>, A15165<u>G</u>, and A15165<u>E</u> (**Supplementary Fig. 8a-c;** Tokarew et al., *in preparation*), which were applied to microscopy studies.

Serial sections of control adult, human midbrains were developed by traditional immunohistochemistry (IHC) using enhanced 3-3’-diaminobenzadine (eDAB) generating a black signal. There, anti-parkin clones A15165<u>D</u>, A15165<u>G</u> and A15165<u>E</u> revealed dark, granular staining throughout the cytoplasm of pigmented cells (ages, > 55 yrs) (**Fig. 8a,b,d**). Using sections of anterior midbrains from nine adult control subjects, >83% of the anti-tyrosine hydroxylase (TH)-positive neurons were also positive for parkin, as quantified by double labelling (**Fig. 8c**). Under these conditions and Ab concentrations, no anti-parkin signal was detected in glial cells.

Intriguingly, sections from younger control subjects (ages, <33 yrs) that were processed in parallel revealed less intense, anti-parkin reactivity in *S. nigra* neurons, which matched the paucity of their intracellular pigment (**Fig. 8e**); of note, mature neuromelanin consistently generates a brown color in sections developed without any primary Ab. The difference between younger *vs.* older midbrains suggested that the three anti-parkin clones likely reacted with an age-related, modified form of parkin *in situ,* because the *PRKN* gene is already expressed in dopamine cells at a young age (**Fig. 1b; Supplementary Fig. 1a-d**).

To confirm the specificity of the new anti-parkin clones, we serially stained midbrain sections from a 71 yr-old, male ARPD patient, who was entirely deficient in parkin protein due to compound heterozygous deletions of *PRKN* exons 2 and 3 (**Fig. 8f; Supplementary Fig. 9a-c**). Development of serial sections with anti-parkin clones A15165<u>E</u>, -<u>D</u> and -<u>G</u> revealed no immunoreactivity in surviving midbrain neurons of the *S. nigra* from this ARPD case. In the absence of parkin, there was no signal overlap between eDAB reactivity (black color) and either intracellular neuromelanin granules in surviving dopamine cells or with extracellular pigment (brown; **Fig. 8f; Supplementary Fig. 9c**). In parallel, development of midbrain sections from individuals with the diagnoses of dementia with Lewy bodies, of non-*PRKN-*linked, sporadic PD as well as of cases with incidental Lewy bodies readily demonstrated eDAB reactivity overlapping with neuromelanin for all three anti-parkin clones (**Supplementary Fig. 9d-g**; and data not shown). These results demonstrated specific staining by the three anti-parkin clones in our microscopy studies of *post mortem* human brain.

**Figure 8.**
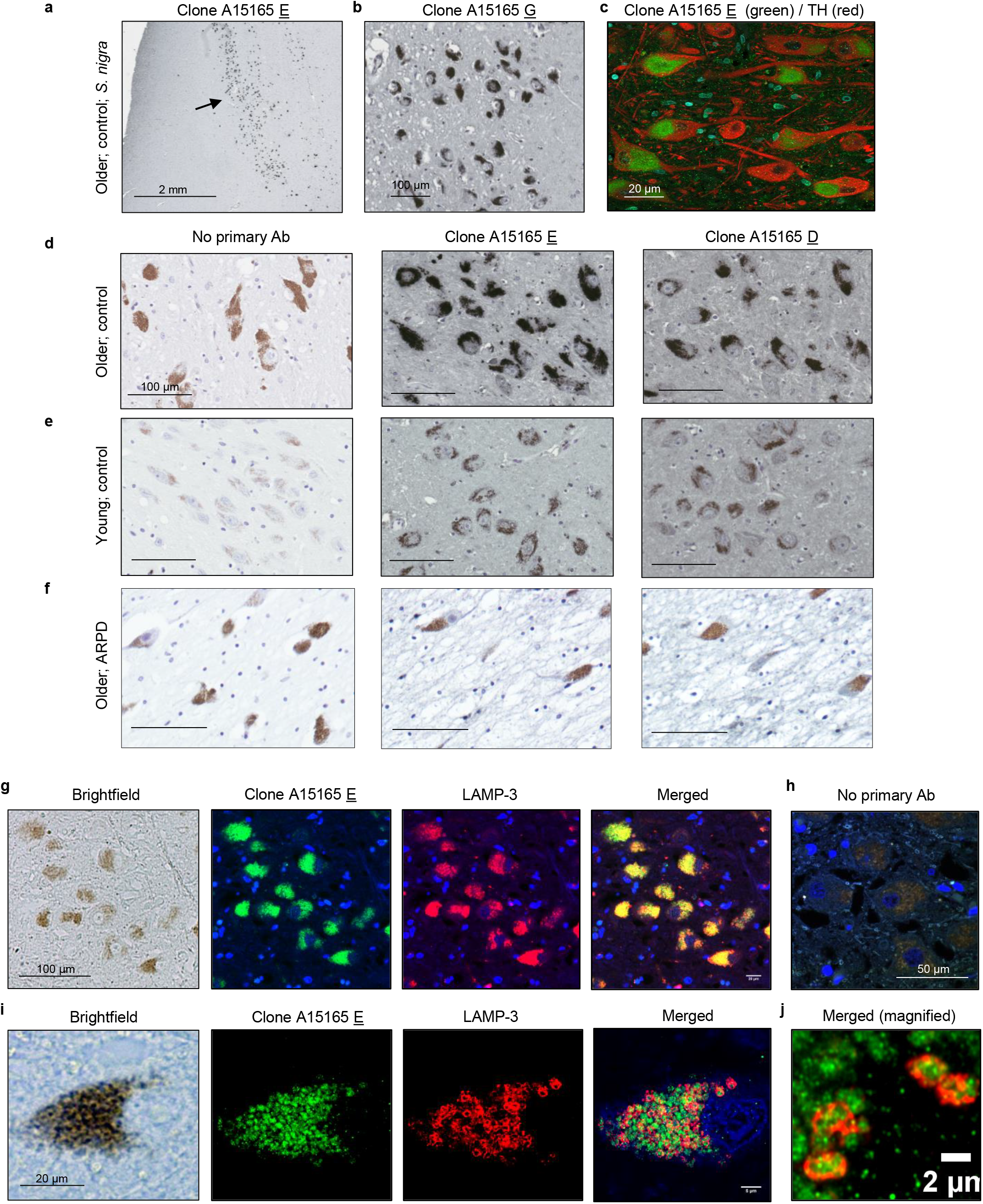
Parkin localizes to neuromelanin pigment in *S. nigra* neurons of normal human midbrain. **(a-b)** Immunohistochemical detection of parkin in adult human brain including dopamine neurons of the *S. nigra* using anti-parkin monoclonal antibody clones A15165 E **(a)** and -G (**b)**. **(c)** Double labelling for tyrosine hydroxylase (TH) and parkin (clone A15165 E) in the *S. nigra* from an adult control subject using indirect immunofluorescence microscopy. **(d-f)** Immunohistochemical reactivities generated by no primary antibody *vs.* two anti-parkin (Clones A15165 E, D) antibodies on sections of the *S. nigra* from two control subjects, aged **(d)** 66 yrs and **(e)** 24 yrs, as well as **(f)** from a parkin-deficient ARPD case, aged 71 years. In the indicated panels, immunoreactivity was detected by metal-enhanced DAB (eDAB; generating black colour) and hematoxyline as a counterstain (blue). No primary antibody added generates a pigment-induced signal for neuromelanin (brown). Scale bars represent 100 mm, or as indicated. **(g-j)** Immunofluorescent signals, as generated by double-labelling of human *S. nigra* sections containing dopamine neurons, using anti-parkin (clone-E; green colour) and anti-LAMP-3/CD63 (red colour) antibodies; (blue colour, Hoechst stain). Brightfield microscopy image in the same field (neuromelanin pigment is visible; left panel) and a no primary antibody **(h)** run in parallel are shown. **(i)** Higher magnification of a single dopamine neuron and **(j)** further magnification for visualization of subcellular signals within a neighbouring dopamine neuron is shown, as indicate**Supplementary Figure Legends**

### Parkin frequently localizes to LAMP-3^+^-lysosomes within *S. nigra* neurons

Neuromelanin granules have been shown to occur in specialized autolysosomes[60]. When screening for co-localization of parkin reactivity with markers of subcellular organelles in sections of adult control brain, we detected that immunofluorescent signals by anti-parkin (green) and anti-CD63/LAMP-3 (red) antibodies strongly overlapped with pigmented granules of nigral neurons (**Fig. 8g-i**; see also **Supplementary Fig. 9h**).

Using confocal microscopy, we demonstrated that in adult midbrain anti-parkin signals, as generated by clone A15165<u>E</u>, and neuromelanin granules were frequently surrounded by circular, ~2 μM (diameter)-sized rings of anti-LAMP-3 reactivity (**Fig. 8i,j**). A z-stack video for the parkin and LAMP-3 co-labelling studies is appended (**Supplemental Information**). We concluded that in the adult, human midbrain from neurologically healthy controls and in subjects, who suffer from parkinsonism that is not linked to bi-allelic *PRKN* deletion, a pool of parkin appears physically associated with neuromelanin pigment in close association with lysosomal structures.

## Discussion

Here, we demonstrate that the state of parkin’s cysteines is linked to its age-related insolubility and redox homeostasis in human brain. Our study provides first insights into the metabolism of wild-type parkin in adult midbrain, where we discovered -and quantified-that >90% of detectable parkin is insoluble. The loss in parkin solubility in the brain is unique, when compared to other PD-linked proteins and mitochondrial constituents tested. It is also tissue and species-specific. Approximately 50% of parkin remained soluble in spinal cord and skeletal muscle from aged human donors, and its insolubility was not observed in aged rodent and adult monkey brain (**Fig. 1, Supplementary Figs. 1,2**).

In human brain, the loss of parkin solubility correlated with a rise in H_2_O_2_ concentrations and with age. The transition to insolubility in the cortex occurs between 28 and 42 yrs (**Figs. 1,2,4; Supplementary Table 1**); the age at which parkin transitions in the *S. nigra* will require a larger number of midbrain specimens from young, neurologically normal subjects. While we were unable to assess solubility in such midbrains (<20 yrs), parkin’s distribution in adult midbrain was the same as in cortices. In brainstem nuclei, parkin partitioning across fractions was not affected by disease state (controls (n=11) *vs.* neuropathological cases (n=9); **Fig. 1b**; **Supplementary Table 1**), although its total abundance was lower in the *S. nigra* of cases from subjects with neurodegenerative parkinsonism, as expected (not shown).

We demonstrate that the observed loss of parkin solubility occurs via thiol-oxidation and that these post-translational modifications are linked to three protective outcomes: i) the neutralization of otherwise toxic reactive species (ROS, RES); ii) the net reduction of H_2_O_2_; and iii) the strong possibility that parkin has a role in dopamine metabolism through enhanced, Cys95-linked melanin formation. We have modeled parkin’s redox chemistry-based function *in vitro*, in cells and in mice, and provide evidence that these outcomes are physiologically relevant to human brain. From these observations we propose that insoluble parkin proteins represent functionally important metabolites of the ageing human brain including those of the *S. nigra*. Further, our findings integrate the early literature related to parkin mutations and stress-induced modifications *vis a vis* its insolubility and aggregate formation, which included a wide range of complementary investigations [27, 28, 33, 55, 61-66], such as findings from induced pluripotent stem cell-derived, human dopamine neurons [67–69]. Our discovery of parkin’s function in redox homeostasis also helps explain seemingly disparate evidence of previous observations made in flies, mice [19, 20] and humans [52].

The reactivity of cysteines is governed by their redox state (oxidized *vs.* reduced). It is influenced by the surrounding electrostatic environment, including via the charges of neighbouring residues [70]. Unlike parkin, 34 out of 35 cysteines found in BSA are engaged in disulphide-bond formation [71, 72]; it follows, that BSA was not able to reduce H_2_O_2_ *in vitro*, nor did it enhance the formation of insoluble melanin *in vitro*. Two other Zn^2+^-coordinating, cysteine-containing proteins that we tested, RNF43 and HOIP^CD^ (**Fig. 5c**), also failed to lower H_2_O_2_, thus suggesting that select cysteines in parkin have a higher affinity for ROS and, as discussed below, RES molecules. When mapping the redox state of parkin cysteines under progressively pro-oxidant conditions, we found that Zn^2+^-coordinating residues are not protected from ROS modification[73] (**Supplementary Table 2**).

In our experiments, we also estimated the levels of pro-*vs.* anti-oxidant forces. There, the ratio of H_2_O_2_-to-r-parkin (0.1-1 mM of H_2_O_2_ : 1 ng of r-parkin) was within the physiological range of what we calculated for human control brain extracts (*i.e.,* 0.4-6 mM of H_2_O_2_ : 1 ng of parkin). In human brain extracts, H_2_O_2_ concentrations were calculated to lie between 700 and 9,100 μM/mg of tissue (see **Supplementary Table 1**) and total parkin protein concentrations were estimated to be ~1.42 ng/mg brain tissue using r-parkin dilutions as standards; these had been run in parallel with brain lysates to demonstrate specificity and perform semiquantitative Western blots. To our knowledge, these estimates represent the first assessment of the concentration of wild-type parkin in adult mammalian brain.

As observed in r-parkin, we also found cysteine residues oxidized in parkin proteins after their affinity isolation from human control cortices and mouse brains, including of Zn^2+^-binding ones. For example, Cys253 (Cys252 in mice), which helps coordinate Zn^2+^ within parkin’s RING1 domain, was frequently identified by us as being oxidized (**Fig. 3i; Fig. 4e,g**). We predict that variable modifications of non-Zn^2+^-coordinating residues in human parkin, such as of Cys95, which is located in the - heretofore structurally understudied - linker region, or Cys59, as positioned in its ubiquitin-like (UbL) domain[38], could induce early, conformational changes in parkin’s tertiary structure (see **Figs. 6,7**). Such changes could profoundly affect parkin function mediated by other domains, as has been shown in several studies involving its E3 ligase activity as a readout following modifications in the UbL domain [27, 29-31, 34, 55, 74-76] (and reviewed in Yi e*t al.* [77]). Our results do not exclude the possibility that other non-thiol-based, posttranslational modifications alter parkin solubility, such as phosphorylation at Ser65 [78], or at Ser77 [38], which we found in MPTP-treated murine brain (not shown).

As mentioned above, *PRKN*-linked ARPD is pathologically restricted to catecholamine producing cells of the brainstem [3, 79-82]. Dopamine neurons of the *S. nigra* have unique biophysical properties that lead to high bioenergetic demands and the related rise in oxidative stress [83]. Further, unlike in other animals, dopamine is not completely catabolized in human brain, and neuromelanin is thought to be essential for the sequestration and long-term storage of its otherwise toxic metabolites [16]. We found parkin to be involved in mitigating two well-established, PD-linked stressors (*i.e.,* ROS; dopamine radicals), which is indirectly supported by our findings in human brain.

We show that parkin functions as a classical redox molecule that is able to lower H_2_O_2_ in a thiol-dependent manner. In the absence of wild-type parkin, H_2_O_2_ concentrations are elevated in human brain (**Fig. 5i**), in dopaminergic cells (**Fig. 5j,k**) and in brains from mice exposed to MPTP-toxin (**Fig. 5h**). There, acute MPTP exposure also correlated with the loss of parkin solubility and oxidation of its cysteines (**Fig. 4a**). Hence, *PRKN* expression contributes to the net reduction of H_2_O_2_ levels *in vivo.*

Because both MPTP toxin exposure and *Sod2* gene function affect mitochondrial integrity [84, 85], we reason that redox homeostasis in the cytosol, as coregulated by parkin oxidation, could also indirectly influence the health of mitochondria, in addition to E3 ligase-associated mitophagy (and MITAP). Such a cross-talk between cytosol and mitochondria likely includes glutathione metabolism-linked pathways, in which we and others found parkin cysteines to be involved in as well [21, 38, 86–89].

A role for *PRKN* expression in the neutralization and sequestration of dopamine metabolites may explain why dopamine synthesizing neurons are at great risk in humans with parkin deficiency. Previously, parkin has been shown to be uniquely sensitive to dopamine stress leading to aggregate formation [34, 55] (**Supplementary Fig. 6a,b**). In both cells and mice, *prkn* gene expression has been indirectly implicated in the metabolism of this neurotransmitter, in particular under *ex vivo* conditions of higher dopamine level-induced stress [21, 34, 68, 86, 90, 91] (see also **Supplementary Fig. 6c,d**).

Our results, and those by others, suggest that dopamine-mediated stress in neural cells is ameliorated when parkin undergoes modifications by dopamine metabolites. However, in contrast to current interpretations, which stipulate oxidation by quinones is equal to a loss of parkin activity, we posit that such oxidation is part of parkin’s physiological role within post-mitotic cells of the adult brain based on two principal findings. First, we demonstrate that wild-type parkin directly interacts with highly electrophilic dopamine metabolites at specific residues, foremost Cys95 (**Fig. 6**). This primate-specific cysteine is located within the linker region next to charged residues that impact its electrostatic properties and likely its redox reactivity [70, 92]. In support, we found that in addition to dopamine adduct conjugation, Cys95 is vulnerable to ROS attacks (**Figs. 3,4,6**), and in parallel studies, to be S-glutathionylated when exposed to rising concentrations of oxidized glutathione [38]. Strikingly, we found that Cys95 is not only required for parkin-dependent enhanced melanin formation, but also for participation in effective H_2_O_2_ reduction in M17 cells during dopamine toxicity (**Fig. 6e-g; Fig. 7e**).

Second, our finding that parkin augments melanin formation *in vitro*, together with our finding that the protein is closely associated with neuromelanin granules within LAMP-3^+^-lysosomes of human brain (**Fig.8; Supplementary Fig. 9**), suggest a role for parkin in *dopamine metabolism-linked neuroprotection* (**Supplementary Fig. 10**). We noted with interest that several autopsy reports have described lesser neuromelanin content in surviving neurons of the *S. nigra* in *PRKN*-linked ARPD [93–98] (**Fig. 8f**). Intriguingly, variants at the *LAMP3/CD63* locus, as well as of other dopamine metabolism-related genes, *e.g.*, *GCH-1,* have been recently identified as modifiers of susceptibility to late-onset, typical PD [99–101]. However, proof of any causality for parkin playing an essential role in neuromelanin formation awaits a suitable animal model.

To date, parkin is best known for its function as an E3 ligase, and the ubiquitin ligation-dependent involvement in mitophagy. Because ubiquitin-ligating activity occurs via cysteine-mediated trans-thiolation, controlling the redox state and functioning as an E3 ligase may not be mutually exclusive. For example, low concentrations of pro-oxidants, as well as sulfhydration, can activate parkin’s E3 activity *in vitro* [31, 32, 76]. A similar duality in functions, *i.e.,* regulating ubiquitylation and redox state in cells, has been previously described for the sensitive-to-apoptosis gene (SAG) product, also known as RBX2 / ROC2 / RNF7 [102, 103]. It contains a RING finger, and similar to parkin, was found to form HMW oligomers through oxidation of its cysteines [102, 103]. SAG protects cells from oxidative stress in a thiol-mediated manner in addition to functioning as an E3 ligase.

From this analogy, we postulate that parkin’s *cytoprotective E3 function* and role in mitophagy is possibly linked to its soluble form within the cytosol, which could be most important during early developmental stages, such as during cardiac development [13], in dividing striated muscle cells[104], and in relatively younger, neural cells including glia [88]. In support, Yi et al. recently described a strong correlation between parkin point mutants, their impact on structure and protein stability *vs.* ubiquitin ligase activity and the degree of mitophagy efficiency [77]. In addition, other parkin functions, such as those related to MITAP [9], inflammation signalling [10, 11] and redox-based neutralization of radicals could be more essential to the sustained health of older, postmitotic cells, *e.g., S. nigra* neurons.

The strength of our study is the focus on parkin metabolism in human midbrain and other tissues, which has never been undertaken before at these biochemical and structural levels since the gene’s discovery in 1998. In summary, we have shown that parkin fulfils criteria of a typical redox molecule: the sensing of oxidative (and reducing) stress via its thiols; and the direct, reciprocal redox regulation of its environment, thus conferring protective outcomes. If confirmed by future work, this redox chemistry-based expansion of parkin’s functions in the ageing human midbrain (**Supplementary Fig. 10**) may open the door to test its role in other neurodegenerative conditions, such as late-onset, non-*PRKN*-linked PD [105]. Most important, our findings emphasize the need for early identification of persons afflicted by *PRKN* gene mutations for the prioritization of appropriate interventions in the future, such as via gene therapy [106] and polyvalent, anti-oxidant therapy [107].

## Acknowledgments

We are very grateful for the commitment of patients and their families to participate in autopsy studies. We thank Dr. J. Palacino for creating stable M17 cell lines, Drs. A. Brice and E. Fon for sharing *prkn*-null mice, Dr. B. Madras for providing specimens of cynomolgus brain, Drs. R. Tam, L. Dong, Ms. K. Solti and Ms. H. Boston for technical support, Dr. D. Gibbings for antibodies, Dr. D. Gray for assistance with confocal imaging, Drs. M. Medina and R. R. Ratan for encouragement, Drs. S. Bennett, D. Pratt for discussions, and Drs. H. Lochmueller, M. Rousseaux and past members of the Schlossmacher lab for their suggestions.

## Author contributions

*Study design:* J.M.T., D.N.E.K., P.T., J.J.T., M.G.S.; *Writing and Figure preparation:* J.M.T., D.N.E.K., N.A.L., T.K.F., M.J., A.P.N., J.L., G.S.S., J.M.W., G.T., P.T., J.J.T., and M.G.S. prepared the initial draft of the manuscript and figures. All authors reviewed and / or edited the manuscript and approved of the submitted versions. *Experiments*: J.M.T., D.N.E.K., N.A.L., T.K.F., M.J., A.P.N., B.O., L.W., J.K., A.C.N., Q.J., R.S., J.L., M.Z., K.R.B., A.T., X.D., L.P., G.T. performed experiments; and C.R.S., A.B.W., E.T., A.H., A.P., J.A.C., provided data, tissue specimens and critical comments. *Analysis:* J.M.T., D.N.E.K., J.L., T.K.F., G.S.S., L.P., G.T., J.M.W., P.T., J.J.T., M.G.S. performed data analyses. *Study supervision*: P.T., J.J.T., M.G.S. *Overall responsibility:* M.G.S.

## Dedication

This work is dedicated to the memories of Mr. Bruce Hayter (1962-2019), a tireless advocate for persons with young-onset parkinsonism, and our co-author, Dr. Arne Holmgren (1940-2020), a pioneer in redox biology; both men died during the preparation of earlier versions of this manuscript. We are grateful to Dr. Oleh Hornykiewicz (1927-2020), the founding father of the dopamine era in neuroscience, for his tireless advocacy for biochemical investigations of the human brain and his mentorship.

## Additional information

### Data and materials availability

Original data associated with this study are available in the main text and supplementary figures and tables; additional data will be made available upon request.

### Supplementary Information

is available for this manuscript in the form of a videoclip.

### Correspondence and requests for materials

should be addressed to J.J.T or M.G.S.

---

**Supplementary Figure 1.**
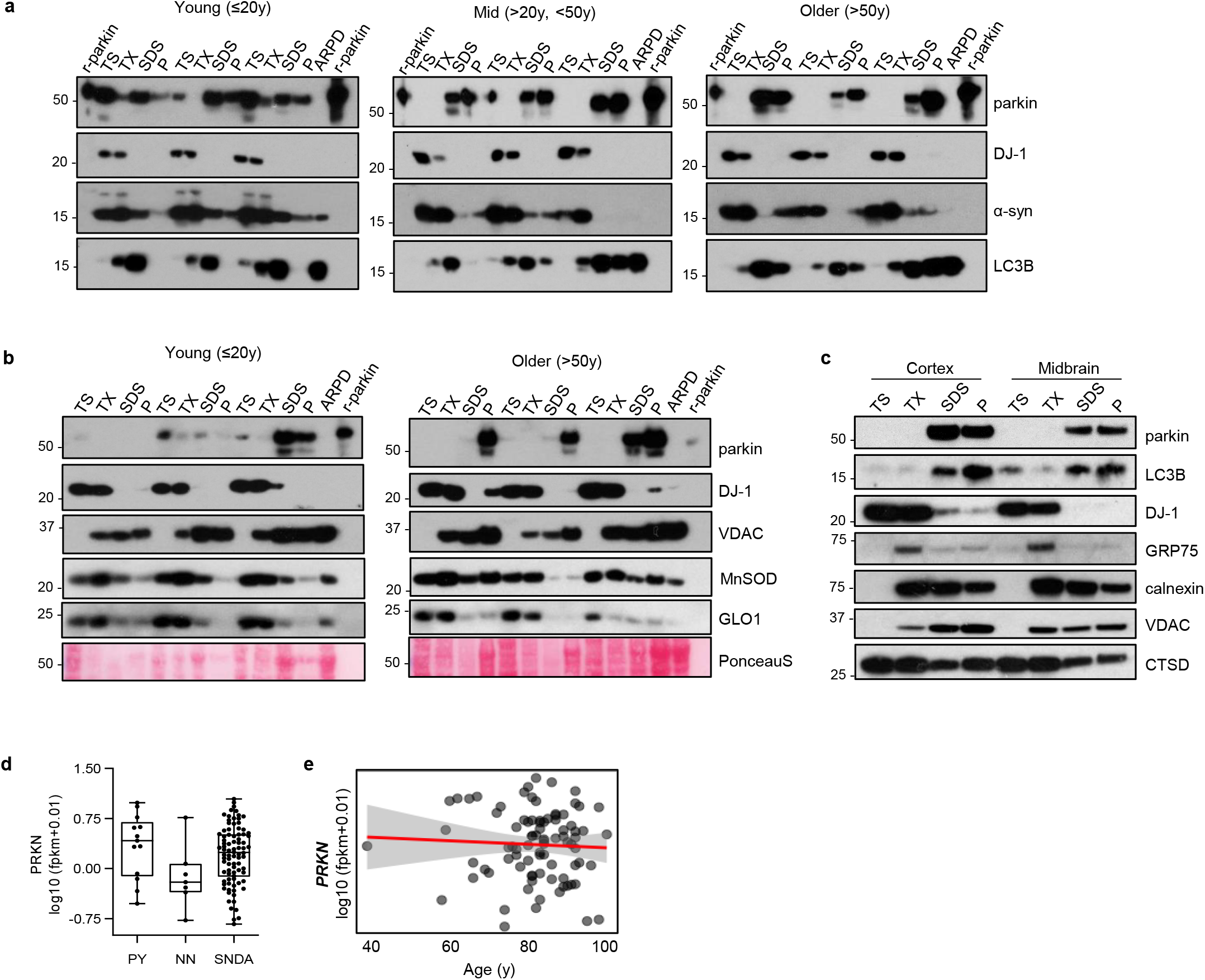
Parkin becomes progressively insoluble in aged human brain. **(a)** Western blots of parkin, DJ-1, α-synuclein and LC3B distribution in 9 representative human cortices (see **Extended Data Table 1**). Tissue fractionation and age ranges were as described in **Fig. 1**; SDS/PAGE experiments run under reducing conditions; SDS-extracted fractions of parkin-deficient PD brain (ARPD) lysates and r-parkin are included as controls. **(b)** Western blots of parkin, DJ-1, VDAC, MnSOD and glyoxalase-1 proteins, and Ponceau S staining in serially fractionated human cortices from younger (n=3) and older (n=3) individuals. Quantification of relative protein distribution is shown in **Fig. 1g**. **(c)** Western blot of indicated proteins from serially fractionated cortex and midbrain from a single donor as described in (a). **(d)** Quantification of log-transformed *PRKN* mRNA signals from individual pyramidal neurons (PY), leukocytes (non-neuronal cells; NN) and *S. nigra* dopamine neurons (SNDA) isolated from healthy controls (age range, 38 to 99 yrs). **(e)** Linear regression analysis of log-transformed *PRKN* transcripts as a function of age in human control *S. nigra* dopamine neurons where each dot represents values for a single neuron, as shown in (**d)**.

**Supplementary Figure 2.**
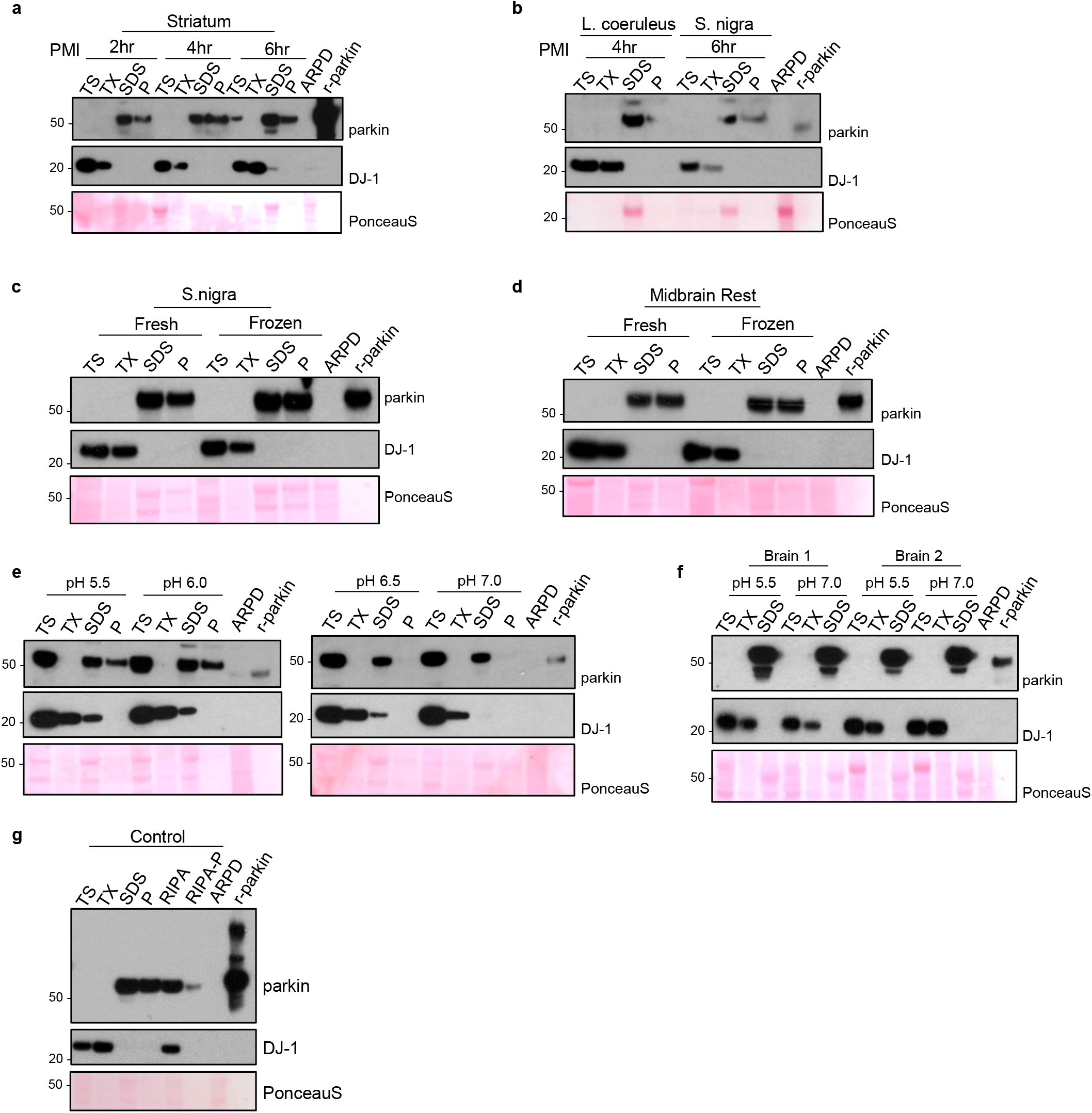
Parkin solubility is not altered by length of *post mortem* nterval, tissue freezing, or pH levels of the buffer. **(a-b)** Western blots of parkin and DJ-1 distribution as well as Ponceau S staining for fractions of human brain tissue from striatum **(a)**, *L. coeruleus* and *S. nigra* **(b)** with short *post mortem* interval (2-6 hrs, as indicated). **(c-d)** Western blots, as described in (a), from dissected *S. nigra* **(c)** and posterior midbrain structures comprising nucleus of cranial nerve-III and the periaqueductal grey (**d;** rest). Tissues were collected *post mortem* and parkin distribution visualized in aliquots of the same specimens processed in parallel after being kept at 4°C or processed via one-time freezing and thawing prior to serial fractionation. **(e-f)** Western blots of parkin and DJ-1 distribution as well as Ponceau S staining in fractions of human cortex (**e**, single brain; **f**, two different brains) serially extracted in parallel using standard buffers with varying pH, as indicated. **(g)** Western blots of parkin and DJ-1 distribution in a human cortex sample following serial fractionation with TS-TX-, SDS- and Pellet buffers compared to processing by standard RIPA buffer, where the pellet after RIPA extraction is denoted as RIPA-P.

**Supplementary Figure 3.**
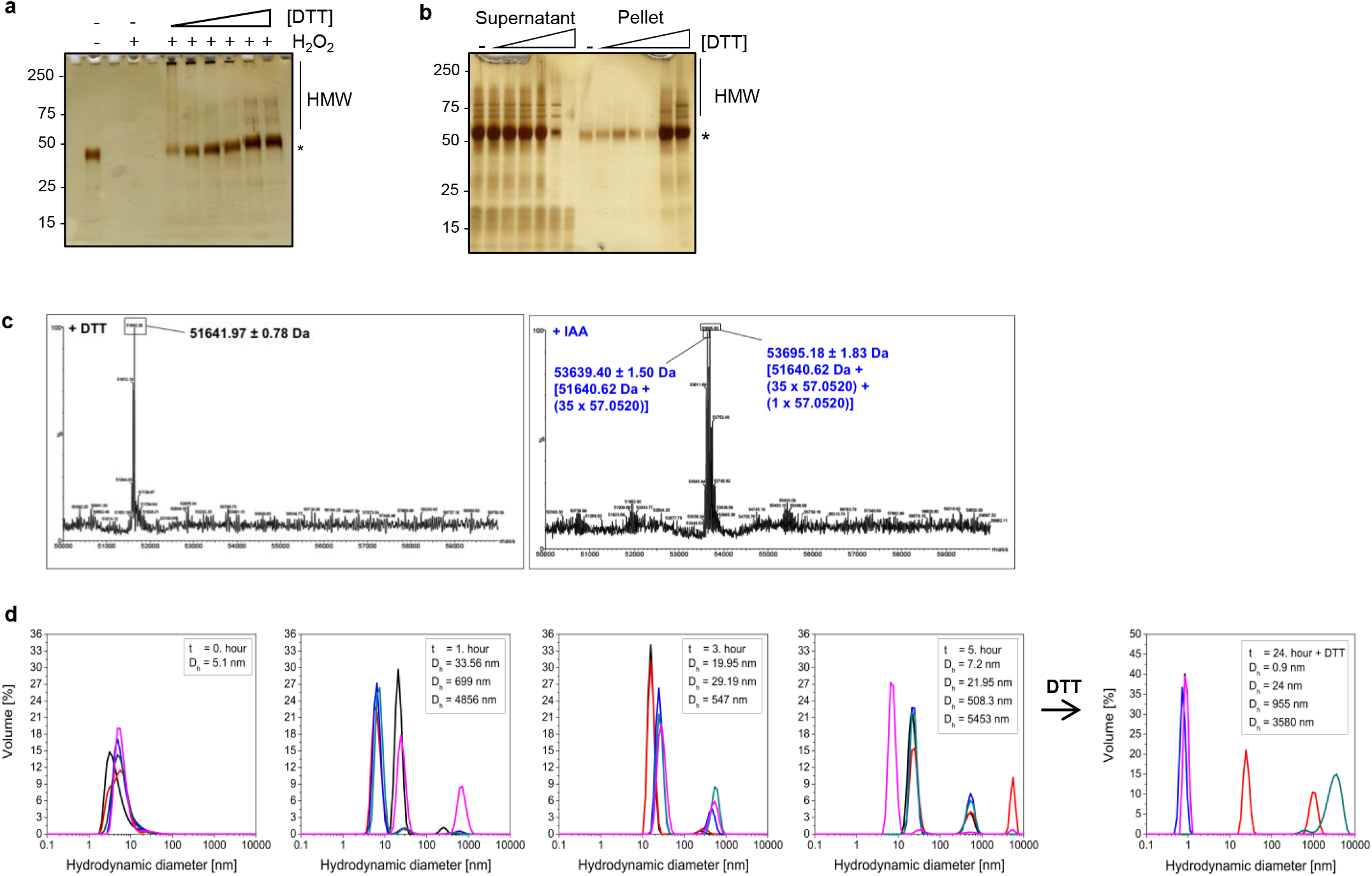
Oxidation of parkin thiols promotes insolubility. **(a)** Silver stained gel of wild-type, human r-parkin exposed to H_2_O_2_ (10 mM), followed by treatment with increasing concentrations of DTT (0-100 mM) prior to centrifugation and loading of the supernatant onto SDS-PAGE. **(b)** Detection of r-parkin in soluble (supernatant) and insoluble phases (pellet; recovered by 10% SDS-containing buffer) following exposure to increasing concentrations of DTT (0-1M). **(c)** Spectra from LC-MS/MS analyses of recombinant (r-), human, wild-type parkin holoprotein (without any trypsin digestion) without pre-labelling (panel on the left) and after tagging of 35 *vs.* 36 thiol-carrying residues by iodoacetamide (IAA; right panel), corresponding to the three main peaks (one in left panel; two in right panel), as indicated. The 51,641.97 Da peak closely matches its calculated mass of 51,640.62 Da; 53,639.40 Da corresponds to the conjugation of 35 IAA adducts; 53,695.18 Da corresponds to 36 IAA adducts, indicating that all 35 cysteine residues and either the N-terminal amino group or a single methionine residue was IAA-modified. **(d)** Dynamic light scattering analysis showing progressive size changes, as measured in hydrodynamic diameters (nm), monitored during 0, 1, 3 and 5 hrs at room temperature. The structural state for wild-type, human r-parkin under non-reducing, native conditions showed increased aggregate formation over time, which was partially reversed by DTT.

**Supplementary Figure 4.**
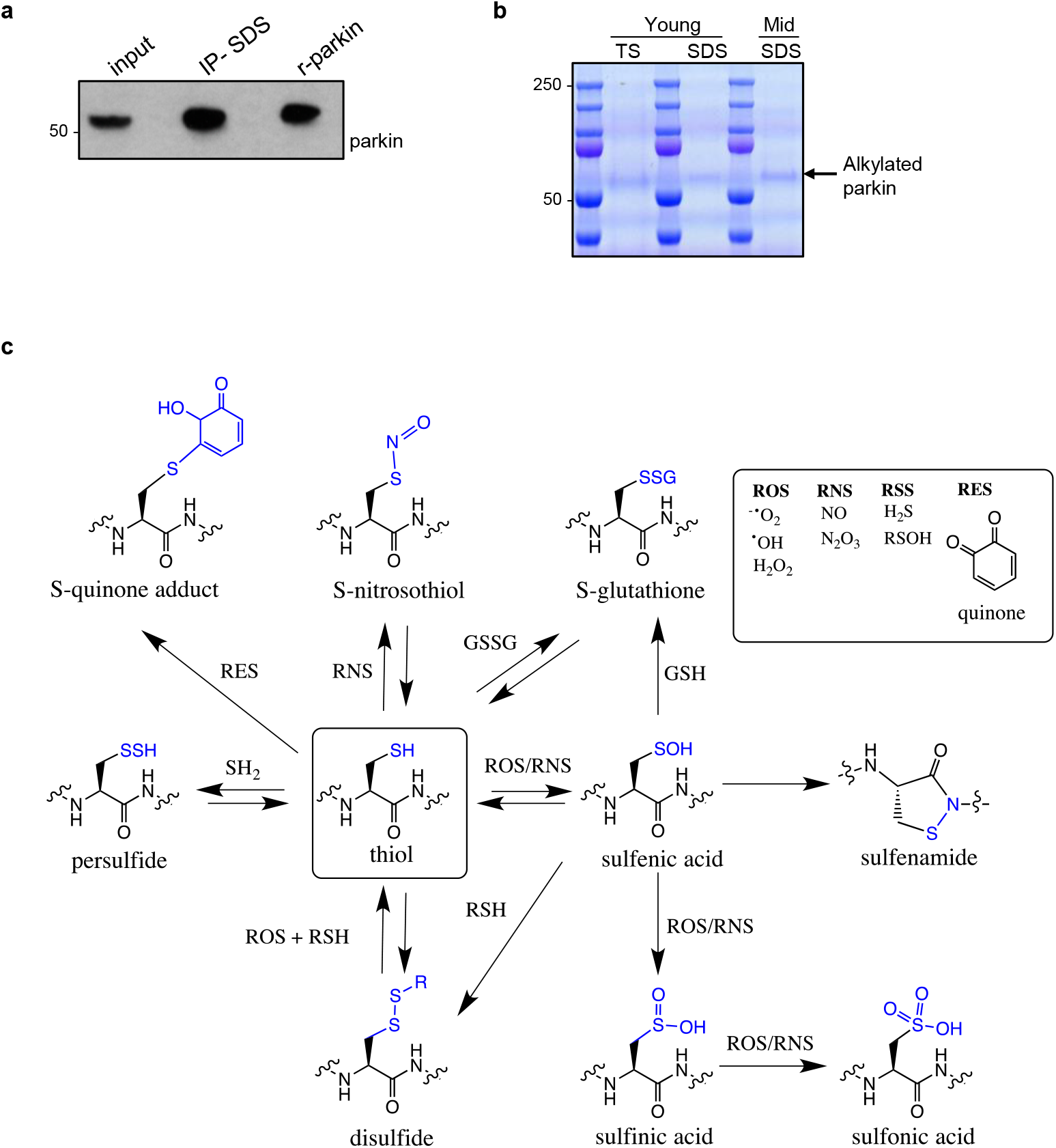
Immunoprecipitation of brain parkin and summary of redox-related thiol chemistry. **(a-b)** Representative Western blot **(a)** and Coomassie blue-stained **(b)** visualization of parkin immunoprecipitated from human frontal lobe cortex, as described [Shimura and Schlossmacher, Methods Enzymol 2005] by monoclonal anti-parkin A15165-B and visualized by polyclonal anti-parkin 2132, in preparation for LC-MS/MS (see also **Fig. 4**). Brain tissue was homogenized in the presence of IAA to prevent the oxidation of reduced thiols during processing, thereby generating alkylated-parkin monomers at the 51-54 kDa position. **(c)** Schema of select, reversible and irreversible cysteine modifications that can occur on thiols (-SH) due to attacks by reactive oxygen species (ROS), reactive nitrogen species (RNS), reactive sulfur species (RSS) and reactive electrophilic species (RES), which include dopamine quinones. Graphic summary was modified from Alcock *et al.,* 2018.

**Supplementary Figure 5.**
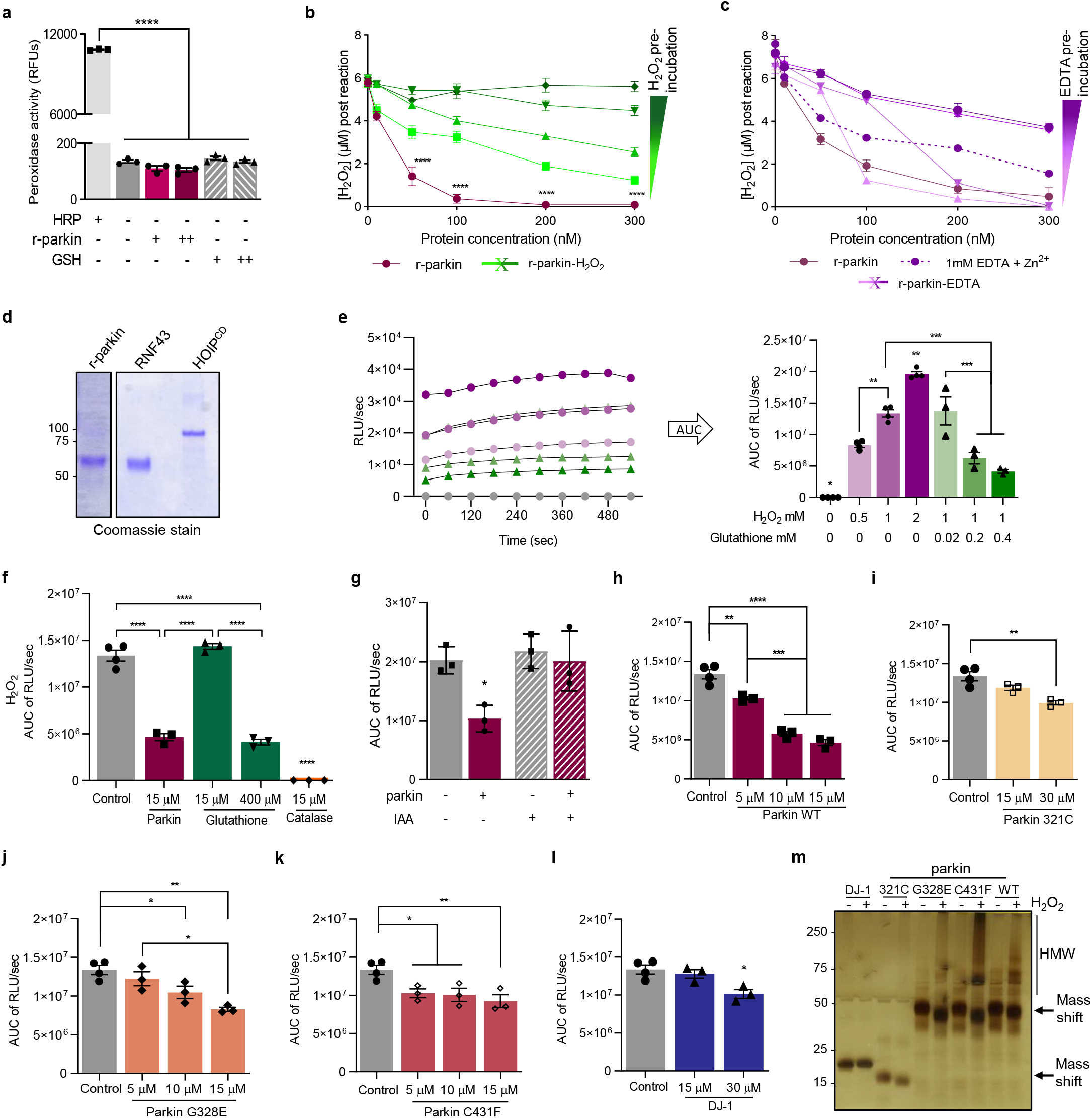
Parkin directly reduces hydrogen peroxide in a concentration- and thiol integrity-dependent but non-enzymatic manner. **(a)** Peroxidase enzymatic activity for r-parkin and glutathione (GSH; +, 0.5μM; ++, 1μM), as tested *in vitro* in comparison to horseradish peroxidase (HRP, 1mU/mL). Mean peroxidase activity ± SEM. ****P<0.0001 by 1-way ANOVA. **(b-c)** Quantification of H_2_O_2_ concentrations by AmplexRed following incubation of increasing levels of r-parkin **(b)** pre-oxidized with increasing concentrations of H_2_O_2_, or **(c)** treated with increasing concentrations of EDTA. A two-way ANOVA was used for statistical analysis (****P<0.0001). **(d)** Commassie Blue-stained visualization of r-parkin, RNF43 and HOIP^cd^ proteins, used in the AmplexRed assay shown in **Fig. 5c**. **(e)** Kinetic readings from *in vitro* colorimetric H_2_O_2_ assays (left panel) comparing increasing concentrations of input H_2_O_2_ (green lines) and the effect of increasing concentrations of glutathione (purple, pink lines). Curves were converted to the area under the curve (AUC) where AUC integrates the total value of H_2_O_2_ signals generated over the 10 mins time course of the assay (right panel). **(f-l)** AUC graphs for results from *in vitro* H_2_O_2_ assays for various concentrations of recombinant proteins, as indicated. Statistical analysis was performed as in **Fig. 5e**. **(m)** Visualization of recombinant PD proteins post H_2_O_2_ exposure by silver staining where SDS/PAGE gel was run under non-reducing conditions.

**Supplementary Figure 6.**
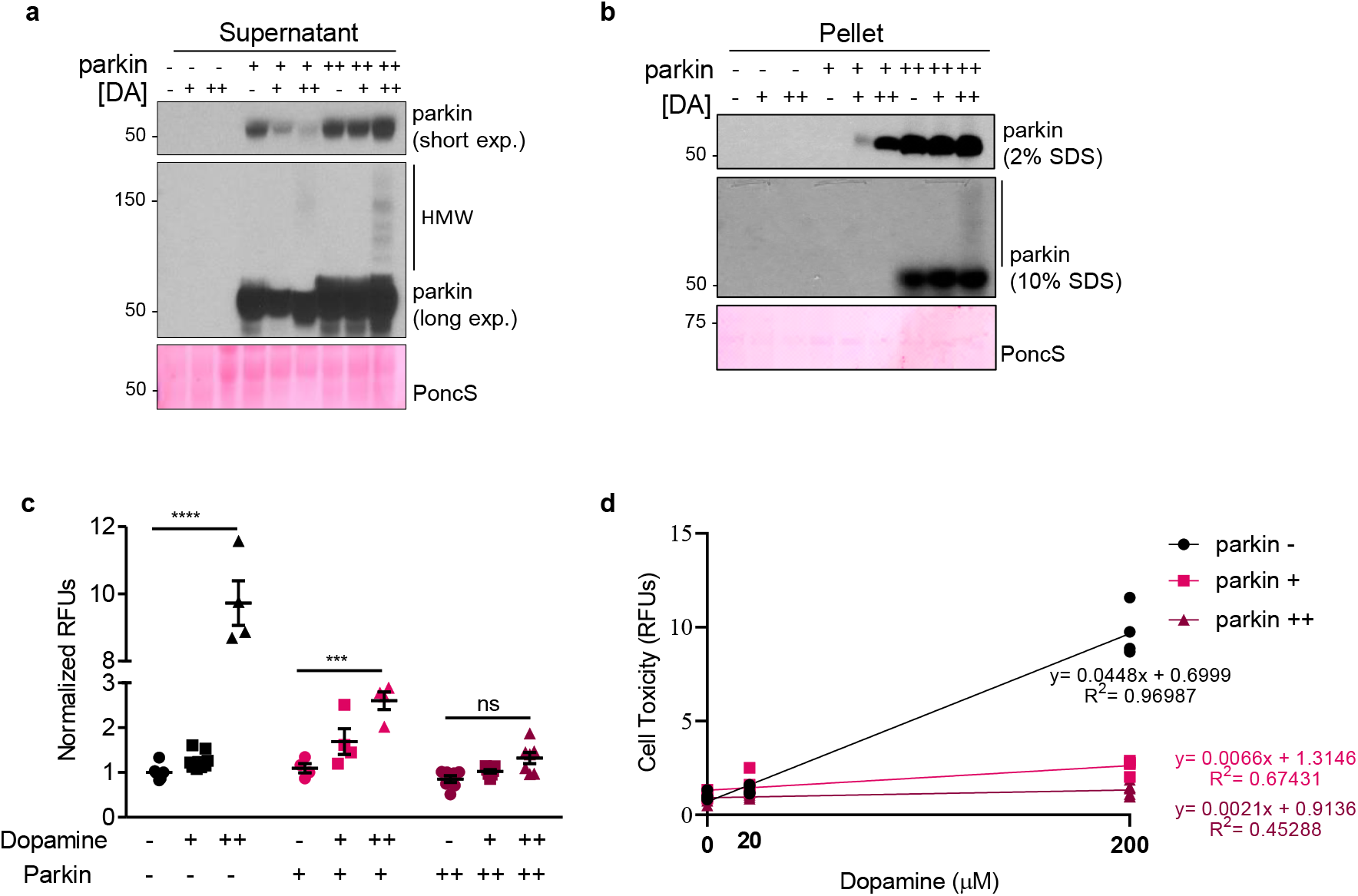
Parkin protects neural cells from dopamine toxicity in a protein concentration-dependent manner. **(a-b)** Western blots of parkin in the soluble supernatant (**a**) and insoluble, serial pellet (**b**) fractions of lysates from dopamine-treated human M17 neuroblastoma cells, which stably express vector-control plasmid (parkin -) or human *PRKN* cDNA at mid-(+) or high (++) levels. Cells were exposed to 20 mM (+) and 200 mM (++) dopamine for 20 hrs, as indicated. SDS/PAGE gels were run under reducing conditions. **(c)** Cell viability assay of cells highlighted in (a, b). Representative data are shown for the mean of duplicates ± SEM from n=4-8 independent experiments; *P<0.05 by 1-way ANOVA. **(d)** Correlation studies of experiments, as conducted in (a, b), to monitor parkin expression levels *vs*. cell survival.

**Supplementary Figure 7.**
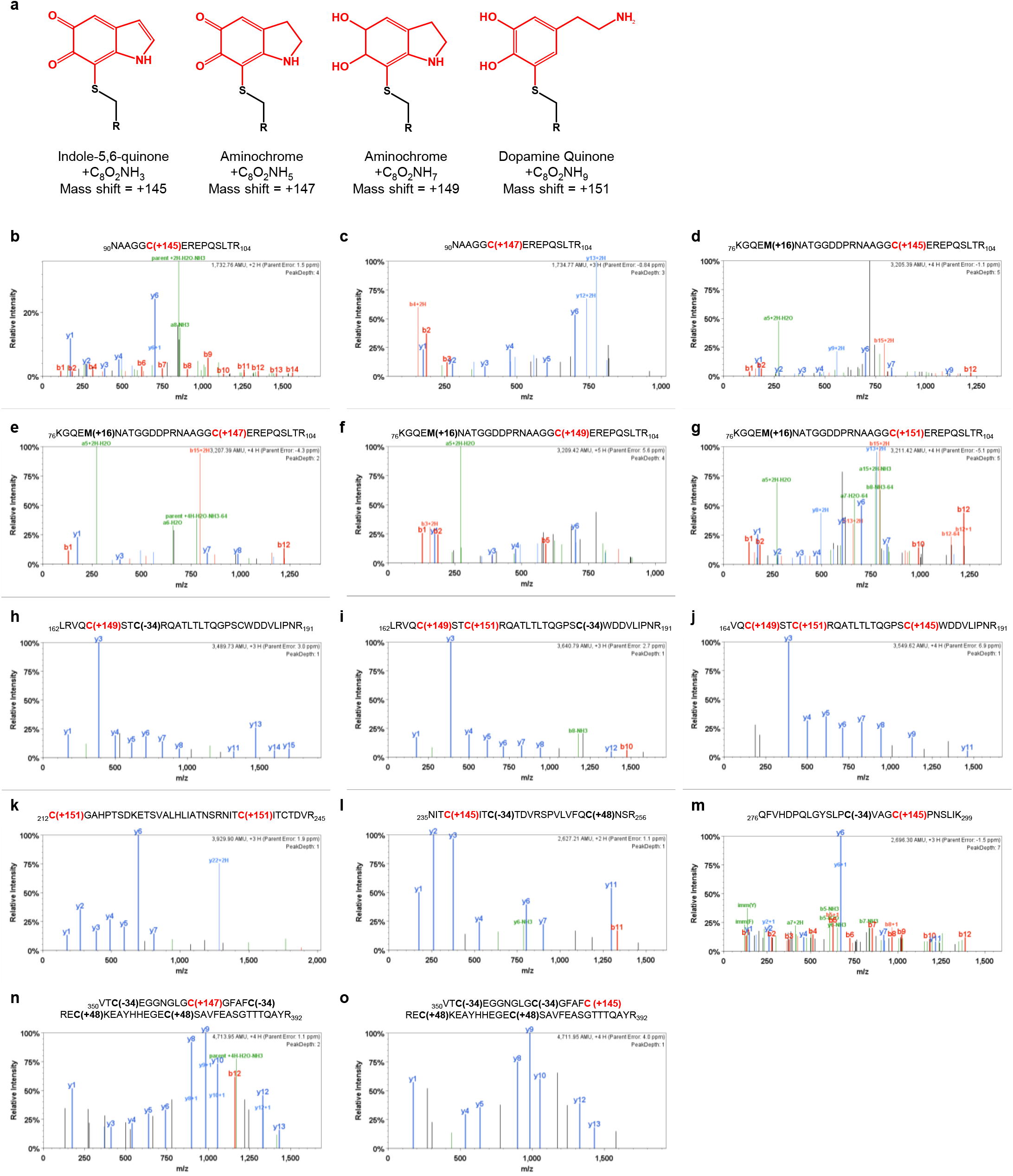
Human parkin conjugates dopamine metabolites at cysteine 95 and other cysteine residues. **(a)** Chemical structures of 4 dopamine metabolites (red) conjugated to a thiol group (black) that were screened for in LC-MS/MS experiments, with their corresponding mass shift added. **(b-o)** LC-MS/MS-generated spectra following trypsin digestion of aminochrome-treated, human r-parkin protein highlighting representative adduct conjugation events, which were identified by mass shift gains as shown in (**a**), at the following residues: Cys95 (**b-g**), Cys166, Cys169 and Cys 182 (**h-j**), to Cys212 (**k**), Cys238 (**l**), Cys293 (**m**), Cys360 (**n**), and Cys365 (**o**). See also **Fig. 6e,g**.

**Supplementary Figure 8.**
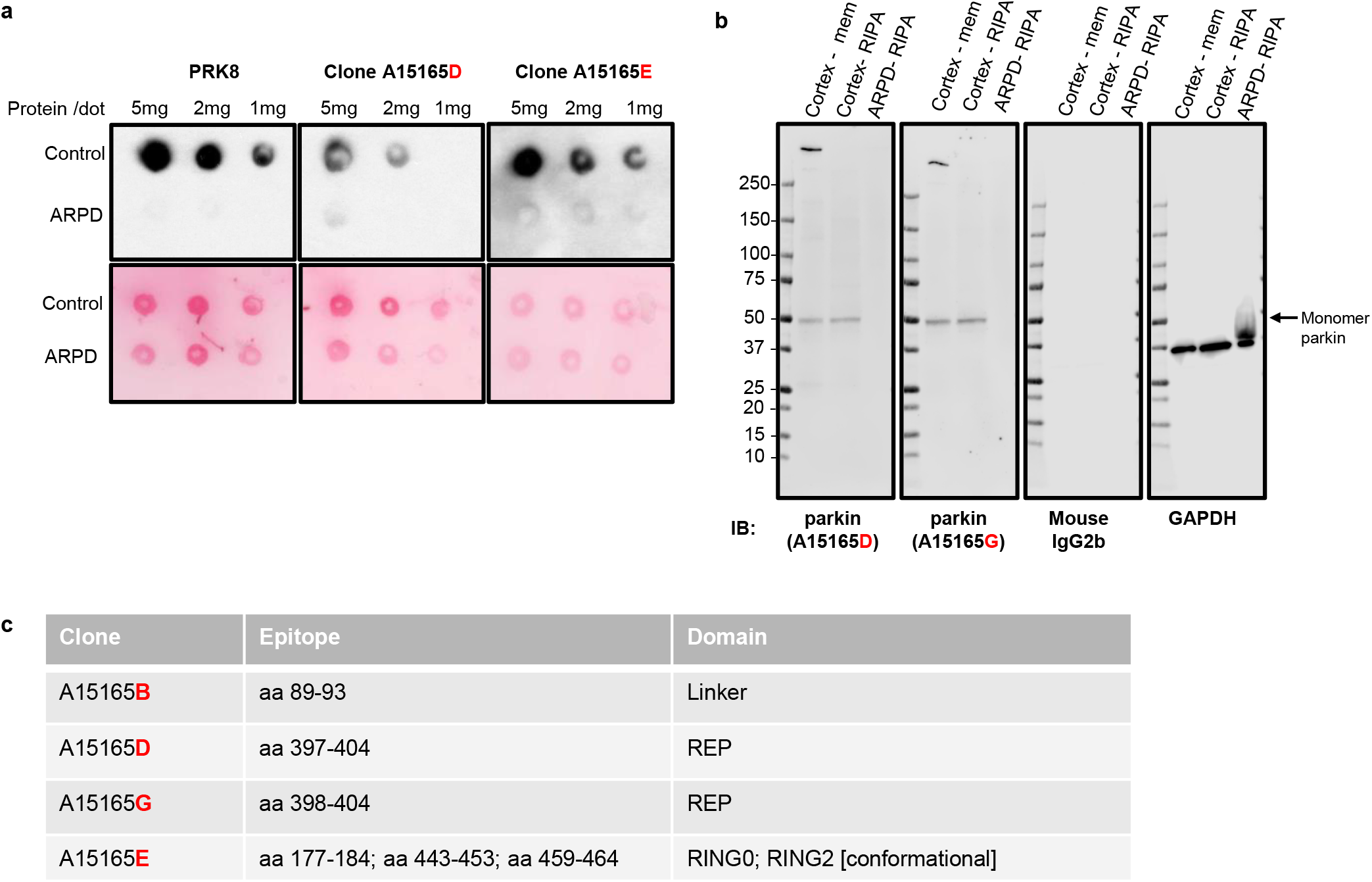
Characterization of four, new monoclonal antibodies raised in mice against human parkin. **(a-b)** Characterization of four murine, monoclonal antibodies (of IgG_2_ isotype; clone-B, -E, -D, and -G) by **(a)** non-denaturing dot blots against human brain lysates (SDS fractions from control and *PRKN-*linked ARPD cases); and **(b)** by denaturing SDS/PAGE under reducing conditions and Western blotting of extracts from cortical specimens of a control brain and a parkin-deficient ARPD case. Screening by these three methods as well as by cell-based microscopy using indirect immunofluorescence (not shown) revealed specific staining for four anti-parkin clones (-B, -E, -D and -G), which was conformation-dependent for clone-E. **(c)** List of epitopes within the sequence of human parkin, as recognized by clones -B, -E, -D, and -G and identified by screening with overlapping 7-12 amino acid-long peptides covering full-length, human parkin. Note, the clone E epitope is conformational, comprised of the three regions as indicated.

**Supplementary Figure 9.**
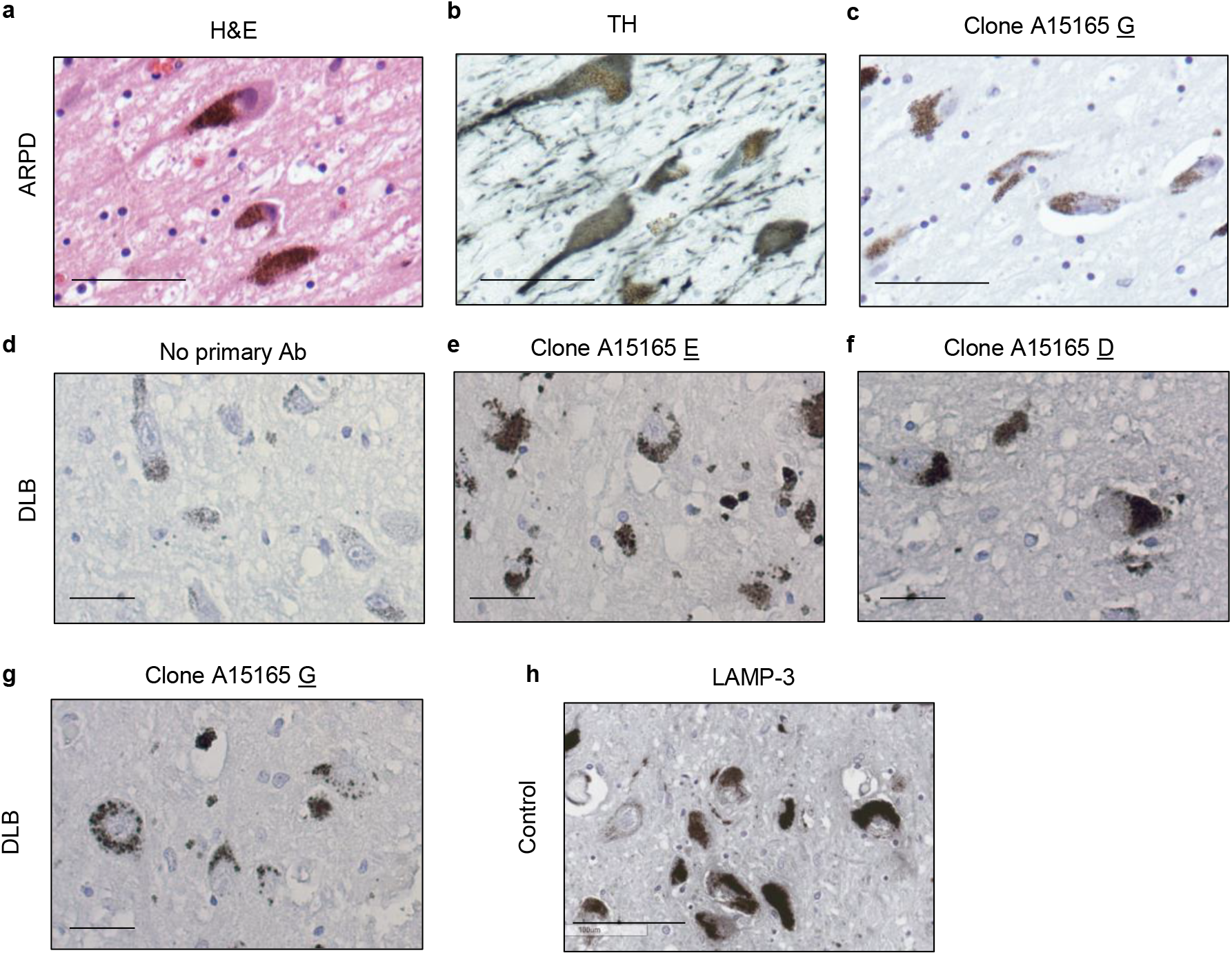
Parkin is specifically detected in human brain sections by routine microscopy. **(a-c)** H&E **(a)**, anti-tyrosine hydroxylase (TH) **(b)** and anti-parkin (clone A15165-G) (**c**) staining of dopamine neurons in the *S. nigra* of midbrain sections from a parkin-deficient ARPD case. (**d-g**) Immunohistochemical detection of parkin in the *S. nigra* of an individual with dementia with Lewy bodies. Both intra- and extracellular anti-parkin-reactive neuromelanin granules are visible. **(b)** No primary antibody control and staining with anti-parkin monoclonal antibodies **(e)** A15165-E, **(f)**-D and **(g)**-G are shown. **(h)** Immunohistochemical detection of LAMP-3 protein in dopamine neurons of the *S. nigra* from an adult control brain. Scale bars represent 100 mm.

**Supplementary Figure 10.**
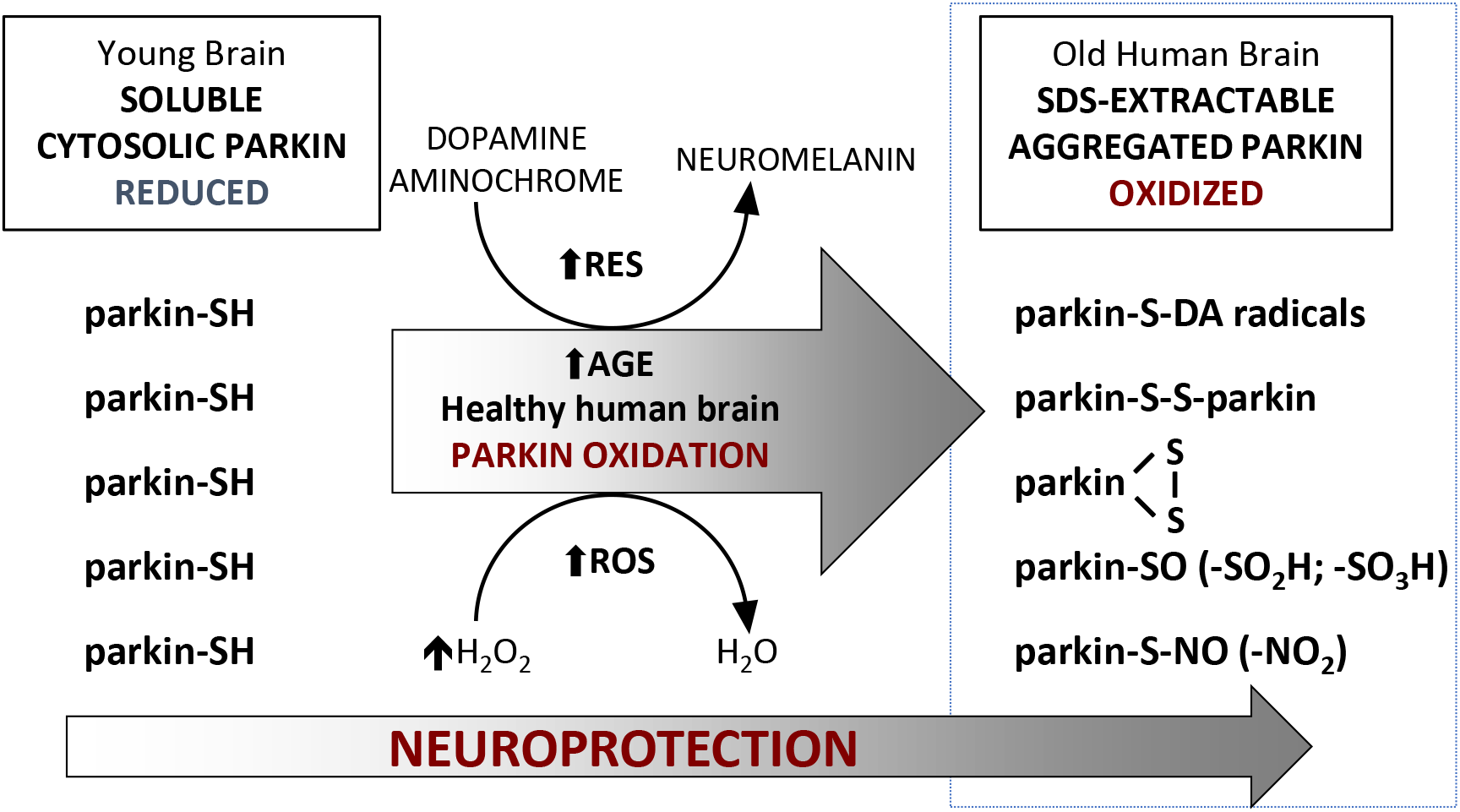
Graphic summary of a working model for parkin’s redox functions in adult, human dopamine neurons. In human brain, parkin thiol (-SH) oxidation neutralizes cellular reactive oxygen species (ROS; H_2_O_2_) and potentially toxic dopamine (DA) radicals (*e.g.,* DA quinones; RES) during normal ageing. The oxidation of wild-type parkin promotes insolubility and aggregation. In human brain, both reversible and irreversible oxidation events occur, which promote parkin’s transition into a mostly insoluble, aggregate-associated state by the beginning of the 5^th^ decade. In adult dopamine neurons of the *S. nigra*, its oxidation and aggregation lead to the accumulation of a pool of parkin within LAMP-3-positive lysosomes. This multimodal oxidation of parkin confers neuroprotection. In *PRKN*-linked ARPD, the absence of parkin’s redox effects contributes to a rise in ROS (and RNS) levels, reduced sequestration of dopamine radicals (RES), and possibly, less neuromelanin formation.

**Supplementary Table 1.** List of human tissue specimens examined in this study. Characteristics listed include brain regions of frontal cortex (F ctx), midbrain, thoracic spinal cord (harvested with skeletal muscle); age (in years); sex (F, female; M, male); PMI, *post mortem* interval recorded in hours (hrs); n.d., not determined with accuracy (*i.e.,* inconsistent PMI information); brain diagnosis, where * indicates that the tissue examined was not affected by a detectable disease process; parkin solubility lists 1 for it being present in Tris-saline (TS) buffer or 0 (absent in TS-buffer); and the figure(s) that specimens were analyzed in this study; E, extended data.

| Sample ID | Brain region | Age | Sex | PMI (hrs) | Diagnosis | Parkin Solubility | H<sub>2</sub>O<sub>2</sub>/ Tissue ratio | Appears in figures |
| --- | --- | --- | --- | --- | --- | --- | --- | --- |
| 1 | FC | 5 | F | 33 | Healthy control | 1 |  | 1b, 1e, 1f, 1g, 1h, 1k, E1a, E1b |
| 2 | FC | 5 | F | 20 | Healthy control | 1 | 2.551 | 1b, 1e, 1f, 1g, 1h, 1k, 2<sup>a</sup>, 2b, 2c, E1a, E1b |
| 3 | FC | 8 | M | 5 | Healthy control | 0 | 1.780 | 1b, 1e, 1k, 2a, 2b, 2c |
| 4 | FC | 13 | M | 13 | Healthy control | 1 | 2.217 | 1b, 1e, 1f, 1k, 2a, 2b, 2c |
| 5 | FC | 15 | F | 9 | Healthy control | 1 | 1.349 | 1b, 1e, 1f, 1k, 2a, 2b, 2c |
| 6 | FC | 16 | F | 20 | Healthy control | 1 |  | 1b, 1e, 1f, 1g, 1h, E1b |
| 7 | FC | 16 | F | 14 | Healthy control | 1 | 1.016 | 1b, 1e, 1f, 1k, 2a, 2b, 2c, E1a |
| 8 | FC | 17 | M | 23 | Healthy control | 0 | 0.701 | 1b, 1e, 1k, 2a, 2b, 2c |
| 9 | FC | 17 | M | 22 | Healthy control | 1 |  | 1b, 1e, 1k |
| 10 | FC | 20 | F | 19 | Healthy control | 1 |  | 1b, 1e, 1f, 1k |
| 11 | FC | 20 | M | 8 | Healthy control | 1 |  | 1b, 1e, 1f, 1k |
| 12 | FC | 20 | M | 6 | Healthy control | 0 |  | 1b, 1e, 1f, 1k |
| 13 | FC | 20 | M | 5 | Healthy control | 1 |  | 1b, 1e, 1f, 1k |
| 14 | FC | 21 | M | 30 | Healthy control | 1 |  | 1b, 1e, 1f, 1k |
| 15 | FC | 28 | M | 33 | Epilepsy * | 1 |  | 1b, 1e, 1f, 1k |
| 16 | FC | 29 | F | 18 | Epilepsy * | 0 | 3.061 | 1b, 1e, 1f, 1k, 2a, 2b, 2c, E1a |
| 17 | FC | 30 | M | 20 | Healthy control | 0 | 6.089 | 1b, 1e, 1f, 1k, 2a, 2b, 2c |
| 18 | FC | 36 | M | 20 | Healthy control | 0 | 5.665 | 1b, 1e, 1f, 1k, 2a, 2b, 2c |
| 19 | FC | 37 | F | 13 | Healthy control | 0 |  | 1b, 1e,1f, 1k |
| 20 | FC | 38 | M | 17 | Healthy control | 0 |  | 1b, 1e, 1f, 1k |
| 21 | FC | 39 | M | 23 | Healthy control | 1 |  | 1b, 1e, 1f, 1k |
| 22 | FC | 39 | M | 14 | Healthy control | 1 |  | 1b, 1e, 1f, 1k, E1a |
| 23 | FC | 42 | M | 18 | Healthy control | 1 |  | 1b, 1e, 1f, 1k |
| 24 | FC | 43 | F | 22 | Healthy control | 0 | 4.601 | 1b, 1e, 1f, 1k, 2a, 2b, 2c, E1a |
| 25 | FC | 44 | F | 21 | Spina bifida * | 0 |  | 1b, 1e, 1k, E1c |
| 26 | FC | 49 | F | 16 | Healthy control | 0 | 5.622 | 1b, 1e, 1f, 1k, 2a, 2b, 2c |
| 27 | FC | 49 | F | 14 | Healthy control | 0 |  | 1b, 1e, 1k |
| 28 | FC | 54 | M | 16 | Alzheimer disease | 0 |  | 1b, 1e, 1f, 1k |
| 29 | FC | 54 | F | 23 | Huntington disease | 1 |  | 1b, 1e, 1k |
| 30 | FC | 55 | F | 16 | Healthy control | 0 | 5.829 | 1b, 1e, 1f, 1k, 2a, 2b, 2c |
| 31 | FC | 56 | M | 23 | Healthy control | 0 |  | 1b, 1e, 1f, 1g, 1h, 1k, E1b |
| 32 | FC | 57 | M | n.d. | Healthy control | 0 |  | 1e |
| 33 | FC | 62 | M | 15 | Brain hemorrhage * | 0 | 6.525 | 1b, 1e, 1f, 1k, 2a, 2b, 2c, E1a |
| 34 | FC | 65 | M | 5 | Lewy body dementia | 0 | 6.473 | 1e, 1j, 1k, 1l, 1m, 2a, 2b, 2c |
| 35 | FC | 65 | M | 14 | Sporadic Parkinson's | 0 | 9.112 | 1e, 1k, 2a, 2b, 2c |
| 36 | FC | 65 | F | 42 | Healthy control | 0 | 5.768 | 1a, 1b, 1e, 1k, 2a, 2b, 2c, E2e |
| 37 | FC | 66 | M | n.d. | Healthy control | 0 | 6.674 | 1b, 1e, 1f, 2a, 2b, 2c, E1a |
| 38 | FC | 68 | M | 17 | Healthy control | 0 | 2.514 | 1b, 1e, 1f, 1g, 1h, 1k, 2a, 2b, 2c, E1b |
| 39 | FC | 70 | F | n.d. | Healthy control | 0 | 6.897 | 1b, 1e, 1f, 2a, 2b, 2c, E1a |
| 40 | FC | 70 | M | n.d. | Healthy control | 0 | 5.459 | 1b, 1e, 1f, 1g, 1h, 2a, 2b, 2c, E1b |
| 41 | FC | 72 | M | n.d. | Lewy body dementia | 1 | 6.274 | 1e, 2a, 2b, 2c |
| 42 | FC | 75 | M | 48 | Healthy control | 1 | 2.878 | 1b, 1e, 1k, 2a, 2b, 2c |
| 43 | FC | 75 | F | 13 | Ischaemic stroke * | 0 |  | 1b, 1e, 1f, 1k |
| 44 | FC | 75 | M | 17 | Lewy body dementia | 0 |  | 1e, 1j, 1k |
| 45 | FC | 76 | M | 74 | Pick's disease | 0 |  | 1e, 1k |
| 46 | FC | 85 | F | 15 | Alzheimer disease | 0 |  | 1b, 1e, 1f, 1j, 1k |
| ARPD1 | FC | 66 | M | 17 | PRKN-linked Parkinson's | n/a |  |  |
| ARPD2 | FC | 57 | M | 31 | PRKN-linked Parkinson's | n/a |  |  |
| ARPD3 | FC | 70 | F | 42 | PRKN-linked Parkinson's | n/a |  |  |
| ARPD4 | FC | 70 | M | 13 | PRKN-linked Parkinson's | n/a |  |  |
| 47 | MB | 26 | M | 2 | Multiple sclerosis * | 0 |  | 1b, E2b |
| 65 | MB | 34 | M | n.d. | Encephalitis* | 0 |  | 1b |
| 25 | MB | 44 | F | 21 | Healthy control | 0 |  | 1b |
| 48 | MB | 44 | F | 5 | Multiple sclerosis * | 1 |  | 1c, E2b |
| 49 | MB | 45 | M | 13 | Ischaemic stroke * | 0 |  | 1b |
| 50 | MB | 47 | F | 20 | Brain hemorrhage * | 0 |  | 1b |
| 51 | MB | 56 | M | 44 | Multiple system atrophy | 0 |  | 1b |
| 52 | MB | 60 | M | 16 | Progressive supranuclear palsy | 0 |  | 1b |
| 53 | MB | 61 | M | 20 | Healthy control | 1 |  | 1b |
| 54 | MB | 61 | M | 3.5 | Multiple sclerosis * | 0 |  | 1b |
| 36 | MB | 65 | F | 42 | Healthy control | 0 |  | 1b, E2a |
| 55 | MB | 65 | M | 6 | Multiple sclerosis * | 0 |  | 1b |
| 64 | MB | 71 | M | n.d. | Lewy body dementia | 0 |  | 1b |
| 41 | MB | 72 | M | n.d. | Lewy body dementia | 1 |  | 1b |
| 56 | MB | 74 | F | n.d. | Amyotrophic lateral sclerosis | 0 |  | 1b |
| 42 | MB | 75 | M | 48 | Healthy control | 0 |  | 1b |
| 57 | MB | 75 | M | 70 | Sporadic Parkinson's | 0 |  | 1b |
| 45 | MB | 76 | M | 74 | Pick's disease | 0 |  | 1b |
| 58 | MB | 79 | M | n.d. | Progressive supranuclear palsy | 0 |  | 1b |
| 59 | MB | 82 | M | 48 | Lewy body dementia | 1 |  | 1b |
| 60 | SC/Muscle | 68 | F | 2.5 | Ischaemic stroke * | 1 |  | 1c, 1d |
| 61 | SC/Muscle | 64 | F | 2 | Subarachnoid hemorrhage* | 1 |  | 1d |
| 62 | SC/Muscle | 74 | M | 2 | Cerebellar hemorrhage * | 1 |  | 1d |
| 63 | SC/Muscle | 50 | M | 4 | Subarachnoid hemorrhage * | 1 |  | 1d |

**Supplementary Table 2.** Parkin’s cysteine residues are redox active. Aliquots of human recombinant (r-) parkin that were oxidized by variable concentrations of H_2_O_2_ *vs.* control preparations were differentially labelled with iodoacetamide (IAA) and/or N-ethylmaleimide (NEM; as in Figure 4A) to identify reduced cysteines (IAA) or reversibly-oxidized residues (NEM). Proteins were subjected to LC-MS/MS and analyzed using Mascot Scaffold PTM to identify IAA (•) or NEM (**+**) adducts indicating when these were detectable on individual residues. Cysteines that were not detected as modified in individual runs are also listed (n/d). Note that cysteines within all four RING domains of parkin as well as in the linker and UbL domains can be variably modified.

| Treatment |  | Control | Control | H<sub>2</sub>O<sub>2</sub> | H<sub>2</sub>O<sub>2</sub> | H<sub>2</sub>O<sub>2</sub> | H<sub>2</sub>O<sub>2</sub> | H<sub>2</sub>O<sub>2</sub> |
| --- | --- | --- | --- | --- | --- | --- | --- | --- |
| Run |  | IAA+NEM | IAA | 20µM | 1mM | 4.5mM | 4.5mM | 4.5mM |
| Region | Cysteine Residue |  |  |  |  |  |  |  |
| UBL | 59 |  | • | n/d | • | • + | • | • |
| Linker | 95 | • | • | • | • | • + | • + | • + |
| RING0 | 150 | • | • | • | • | • + | • + | • |
|  | 154 | • | • | n/d | • | • + | • + | • |
|  | 166 | n/d | n/d | • | • + | n/d | • | • |
|  | 169 | n/d | • | • | • + | n/d | • | • |
|  | 182 | • | • | • | n/d | • + | • + | • |
|  | 196 | • | • | • | • | • + | • + | • + |
|  | 201 | • | • | • | • | • + | • + | • + |
|  | 212 | • | • | • | n/d | • + | • + | • + |
| RING1 | 238 | • | • | • + | • | • + | • | • + |
|  | 241 | • | • | • + | • | • + | • | • |
|  | 253 | • | • | • + | • | • + | • | • |
|  | 260 | • | • | n/d | n/d | • + | • | • |
|  | 263 | • | • | n/d | n/d | • + | • | • |
|  | 268 | • | • | n/d | n/d | • + | • | • |
|  | 289 | • | • | n/d | n/d | • + | • + | • + |
|  | 293 | • | • | n/d | • | • | • + | • + |
|  | 323 | • | • | n/d | n/d | • + | • + | • + |
| IBR | 332 | • | • | n/d | n/d | • + | • | • |
|  | 337 | • | • | n/d | • | • + | • | • + |
|  | 352 | • | • | • | • | • | • + | • + |
|  | 360 | • | • | • | • | • + | • | • |
|  | 365 | • | • | • | n/d | • + | • | • |
|  | 368 | • | • | n/d | n/d | • + | • + | • |
|  | 377 | • | • | • | n/d | • + | • + | • + |
| RING2 | 418 | n/d | n/d | n/d | • | n/d | n/d | n/d |
|  | 421 | n/d | n/d | n/d | • + | n/d | • + | • + |
|  | 431 | n/d | • | n/d | • | n/d | • | • |
|  | 436 | n/d | n/d | n/d | • + | n/d | • | • |
|  | 441 | n/d | n/d | n/d | • + | n/d | • | • |
|  | 446 | • | • | n/d | n/d | • + | • + | • + |
|  | 449 | • | • | n/d | n/d | • + | • + | • |
|  | 451 | • | • | n/d | n/d | • + | • | • |
|  | 457 | • | • | • | • | • + | • | • + |
|  | % Parkin protein coverage | 83 | 89 | 57 | 60 | 86 | 98 | 97 |
|  | # peptides identified | 40 | 35 | 22 | 33 | 38 | 51 | 47 |
|  | # Cysteines identified | 27 | 31 | 16 | 21 | 28 | 34 | 34 |
|  | # IAA-cysteines | 27 | 30 | 16 | 21 | 28 | 34 | 34 |
|  | IAA-cys/identified-cys | 27/27 | 30/31 | 16/16 | 21/21 | 28/28 | 34/34 | 34/34 |
|  | % | 100 | 97 | 100 | 100 | 100 | 100 | 100 |
|  | # NEM-cysteines | 0 | n/a | 3 | 5 | 26 | 16 | 14 |
|  | NEM-cys/identified-cys | 0/27 | n/a | 3/16 | 5/21 | 26/28 | 16/34 | 14/34 |
|  | % | 0 | n/a | 19 | 24 | 93 | 47 | 41 |

• IAA + NEM

## References

1. T. Kitada, S. Asakawa, N. Hattori, H. Matsumine, Y. Yamamura, S. Minoshima, et al., Mutations in the parkin gene cause autosomal recessive juvenile parkinsonism, Nature 392(6676) (1998) 605–8.

2. M. Kasten, C. Hartmann, J. Hampf, S. Schaake, A. Westenberger, E.J. Vollstedt, et al., Genotype-Phenotype Relations for the Parkinson’s Disease Genes Parkin, PINK1, DJ1: MDSGene Systematic Review, Mov Disord 33(5) (2018) 730–741.

3. K.M. Doherty, L. Silveira-Moriyama, L. Parkkinen, D.G. Healy, M. Farrell, N.E. Mencacci, et al., Parkin disease: a clinicopathologic entity?, JAMA Neurol 70(5) (2013) 571–9.

4. A.K. Berger, G.P. Cortese, K.D. Amodeo, A. Weihofen, A. Letai, M.J. LaVoie, Parkin selectively alters the intrinsic threshold for mitochondrial cytochrome c release, Hum Mol Genet 18(22) (2009) 4317–28.

5. N. Matsuda, S. Sato, K. Shiba, K. Okatsu, K. Saisho, C.A. Gautier, et al., PINK1 stabilized by mitochondrial depolarization recruits Parkin to damaged mitochondria and activates latent Parkin for mitophagy, J Cell Biol 189(2) (2010) 211–21.

6. D.P. Narendra, S.M. Jin, A. Tanaka, D.F. Suen, C.A. Gautier, J. Shen, et al., PINK1 is selectively stabilized on impaired mitochondria to activate Parkin, PLoS Biol 8(1) (2010) e1000298.

7. G.L. McLelland, V. Soubannier, C.X. Chen, H.M. McBride, E.A. Fon, Parkin and PINK1 function in a vesicular trafficking pathway regulating mitochondrial quality control, EMBO J 33(4) (2014) 282–95.

8. A.K. Muller-Rischart, A. Pilsl, P. Beaudette, M. Patra, K. Hadian, M. Funke, et al., The E3 ligase parkin maintains mitochondrial integrity by increasing linear ubiquitination of NEMO, Mol Cell 49(5) (2013) 908–21.

9. D. Matheoud, A. Sugiura, A. Bellemare-Pelletier, A. Laplante, C. Rondeau, M. Chemali, et al., Parkinson’s Disease-Related Proteins PINK1 and Parkin Repress Mitochondrial Antigen Presentation, Cell 166(2) (2016) 314–327.

10. F. Mouton-Liger, T. Rosazza, J. Sepulveda-Diaz, A. Ieang, S.M. Hassoun, E. Claire, et al., Parkin deficiency modulates NLRP3 inflammasome activation by attenuating an A20-dependent negative feedback loop, Glia 66(8) (2018) 1736–1751.

11. D.A. Sliter, J. Martinez, L. Hao, X. Chen, N. Sun, T.D. Fischer, et al., Parkin and PINK1 mitigate STING-induced inflammation, Nature (2018).

12. S.K. Barodia, R.B. Creed, M.S. Goldberg, Parkin and PINK1 functions in oxidative stress and neurodegeneration, Brain Res Bull 133 (2017) 51–59.

13. G. Gong, M. Song, G. Csordas, D.P. Kelly, S.J. Matkovich, G.W. Dorn2nd,, Parkin-mediated mitophagy directs perinatal cardiac metabolic maturation in mice, Science 350(6265) (2015) aad2459.

14. A.M. Pickrell, C.H. Huang, S.R. Kennedy, A. Ordureau, D.P. Sideris, J.G. Hoekstra, et al., Endogenous Parkin Preserves Dopaminergic Substantia Nigral Neurons following Mitochondrial DNA Mutagenic Stress, Neuron 87(2) (2015) 371–81.

15. J.H. Shin, H.S. Ko, H. Kang, Y. Lee, Y.I. Lee, O. Pletinkova, et al., PARIS (ZNF746) repression of PGC-1alpha contributes to neurodegeneration in Parkinson’s disease, Cell 144(5) (2011) 689–702.

16. F.A. Zucca, E. Basso, F.A. Cupaioli, E. Ferrari, D. Sulzer, L. Casella, et al., Neuromelanin of the human substantia nigra: an update, Neurotox Res 25(1) (2014) 13–23.

17. A.J. Whitworth, D.A. Theodore, J.C. Greene, H. Beneš, P.D. Wes, L.J. Pallanck, Increased glutathione S-transferase activity rescues dopaminergic neuron loss in a Drosophila model of Parkinson’s disease, Proceedings of the National Academy of Sciences 102(22) (2005) 8024–8029.

18. P. Ge, V.L. Dawson, T.M. Dawson, PINK1 and Parkin mitochondrial quality control: a source of regional vulnerability in Parkinson’s disease, Mol Neurodegener 15(1) (2020) 20.

19. J.J. Palacino, D. Sagi, M.S. Goldberg, S. Krauss, C. Motz, M. Wacker, et al., Mitochondrial dysfunction and oxidative damage in parkin-deficient mice, J Biol Chem 279(18) (2004) 18614–22.

20. M. Periquet, O. Corti, S. Jacquier, A. Brice, Proteomic analysis of parkin knockout mice: alterations in energy metabolism, protein handling and synaptic function, J Neurochem 95(5) (2005) 1259–76.

21. J.M. Itier, P. Ibanez, M.A. Mena, N. Abbas, C. Cohen-Salmon, G.A. Bohme, et al., Parkin gene inactivation alters behaviour and dopamine neurotransmission in the mouse, Hum Mol Genet 12(18) (2003) 2277–91.

22. S.J. Lee, D.G. Kim, K.Y. Lee, J.S. Koo, B.J. Lee, Regulatory mechanisms of thiol-based redox sensors: lessons learned from structural studies on prokaryotic redox sensors, Arch Pharm Res 41(6) (2018) 583–593.

23. L.J. Alcock, M.V. Perkins, J.M. Chalker, Chemical methods for mapping cysteine oxidation, Chem Soc Rev 47(1) (2018) 231–268.

24. J. Segura-Aguilar, I. Paris, P. Munoz, E. Ferrari, L. Zecca, F.A. Zucca, Protective and toxic roles of dopamine in Parkinson’s disease, J Neurochem 129(6) (2014) 898–915.

25. V.A. Hristova, S.A. Beasley, R.J. Rylett, G.S. Shaw, Identification of a novel Zn2+-binding domain in the autosomal recessive juvenile Parkinson-related E3 ligase parkin, J Biol Chem 284(22) (2009) 14978–86.

26. M.J. LaVoie, G.P. Cortese, B.L. Ostaszewski, M.G. Schlossmacher, The effects of oxidative stress on parkin and other E3 ligases, J Neurochem 103(6) (2007) 2354–68.

27. F. Meng, D. Yao, Y. Shi, J. Kabakoff, W. Wu, J. Reicher, et al., Oxidation of the cysteine-rich regions of parkin perturbs its E3 ligase activity and contributes to protein aggregation, Mol Neurodegener 6 (2011) 34.

28. K.F. Winklhofer, I.H. Henn, P.C. Kay-Jackson, U. Heller, J. Tatzelt, Inactivation of parkin by oxidative stress and C-terminal truncations: a protective role of molecular chaperones, J Biol Chem 278(47) (2003) 47199–208.

29. K.K. Chung, B. Thomas, X. Li, O. Pletnikova, J.C. Troncoso, L. Marsh, et al., S-nitrosylation of parkin regulates ubiquitination and compromises parkin’s protective function, Science 304(5675) (2004) 1328–31.

30. K.K. Chung, V.L. Dawson, T.M. Dawson, S-nitrosylation in Parkinson’s disease and related neurodegenerative disorders, Methods Enzymol 396 (2005) 139–50.

31. D. Yao, Z. Gu, T. Nakamura, Z.Q. Shi, Y. Ma, B. Gaston, et al., Nitrosative stress linked to sporadic Parkinson’s disease: S-nitrosylation of parkin regulates its E3 ubiquitin ligase activity, Proc Natl Acad Sci U S A 101(29) (2004) 10810–4.

32. M.S. Vandiver, B.D. Paul, R. Xu, S. Karuppagounder, F. Rao, A.M. Snowman, et al., Sulfhydration mediates neuroprotective actions of parkin, Nat Commun 4 (2013) 1626.

33. J. Chakraborty, V. Basso, E. Ziviani, Post translational modification of Parkin, Biol Direct 12(1) (2017) 6.

34. M.J. LaVoie, B.L. Ostaszewski, A. Weihofen, M.G. Schlossmacher, D.J. Selkoe, Dopamine covalently modifies and functionally inactivates parkin, Nat Med 11(11) (2005) 1214–21.

35. C.H. Polman, S.C. Reingold, B. Banwell, M. Clanet, J.A. Cohen, M. Filippi, et al., Diagnostic criteria for multiple sclerosis: 2010 revisions to the McDonald criteria, Ann Neurol 69(2) (2011) 292–302.

36. T. Dhaeze, L. Tremblay, C. Lachance, E. Peelen, S. Zandee, C. Grasmuck, et al., CD70 defines a subset of proinflammatory and CNS-pathogenic TH1/TH17 lymphocytes and is overexpressed in multiple sclerosis, Cell Mol Immunol 16(7) (2019) 652–665.

37. T. Kuhlmann, S. Ludwin, A. Prat, J. Antel, W. Bruck, H. Lassmann, An updated histological classification system for multiple sclerosis lesions, Acta Neuropathol 133(1) (2017) 13–24.

38. D.N. El Kodsi, J.M. Tokarew, R. Sengupta, N.A. Lengacher, A.C. Ng, H. Boston, et al., Parkinson Disease-Linked Parkin Mediates Redox Reactions That Lower Oxidative Stress In Mammalian Brain, bioRxiv (2020) 2020.04.26.062380.

39. X. Dong, Z. Liao, D. Gritsch, Y. Hadzhiev, Y. Bai, J.J. Locascio, et al., Enhancers active in dopamine neurons are a primary link between genetic variation and neuropsychiatric disease, Nat Neurosci 21(10) (2018) 1482–1492.

40. D.E. Spratt, R.J. Martinez-Torres, Y.J. Noh, P. Mercier, N. Manczyk, K.R. Barber, et al., A molecular explanation for the recessive nature of parkin-linked Parkinson’s disease, Nat Commun 4 (2013) 1983.

41. A. Kumar, J.D. Aguirre, T.E. Condos, R.J. Martinez-Torres, V.K. Chaugule, R. Toth, et al., Disruption of the autoinhibited state primes the E3 ligase parkin for activation and catalysis, EMBO J 34(20) (2015) 2506–21.

42. J.D. Aguirre, K.M. Dunkerley, P. Mercier, G.S. Shaw, Structure of phosphorylated UBL domain and insights into PINK1-orchestrated parkin activation, Proc Natl Acad Sci U S A 114(2) (2017) 298–303.

43. R. Kiss, M. Zhu, B. Jojart, A. Czajlik, K. Solti, B. Forizs, et al., Structural features of human DJ-1 in distinct Cys106 oxidative states and their relevance to its loss of function in disease, Biochim Biophys Acta Gen Subj 1861(11 Pt A) (2017) 2619–2629.

44. K. Solti, Kuan, W. L., Fórizs, B., Kustos, G., Judith, M., Varga, Z., Herberth, B., Moravcsik, E., Kiss, R., Kárpáti, M., Mikes, A., Zhao, Y., Imre, T., Rochet, J.C., Aigbirhio, F., Williams-Gray, C. H., Barker, R. A., Tóth, G., DJ-1 can form beta-sheet structured aggregates that co-localize with pathological amyloid deposits, Neurobiology of Disease (2019).

45. C.H. Muller, T.K. Lee, M.A. Montano, Improved chemiluminescence assay for measuring antioxidant capacity of seminal plasma, Methods Mol Biol 927 (2013) 363–76.

46. A. Shevchenko, H. Tomas, J. Havlis, J.V. Olsen, M. Mann, In-gel digestion for mass spectrometric characterization of proteins and proteomes, Nat Protoc 1(6) (2006) 2856–60.

47. J. Cox, M. Mann, MaxQuant enables high peptide identification rates, individualized p.p.b.-range mass accuracies and proteome-wide protein quantification, Nat Biotechnol 26(12) (2008) 1367–72.

48. S.F. Ali, S.N. David, G.D. Newport, J.L. Cadet, W. Slikker, Jr., MPTP-induced oxidative stress and neurotoxicity are age-dependent: evidence from measures of reactive oxygen species and striatal dopamine levels, Synapse 18(1) (1994) 27–34.

49. M.G. Schlossmacher, M.P. Frosch, W.P. Gai, M. Medina, N. Sharma, L. Forno, et al., Parkin localizes to the Lewy bodies of Parkinson disease and dementia with Lewy bodies, Am J Pathol 160(5) (2002) 1655–67.

50. M.G. Schlossmacher, H. Shimura, Parkinson’s disease: assays for the ubiquitin ligase activity of neural Parkin, Methods Mol Biol 301 (2005) 351–69.

51. B. Shutinoski, M. Hakimi, I.E. Harmsen, M. Lunn, J. Rocha, N. Lengacher, et al., Lrrk2 alleles modulate inflammation during microbial infection of mice in a sex-dependent manner, Sci Transl Med 11(511) (2019).

52. A.C. Pawlyk, B.I. Giasson, D.M. Sampathu, F.A. Perez, K.L. Lim, V.L. Dawson, et al., Novel monoclonal antibodies demonstrate biochemical variation of brain parkin with age, J Biol Chem 278(48) (2003) 48120–8.

53. I. Liguori, G. Russo, F. Curcio, G. Bulli, L. Aran, D. Della-Morte, et al., Oxidative stress, aging, and diseases, Clin Interv Aging 13 (2018) 757–772.

54. A. Krezel, W. Maret, The biological inorganic chemistry of zinc ions, Arch Biochem Biophys 611 (2016) 3–19.

55. E.S. Wong, J.M. Tan, C. Wang, Z. Zhang, S.P. Tay, N. Zaiden, et al., Relative sensitivity of parkin and other cysteine-containing enzymes to stress-induced solubility alterations, J Biol Chem 282(16) (2007) 12310–8.

56. E. Ferrari, M. Engelen, E. Monzani, M. Sturini, S. Girotto, L. Bubacco, et al., Synthesis and structural characterization of soluble neuromelanin analogs provides important clues to its biosynthesis, J Biol Inorg Chem 18(1) (2013) 81–93.

57. E. Ferrari, A. Capucciati, I. Prada, F.A. Zucca, G. D’Arrigo, D. Pontiroli, et al., Synthesis, Structure Characterization, and Evaluation in Microglia Cultures of Neuromelanin Analogues Suitable for Modeling Parkinson’s Disease, ACS Chem Neurosci 8(3) (2017) 501–512.

58. H. Shimura, N. Hattori, S. Kubo, M. Yoshikawa, T. Kitada, H. Matsumine, et al., Immunohistochemical and subcellular localization of Parkin protein: absence of protein in autosomal recessive juvenile parkinsonism patients, Ann Neurol 45(5) (1999) 668–72.

59. P.P. Pramstaller, M.G. Schlossmacher, T.S. Jacques, F. Scaravilli, C. Eskelson, I. Pepivani, et al., Lewy body Parkinson’s disease in a large pedigree with 77 Parkin mutation carriers, Ann Neurol 58(3) (2005) 411–22.

60. F.A. Zucca, R. Vanna, F.A. Cupaioli, C. Bellei, A. De Palma, D. Di Silvestre, et al., Neuromelanin organelles are specialized autolysosomes that accumulate undegraded proteins and lipids in aging human brain and are likely involved in Parkinson’s disease, NPJ Parkinsons Dis 4 (2018) 17.

61. C. Hampe, H. Ardila-Osorio, M. Fournier, A. Brice, O. Corti, Biochemical analysis of Parkinson’s disease-causing variants of Parkin, an E3 ubiquitin-protein ligase with monoubiquitylation capacity, Hum Mol Genet 15(13) (2006) 2059–75.

62. W.J. Gu, O. Corti, F. Araujo, C. Hampe, S. Jacquier, C.B. Lucking, et al., The C289G and C418R missense mutations cause rapid sequestration of human Parkin into insoluble aggregates, Neurobiol Dis 14(3) (2003) 357–64.

63. C. Wang, H.S. Ko, B. Thomas, F. Tsang, K.C. Chew, S.P. Tay, et al., Stress-induced alterations in parkin solubility promote parkin aggregation and compromise parkin’s protective function, Hum Mol Genet 14(24) (2005) 3885–97.

64. C. Wang, J.M. Tan, M.W. Ho, N. Zaiden, S.H. Wong, C.L. Chew, et al., Alterations in the solubility and intracellular localization of parkin by several familial Parkinson’s disease-linked point mutations, J Neurochem 93(2) (2005) 422–31.

65. S.R. Sriram, X. Li, H.S. Ko, K.K. Chung, E. Wong, K.L. Lim, et al., Familial-associated mutations differentially disrupt the solubility, localization, binding and ubiquitination properties of parkin, Hum Mol Genet 14(17) (2005) 2571–86.

66. M.R. Cookson, P.J. Lockhart, C. McLendon, C. O’Farrell, M. Schlossmacher, M.J. Farrer, RING finger 1 mutations in Parkin produce altered localization of the protein, Hum Mol Genet 12(22) (2003) 2957–65.

67. D.H. Hyun, M. Lee, N. Hattori, S. Kubo, Y. Mizuno, B. Halliwell, et al., Effect of wild-type or mutant Parkin on oxidative damage, nitric oxide, antioxidant defenses, and the proteasome, J Biol Chem 277(32) (2002) 28572–7.

68. H. Jiang, Y. Ren, E.Y. Yuen, P. Zhong, M. Ghaedi, Z. Hu, et al., Parkin controls dopamine utilization in human midbrain dopaminergic neurons derived from induced pluripotent stem cells, Nat Commun 3 (2012) 668.

69. J. Okarmus, H. Bogetofte, S.I. Schmidt, M. Ryding, S. Garcia-Lopez, B.J. Ryan, et al., Lysosomal perturbations in human dopaminergic neurons derived from induced pluripotent stem cells with PARK2 mutation, Sci Rep 10(1) (2020) 10278.

70. H. Xiao, M.P. Jedrychowski, D.K. Schweppe, E.L. Huttlin, Q. Yu, D.E. Heppner, et al., A Quantitative Tissue-Specific Landscape of Protein Redox Regulation during Aging, Cell 180(5) (2020) 968–983 e24.

71. G.M. Jordan, S. Yoshioka, T. Terao, The aggregation of bovine serum albumin in solution and in the solid state, J Pharm Pharmacol 46(3) (1994) 182–5.

72. G. Paris, S. Kraszewski, C. Ramseyer, M. Enescu, About the structural role of disulfide bridges in serum albumins: evidence from protein simulated unfolding, Biopolymers 97(11) (2012) 889–98.

73. W. Maret, Zinc coordination environments in proteins as redox sensors and signal transducers, Antioxid Redox Signal 8(9–10) (2006) 1419–41.

74. K. Ozawa, A.T. Komatsubara, Y. Nishimura, T. Sawada, H. Kawafune, H. Tsumoto, et al., S-nitrosylation regulates mitochondrial quality control via activation of parkin, Sci Rep 3 (2013) 2202.

75. T. Wauer, D. Komander, Structure of the human Parkin ligase domain in an autoinhibited state, EMBO J 32(15) (2013) 2099–112.

76. N. Panicker, V.L. Dawson, T.M. Dawson, Activation mechanisms of the E3 ubiquitin ligase parkin, Biochem J 474(18) (2017) 3075–3086.

77. W. Yi, E.J. MacDougall, M.Y. Tang, A.I. Krahn, Z. Gan-Or, J.F. Trempe, et al., The landscape of Parkin variants reveals pathogenic mechanisms and therapeutic targets in Parkinson’s disease, Hum Mol Genet 28(17) (2019) 2811–2825.

78. F. Koyano, K. Okatsu, H. Kosako, Y. Tamura, E. Go, M. Kimura, et al., Ubiquitin is phosphorylated by PINK1 to activate parkin, Nature 510(7503) (2014) 162–6.

79. C.B. Lucking, A. Durr, V. Bonifati, J. Vaughan, G. De Michele, T. Gasser, et al., Association between early-onset Parkinson’s disease and mutations in the parkin gene, N Engl J Med 342(21) (2000) 1560–7.

80. N.L. Khan, E. Graham, P. Critchley, A.E. Schrag, N.W. Wood, A.J. Lees, et al., Parkin disease: a phenotypic study of a large case series, Brain 126(Pt 6) (2003) 1279–92.

81. C. Klein, K. Lohmann, Parkinson disease(s): is “Parkin disease” a distinct clinical entity?, Neurology 72(2) (2009) 106–7.

82. S. Lesage, A. Lunati, M. Houot, S.B. Romdhan, F. Clot, C. Tesson, et al., Characterization of recessive Parkinson’s disease in a large multicenter study, Ann Neurol (2020).

83. N. Giguere, C. Pacelli, C. Saumure, M.J. Bourque, D. Matheoud, D. Levesque, et al., Comparative analysis of Parkinson’s disease-associated genes in mice reveals altered survival and bioenergetics of Parkin-deficient dopamine neurons, J Biol Chem 293(25) (2018) 9580–9593.

84. W. Dauer, S. Przedborski, Parkinson’s disease: mechanisms and models, Neuron 39(6) (2003) 889–909.

85. J.M. Flynn, S. Melov, SOD2 in mitochondrial dysfunction and neurodegeneration, Free Radic Biol Med 62 (2013) 4–12.

86. M.S. Goldberg, S.M. Fleming, J.J. Palacino, C. Cepeda, H.A. Lam, A. Bhatnagar, et al., Parkin-deficient mice exhibit nigrostriatal deficits but not loss of dopaminergic neurons, J Biol Chem 278(44) (2003) 43628–35.

87. J.A. Rodriguez-Navarro, M.J. Casarejos, J. Menendez, R.M. Solano, I. Rodal, A. Gomez, et al., Mortality, oxidative stress and tau accumulation during ageing in parkin null mice, J Neurochem 103(1) (2007) 98–114.

88. R.M. Solano, M.J. Casarejos, J. Menendez-Cuervo, J.A. Rodriguez-Navarro, J. Garcia de Yebenes, M.A. Mena, Glial dysfunction in parkin null mice: effects of aging, J Neurosci 28(3) (2008) 598–611.

89. M. Damiano, C.A. Gautier, A.L. Bulteau, R. Ferrando-Miguel, C. Gouarne, M.G. Paoli, et al., Tissue- and cell-specific mitochondrial defect in Parkin-deficient mice, PLoS One 9(6) (2014) e99898.

90. H. Jiang, Y. Ren, J. Zhao, J. Feng, Parkin protects human dopaminergic neuroblastoma cells against dopamine-induced apoptosis, Hum Mol Genet 13(16) (2004) 1745–54.

91. T. Kitada, A. Pisani, M. Karouani, M. Haburcak, G. Martella, A. Tscherter, et al., Impaired dopamine release and synaptic plasticity in the striatum of parkin−/− mice, J Neurochem 110(2) (2009) 613–21.

92. C. Gladkova, S.L. Maslen, J.M. Skehel, D. Komander, Mechanism of parkin activation by PINK1, Nature 559(7714) (2018) 410–414.

93. W.R. Gibb, H. Narabayashi, M. Yokochi, R. Iizuka, A.J. Lees, New pathologic observations in juvenile onset parkinsonism with dystonia, Neurology 41(6) (1991) 820–2.

94. H. Takahashi, E. Ohama, S. Suzuki, Y. Horikawa, A. Ishikawa, T. Morita, et al., Familial juvenile parkinsonism: clinical and pathologic study in a family, Neurology 44(3 Pt 1) (1994) 437–41.

95. Y. Yamamura, S. Kuzuhara, K. Kondo, T. Yanagi, M. Uchida, H. Matsumine, et al., Clinical, pathologic and genetic studies on autosomal recessive early-onset parkinsonism with diurnal fluctuation, Parkinsonism Relat Disord 4(2) (1998) 65–72.

96. S. Hayashi, K. Wakabayashi, A. Ishikawa, H. Nagai, M. Saito, M. Maruyama, et al., An autopsy case of autosomal-recessive juvenile parkinsonism with a homozygous exon 4 deletion in the parkin gene, Mov Disord 15(5) (2000) 884–8.

97. M. Yokochi, H. Narabayashi, R. Iizuka, T. Nagatsu, Juvenile parkinsonism--some clinical, pharmacological, and neuropathological aspects, Adv Neurol 40 (1984) 407–13.

98. N. Gouider-Khouja, A. Larnaout, R. Amouri, S. Sfar, S. Belal, C. Ben Hamida, et al., Autosomal recessive parkinsonism linked to parkin gene in a Tunisian family. Clinical, genetic and pathological study, Parkinsonism Relat Disord 9(5) (2003) 247–51.

99. C. International Parkinson Disease Genomics, M.A. Nalls, V. Plagnol, D.G. Hernandez, M. Sharma, U.M. Sheerin, et al., Imputation of sequence variants for identification of genetic risks for Parkinson’s disease: a meta-analysis of genome-wide association studies, Lancet 377(9766) (2011) 641–9.

100. Y.Q. Wang, B.S. Tang, R.L. Yu, K. Li, Z.H. Liu, Q. Xu, et al., Association analysis of STK39, MCCC1/LAMP3 and sporadic PD in the Chinese Han population, Neurosci Lett 566 (2014) 206–9.

101. M.A. Nalls, N. Pankratz, C.M. Lill, C.B. Do, D.G. Hernandez, M. Saad, et al., Large-scale meta-analysis of genome-wide association data identifies six new risk loci for Parkinson’s disease, Nat Genet 46(9) (2014) 989–93.

102. Y. Sun, M. Tan, H. Duan, M. Swaroop, SAG/ROC/Rbx/Hrt, a zinc RING finger gene family: molecular cloning, biochemical properties, and biological functions, Antioxid Redox Signal 3(4) (2001) 635–50.

103. Y. Sun, H. Li, Functional characterization of SAG/RBX2/ROC2/RNF7, an antioxidant protein and an E3 ubiquitin ligase, Protein Cell 4(2) (2013) 103–16.

104. K.M. Rosen, V. Veereshwarayya, C.E. Moussa, Q. Fu, M.S. Goldberg, M.G. Schlossmacher, et al., Parkin protects against mitochondrial toxins and beta-amyloid accumulation in skeletal muscle cells, J Biol Chem 281(18) (2006) 12809–16.

105. T.M. Dawson, V.L. Dawson, Parkin plays a role in sporadic Parkinson’s disease, Neurodegener Dis 13(2–3) (2014) 69–71.

106. T. Kitada, J.J. Tomlinson, H.S. Ao, D.A. Grimes, M.G. Schlossmacher, Considerations regarding the etiology and future treatment of autosomal recessive versus idiopathic Parkinson disease, Curr Treat Options Neurol 14(3) (2012) 230–40.

107. R.R. Ratan, Antioxidants and the Treatment of Neurological Disease, in: V.E. Koliatsos, Ratan, R. R. (Ed.), Cell Death and Diseases of the Nervous System, Humana Press, Totowa, NJ, 1999.

